# Characterizing genetic intra-tumor heterogeneity across 2,658 human cancer genomes

**DOI:** 10.1101/312041

**Authors:** Stefan C. Dentro, Ignaty Leshchiner, Kerstin Haase, Maxime Tarabichi, Jeff Wintersinger, Amit G. Deshwar, Kaixian Yu, Yulia Rubanova, Geoff Macintyre, Jonas Demeulemeester, Ignacio Vázquez-García, Kortine Kleinheinz, Dimitri G. Livitz, Salem Malikic, Nilgun Donmez, Subhajit Sengupta, Pavana Anur, Clemency Jolly, Marek Cmero, Daniel Rosebrock, Steven Schumacher, Yu Fan, Matthew Fittall, Ruben M. Drews, Xiaotong Yao, Juhee Lee, Matthias Schlesner, Hongtu Zhu, David J. Adams, Gad Getz, Paul C. Boutros, Marcin Imielinski, Rameen Beroukhim, S. Cenk Sahinalp, Yuan Ji, Martin Peifer, Inigo Martincorena, Florian Markowetz, Ville Mustonen, Ke Yuan, Moritz Gerstung, Paul T. Spellman, Wenyi Wang, Quaid D. Morris, David C. Wedge, Peter Van Loo, on behalf of the PCAWG Evolution and Heterogeneity Working Groupthe PCAWG consortium., the PCAWG consortium

**Affiliations:** The Francis Crick Institute, London, United Kingdom; Wellcome Trust Sanger Institute, Cambridge, United Kingdom; Big Data Institute, University of Oxford, Oxford, United Kingdom; Broad Institute of MIT and Harvard, Cambridge, MA, USA; University of Toronto, Toronto, Canada; The University of Texas MD Anderson Cancer Center, Houston, TX, USA; Cancer Research UK Cambridge Institute, University of Cambridge, Cambridge, United Kingdom; Department of Human Genetics, University of Leuven, Leuven, Belgium; University of Cambridge, Cambridge, United Kingdom; Computational Oncology, Memorial Sloan Kettering Cancer Center, New York, NY, USA; Irving Institute for Cancer Dynamics, Columbia University, New York, NY, USA; German Cancer Research Center (DKFZ), Heidelberg, Germany; Heidelberg University, Heidelberg, Germany; Simon Fraser University, Vancouver, Canada; NorthShore University HealthSystem, Evanston, IL, USA; Molecular and Medical Genetics, Oregon Health & Science University, Portland, OR, USA; University of Melbourne, Melbourne, Australia; Weill Cornell Medicine, New York, NY, USA; New York Genome Center, New York, NY, USA; University of California Santa Cruz, Santa Cruz, CA, USA; University of California, Los Angeles, Los Angeles, CA, USA; Indiana University, Bloomington, IN, USA; The University of Chicago, Chicago, IL, USA; University of Cologne, Cologne, Germany; Organismal and Evolutionary Biology Research Programme, Department of Computer Science, Institute of Biotechnology, University of Helsinki, Helsinki, Finland; School of Computing Science, University of Glasgow, Glasgow, United Kingdom; European Molecular Biology Laboratory, European Bioinformatics Institute, Cambridge, United Kingdom; Oxford NIHR Biomedical Research Centre, Oxford, United Kingdom

## Abstract

Intra-tumor heterogeneity (ITH) is a mechanism of therapeutic resistance and therefore an important clinical challenge. However, the extent, origin and drivers of ITH across cancer types are poorly understood. To address this question, we extensively characterize ITH across whole-genome sequences of 2,658 cancer samples, spanning 38 cancer types. Nearly all informative samples (95.1%) contain evidence of distinct subclonal expansions, with frequent branching relationships between subclones. We observe positive selection of subclonal driver mutations across most cancer types, and identify cancer type specific subclonal patterns of driver gene mutations, fusions, structural variants and copy-number alterations, as well as dynamic changes in mutational processes between subclonal expansions. Our results underline the importance of ITH and its drivers in tumor evolution, and provide an unprecedented pan-cancer resource of comprehensively annotated subclonal events from whole-genome sequencing data.

## INTRODUCTION

Cancers accumulate somatic mutations as they evolve (Nowell, 1976; Tabin et al., 1982). Some of these mutations are drivers that confer fitness advantages to their host cells and can lead to clonal expansions (Garraway and Lander, 2013; Greaves and Maley, 2012; Stratton et al., 2009; Vogelstein et al., 2013). Late clonal expansions, spatial segregation, and incomplete selective sweeps result in genetically distinct cellular populations that manifest as intra-tumor heterogeneity (ITH) (Nowell, 1976). Clonal mutations are shared by all cancer cells, whereas subclonal mutations are present only in a fraction of cancer cells.

ITH represents an important clinical challenge, as it provides genetic variation that may drive cancer progression and lead to the emergence of drug resistance (Maley et al., 2006; McGranahan and Swanton, 2017; Mroz et al., 2013). Subclonal drug resistance and associated driver mutations are common (Gerlinger et al., 2012; Gundem et al., 2015; Landau et al., 2013; McGranahan et al., 2015; Shaw et al., 2016; Yates et al., 2015). ITH can impact clinical trial design (Hiley et al., 2014), predict progression (Maley et al., 2004), and can be directly prognostic (Espiritu et al., 2018). For example, ITH at the level of copy number aberrations (CNAs) is associated with increased risk of relapse in non-small cell lung cancer (Jamal-Hanjani et al., 2017), head and neck cancer (Mroz and Rocco, 2013; Rocco, 2015) and glioblastoma multiforme (Brastianos et al., 2017).

ITH can be characterized from massively parallel sequencing data (Campbell et al., 2008; Landau et al., 2013; McGranahan et al., 2015; Nik-Zainal et al., 2012; Sottoriva et al., 2013), as the cells comprising a clonal expansion share a unique set of driver and passenger mutations derived from the expansion-initiating cell. Each mutation within this shared set is present in the same proportion of tumor cells (known as cancer cell fraction, CCF), which may be estimated by adjusting mutation allele frequencies for local copy number and sample purity. Subsequent clustering of mutations based on their CCF yields the ‘subclonal architecture’ of a sample (Dentro et al., 2017): estimates of the number of tumor cell populations in the sequenced sample, the CCF of each population, and assignments of mutations to each population.

To date, ITH remains poorly characterized across cancer types, and there is substantial uncertainty concerning the selective pressures operating on subclonal populations. Previous pan-cancer efforts used the principles above to characterize subclonal events, but have been limited to exomes, which restricts the number and resolution of somatic mutation calls and ignores structural variation (Andor et al., 2016). Two recent studies using pan-]cancer data from The Cancer Genome Atlas found that actionable driver mutations are often subclonal (McGranahan et al., 2015), and that ITH has broad prognostic value (Andor et al., 2016).

Recent studies have relied on multi-region whole-genome, exome or targeted sequencing to characterize ITH in detail in specific cancer types (Jamal-Hanjani et al., 2017; McPherson et al., 2016; Turajlic et al., 2018b; Yates et al., 2015). Due to the ‘illusion of clonality’ (de Bruin et al., 2014), variants found as clonal in one sample may be subclonal in other samples from the same tumor, and therefore single-sample analyses can underestimate the extent of ITH. The converse is also true: any mutations detected as subclonal in any single sample, will by definition be subclonal no matter how many samples have been assayed. Therefore, through analyzing single cancer samples, a conservative lower limit of ITH can be established.

Here, we develop a robust consensus strategy that maintains conservative inferences to call copy number and cluster mutations in order to assess ITH, its origin, its drivers, and its role in tumor development. We apply these approaches to 2,658 tumors from 38 histologically distinct cancer types from the Pan-Cancer Analysis of Whole Genomes (PCAWG) initiative (The ICGC/TCGA Pan-Cancer Analysis of Whole Genomes Consortium, 2020). In comparison to exome sequencing, whole-genome sequencing data provides orders of magnitude more point mutations, greater resolution to detect CNAs and the ability to call structural variants (SVs). Collectively, these substantially increase the breadth and depth of our ITH analyses permitting us to find pervasive ITH across all cancer types. We are further able to observe frequent branching patterns of subclonal evolution and clear signs of positive selection in subclones. We identify subclonal driver mutations in known cancer genes and unanticipated changes in mutation signature activity across many cancer types. In total, these analyses provide detailed insight into tumor evolutionary dynamics.

## RESULTS

### Consensus-based characterization of intra-tumor heterogeneity in 2,658 cancers

We set out to characterize ITH across cancer types, including single-nucleotide variants (SNVs), indels, SVs and CNAs, as well as subclonal drivers, subclonal selection, and mutation signatures. We leveraged the PCAWG dataset, encompassing 2,778 whole-]genome sequences from 2,658 human tumors across 38 distinct histological cancer types (Alexandrov et al., 2020; Gerstung et al., 2020; Rheinbay et al., 2020; The ICGC/TCGA Pan-Cancer Analysis of Whole Genomes Consortium, 2020).

First, to generate high-confidence calls, we developed ensemble approaches for variant calling, copy number calling and subclonal reconstruction (**Figure 1A, STAR Methods**). Specifically, to maximize sensitivity and specificity of calling clonal and subclonal mutations, the PCAWG consortium developed and extensively validated a robust consensus approach integrating the output of four SNV calling algorithms (The ICGC/TCGA Pan-Cancer Analysis of Whole Genomes Consortium, 2020). Similar consensus approaches were employed for indels and SVs.

**Figure 1.**
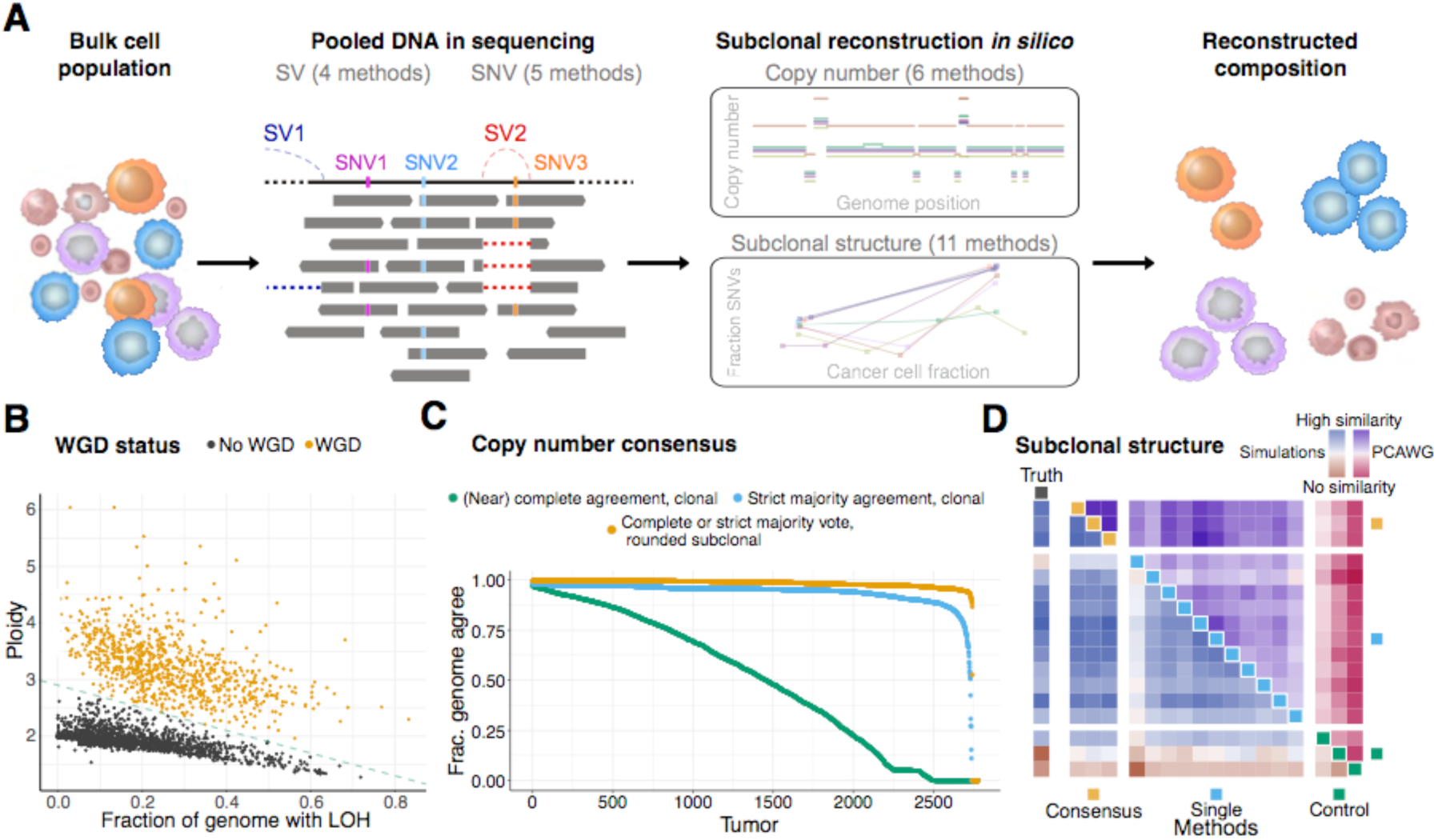
Consensus-based characterization of intra-tumor heterogeneity. (A) Schematic representation of our consensus-based intra-tumor heterogeneity (ITH) reconstruction from sequencing data. (B) Samples with and without whole-genome duplications separate in two clusters according to their consensus ploidy and the fraction of the genome showing loss of heterozygosity. (C) Agreement between the six copy number callers using a multi-tier consensus copy number calling approach. The three lines denote the fraction of the genome at which agreement is reached at different levels of confidence: (near-)complete agreement on both alleles of clonal copy number, a strict majority agreement on both alleles of clonal copy number and (near-)complete or strict majority agreement on both alleles of rounded subclonal copy number (see **STAR Methods**). At the third level, agreement is reached on an average 93% of the genome. (D) Heatmap of the normalized average pairwise similarities of subclonal architectures identified by 11 individual, 3 consensus, and 3 control reconstruction methods. Each method is represented by one colored square on the diagonal. On rows and columns, each method is compared to all other methods. The upper triangle shows the average pairwise similarities on the 2,778 PCAWG samples, the lower triangle shows the same on a validation set of 965 simulated samples. In the leftmost column similarities are computed against the truth of the simulated set. Color intensities scale with the similarities and were normalized separately for PCAWG, simulations and truth.

As previous studies report that the quality of copy number calls has a large effect on the robustness of subclonal reconstruction (Andor et al., 2016; Salcedo et al., 2020), we devised a systematic approach to consensus copy number calling, integrating results from six state-of-the-art copy number callers (**Figure 1A, STAR Methods**). Each algorithm was run twice, first to identify all copy number breakpoints and construct a consensus segmentation. To improve sensitivity and obtain breakpoints at base-pair resolution, SV breakpoints were also inserted into these first runs (**STAR Methods**). In a second run, this consensus segmentation was enforced on all CNA callers, resulting in copy number calls with identical breakpoints across algorithms.

Purity and ploidy assessment of cancer samples can be challenging, as for some samples multiple purity/ploidy combinations can be theoretically possible and these may be difficult to distinguish (Carter et al., 2012; Van Loo et al., 2010). Consensus purity and ploidy were determined by establishing agreement between the six CNA callers (**STAR Methods**). An expert panel reviewed and resolved cases where the callers disagreed. We found that the purity values correlate strongly with a recent cross-omics analysis of tumor purity (Aran et al., 2015) (**Figure S1**). After establishing agreement on the purity and ploidy, we defined samples that had undergone whole-genome duplication in an objective and automated way, based on tumor ploidy and the extent of loss of heterozygosity (**Figure 1B, STAR Methods**). This classification shows 98.7% agreement with an alternative approach leveraging the mode of the major allele (Carter et al., 2012). However, our classification correctly classifies difficult tumors with many large chromosome gains, such as medulloblastomas or pancreatic endocrine tumors, whereas the alternative approach is less suitable for these tumors and occasionally makes errors. In addition, samples with whole-]genome duplications showed synchronous chromosomal gains (Gerstung et al., 2020), further validating our approach. To further support high-quality subclonal reconstruction, the whole genome of each tumor was annotated for the confidence in the consensus copy number calls, which were assigned ‘tiers’ based on the level of agreement between different callers. On average, we reached a high confidence consensus on 93% of the genome (median: 95%, standard deviation: 13%) (**Figure 1C, STAR Methods**).

Consensus copy number profiles, SNVs, and purity estimates served as input to 11 subclonal architecture reconstructing methods, and the results of these methods were combined into a single consensus reconstruction for each tumor (**Figure 1A, STAR Methods**). Due to the probabilistic nature of subclonal reconstruction, we developed three consensus approaches using different summary outputs of individual methods. We validated the results of the consensus strategies on two independently simulated datasets and assessed their robustness on the real data. The consensus methods performed comparably to the best individual methods on both simulated datasets, with the top-]performing individual methods also displaying high similarity scores (**Figure 1D, STAR Methods**). Whereas true mutations and CNAs were used in the analysis of simulated data, in the real data the true subclonal mutations and CNAs are unknown. On the real data, the highest similarities were observed for the consensus approaches, and not among individual methods (**Figure 1D**), confirming that our consensus approaches yield the most robust subclonal reconstruction outcome. Furthermore, using one simulated dataset with 965 samples, we evaluated the performances of our consensus methods over all 2,035 possible combinations of 11 individual methods. We observed that the most robust performance, when the best callers are not known *a priori*, was achieved when all 11 callers were combined (**STAR Methods**). Hence, we used the output of one of our consensus methods, combining all 11 individual callers, as the basis for our global assignment strategy (**STAR Methods**). Through this approach, we obtained the number of detectable subclonal expansions, the fraction of subclonal SNVs, indels, SVs and CNAs, as well as the assignment of SNVs, indels and SVs to subclones for each tumor.

To obtain unbiased estimates of the number of mutations in the detected subclones and their CCFs, we accounted for a detection bias introduced by somatic variant calling. Specifically, as the CCF of a subclone decreases, so does the power to detect the SNVs associated with that subclone. This leads to biases in the estimates of subclone parameters, such as an overestimation of the subclone’s CCF, akin to the “winner’s curse” (Nik-Zainal et al., 2012). In addition, an increasing number of uncalled SNVs in the subclone leads to an underestimation of the number of associated mutations. The larger number of SNVs revealed by WGS (compared to whole-exome sequencing) facilitates quantitation and correction of these biases. We developed two methods to do this, validated them on simulated data (**STAR Methods, Figure 2A**), and combined them to correct the estimated number of SNVs and the CCF of each subclone. We estimate that, on average, 14% of SNVs in detectable subclones are below the somatic caller detection limits (**Figures 2B-C**). In particular, in subclones with CCF < 30%, on average 21% of SNVs are missed. Due to the complexity in modelling sensitivity of indel and SV calling as a function of the number of variant reads, similar models could not be developed for these mutation types. However, we anticipate that higher fractions of SVs and indels are likely missed because of the lower sensitivity of existing algorithms (The ICGC/TCGA Pan-Cancer Analysis of Whole Genomes Consortium, 2020). In addition, these values only include SNVs missed in detected subclones, not SNVs in subclones that remain undetected due to limited sequencing depth.

**Figure 2.**
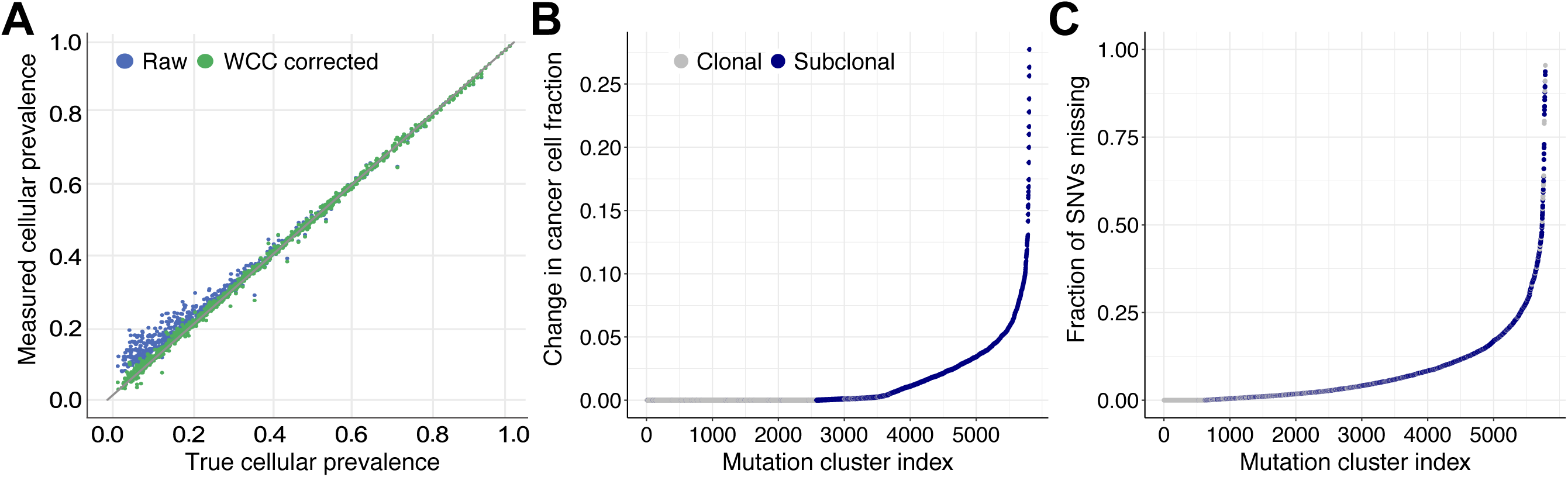
Winner’s curse correction. (A) Validation of our approach to adjust for the “winner’s curse-like effect, and (B-C) the estimated cluster-CCF and mutation adjustment in all mutation clusters identified in the study. Subclonal clusters show a shift to larger CCF values after correction (B) and the majority of clusters are estimated to contain additional missed SNVs (C).

### Pervasive intra-tumor heterogeneity across cancer types

We first evaluated the number of subclones identified by our consensus approach. We noted a strong correlation between the average effective read depth along the genome (sequencing coverage per haploid genome copy, i.e. the amount of sequencing signal) and the number of identified subclones (**Figure S2**), and therefore focused on 1,705 cancer genomes where our approach is powered to detect subclones encompassing >30% of tumor cells (**STAR Methods**). One or more subclonal expansions were evident in 1,621 tumors (95.1%), while only 84 tumors (4.9%) were clonal at the resolution of our methods (**Figure 3A**). Importantly, these estimates, based on single-sample reconstruction and a median ∼46X read coverage, provide only a conservative lower bound for the number of subclones, as this study is not powered to detect rare subclonal populations.

**Figure 3.**
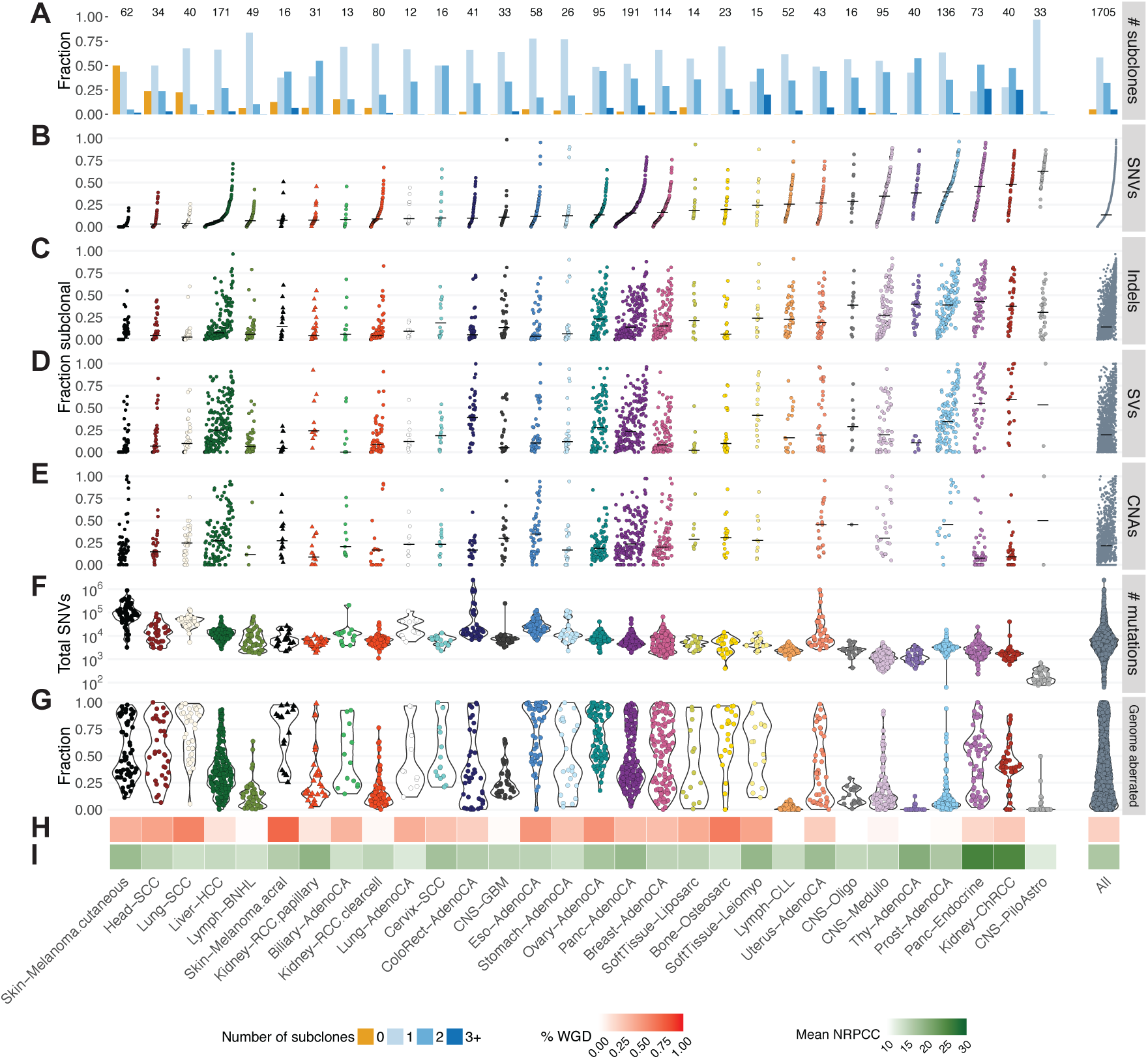
Overview and characterization of ITH across cancer types. Evidence of ITH is shown for 1,705 samples with sufficient power to detect subclones at CCF > 30% (see **STAR Methods**). Samples have been limited to those with less than 2% tumor contamination in the matched normal sample and no activity of any of the identified artefact signatures (Alexandrov et al., 2020). Only representative samples (The ICGC/TCGA Pan-Cancer Analysis of Whole Genomes Consortium, 2020) from multi-]sample cases are shown. (A) Bar plot showing the fraction of samples with given number of subclones; (B-E) Scatter plots showing the fractions of subclonal SNVs, indels, SVs and subclonal arm-level CNAs (the latter two mutation types are only plotted for samples that have at least 5 events, sample order is determined by increasing fraction of subclonal SNVs and conserved in the other three panels); Violin plots showing the total mutation burden (F) and overall fraction of the genome that does not have a copy number state of 1+1, or 2+2 in WGD samples (G); Heatmaps showing the fraction of tumor samples with whole genome duplications (H) and the mean power to identify subclones per cancer types (number of reads per clonal copy – nrpcc, see **STAR Methods)** (I).

Looking across cancer types (**Figure 3A**), our consensus approach finds a high proportion of samples with at least one subclone (>75%) in all cancer types, except cutaneous melanoma, where subclones were detectable by our approach in only half of the samples (31/62). In contrast, acral melanomas followed the pattern observed in other cancer types with higher frequencies of subclonal expansions (14/16 samples, 87.5%). Twenty-five out of 30 cancer types with more than 10 cases comprised >90% of samples with at least one detectable subclone, indicating pervasive ITH across cancer types.

The fraction of subclonal SNVs identified after winner’s curse correction varies widely across cancer types (**Figure 3B**). Squamous cell carcinomas typically show low fractions of subclonal SNVs (head-and-neck, 9.7% ± 11.9; lung, 6.1% ± 6.7; cervix, 20.6% ± 19.6; mean ± standard deviation), while prostate adenocarcinoma (41.2% ± 21.8), thyroid adenocarcinoma (42.8% ± 19.6), chromophobe renal cell carcinomas (45.2% ± 22.7), pancreatic neuroendocrine tumors (46% ± 24.7) and pilocytic astrocytomas (61.3% ± 14.8) showed the highest fractions of subclonal SNVs.

Indels, SVs and CNAs also revealed similarly large differences between cancer types. For indels, the subclonal fraction ranged from 6.2% ± 11.7 in lung squamous carcinomas to 43.4% ± 23.7 in pancreatic neuroendocrine tumors (**Figure 3C**). For SVs, liposarcomas and cutaneous melanomas showed the lowest subclonal fraction (8.0% ± 14.1 and 8.9% ± 15.5 respectively) and chromophobe renal cell cancers the highest (56.8% ± 31.6) (**Figure 3D**). The fraction of subclonal copy number changes was lowest in chromophobe renal cell cancers (13.3% ± 16.8) and highest in prostate adenocarcinoma (53.8% ± 32.0) (**Figure 3E**). Comparing these values to the SNV burden (**Figure 3F**), the fraction of the genome affected by CNAs (**Figure 3G**), the frequency of WGD per cancer type (**Figure 3H**), and the power to identify subclones per cancer type (**Figure 3I**) showed that none of these metrics explain this wide variation. While we observed that cancer types with higher mutation burden showed lower fractions of subclonal SNVs (**Figures 3B and 3F**), we did not see a similar relationship when evaluating individual tumors (**STAR Methods**). The proportions of subclonal indels and SNVs are strongly correlated (*R*^*2*^ = 0.73). SVs follow a similar trend (*R*^*2*^ = 0.62 with indels, *R*^*2*^ = 0.51 with SNVs), except for liver, colorectal and ovarian tumors, which show higher fractions of subclonal SVs than SNVs (**Figures 3B-E and S3**). In contrast, the average proportions of subclonal large-scale CNAs and SNVs are only weakly correlated (*R*^2^ = 0.24), indicating these could be driven by independent mutational processes.

Some cancer types had limited ITH across all mutation types (e.g. biliary cancers, squamous cell carcinomas and stomach cancers), while other cancer types showed an abundance of ITH in specific somatic variant categories. For example, chromophobe kidney cancers and pancreatic neuroendocrine tumors have few subclonal CNAs but a high subclonal burden across all other variant categories (**Figures 3B-E**). Finally, among the tumors of each cancer type, we find substantial diversity in the fraction of subclonal variants (**Figures 3B-E**).

These findings highlight the high prevalence of ITH across cancer types. Nearly all tumors assayed here, irrespective of cancer type, show evidence of subclonal expansions giving rise to detectable subclonal populations, even at a limited read depth. In addition, we find that the average proportions of subclonal SNVs, indels, SVs and CNAs are highly variable across cancer types. These analyses paint characteristic portraits of the nature of ITH, suggesting distinct evolutionary narratives for each histological cancer type.

### Complex phylogenies among subclones revealed by whole genome sequencing

Whole-genome sequencing provides an opportunity to explore and reconstruct additional patterns of subclonal structure by performing phasing of pairs of mutations in the same read pair, to assess evolutionary relationships of subclonal lineages (**Figures 4A-B**). Two subclones can be either linearly related to each other (parent-child relationship), or have a common ancestor, but develop on branching lineages (sibling subclones). Establishing evolutionary relationships between subclones is challenging on single-sample sequencing data due to the limited resolution to separate subclones and the uncertainties on their CCF estimates. We can, however, examine pairs of SNVs in WGS data that are covered by the same read pairs (*i.e.* phaseable SNV pairs), to reconstruct this relationship. Specifically, evidence for a parent-child relationship between two clones is given by a SNV pattern in a region without copy number gains, where the SNV attributed to the child clone is only found on a subset of the reads that carry the SNV attributed to the parent clone (**Figure 4A, STAR Methods**). Similarly, evidence for a sibling relationship between two clones is given by an SNV pattern in a haploid region where overlapping read pairs carry either the SNV attributed to one clone or the other (but not both) (**Figure 4B, STAR methods**). As the number of read pairs carrying two variants depends strongly on mutation burden and specific copy number context, they are generally extremely rare. Our large, curated dataset, however, enables us to identify a sizeable total of these and explore their phylogenetic information content in detail.

**Figure 4.**
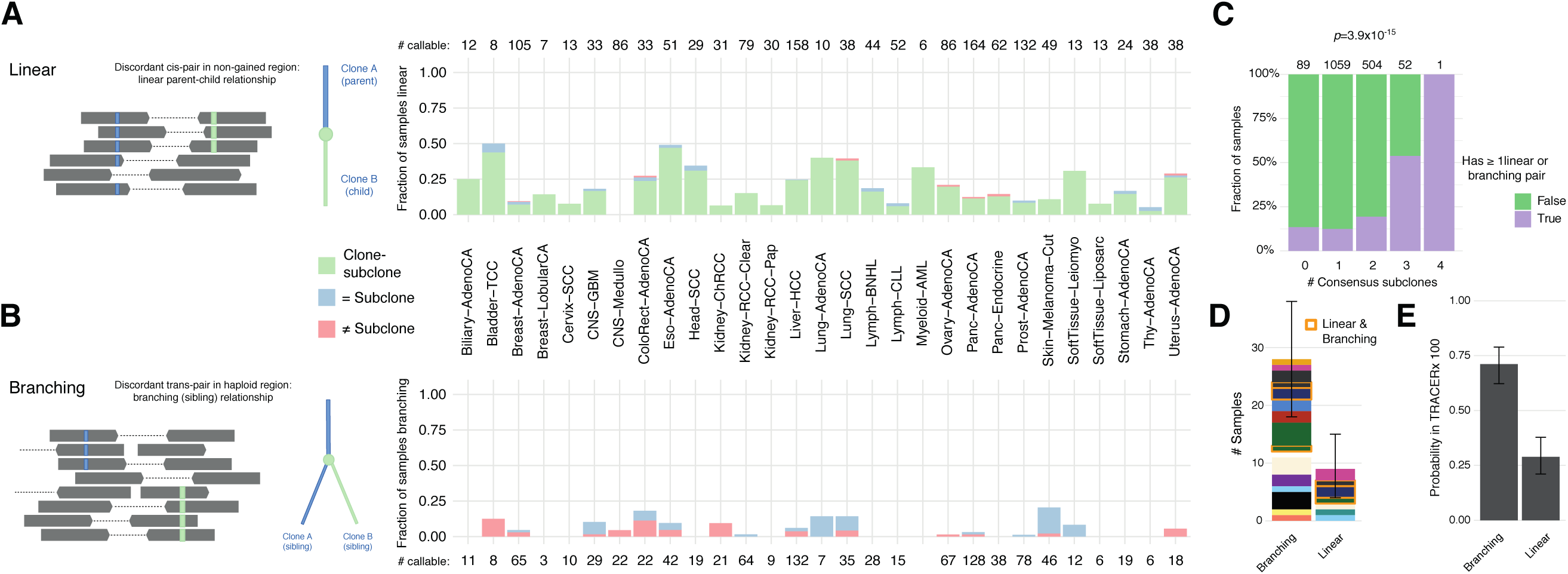
Further characterization of ITH using mutation phasing. (A-B) Proportion of powered tumors with evidence of linear and branching phylogenies, through analysis of phased reads of variants in-*cis* (A) or in-*trans* (B) among tumors with at least one phaseable pair in the appropriate context. (C) Fraction of powered samples, stratified by number of consensus subclones, with at least one linear or branching pair (*χ*^2^-test for independence). (D) Number of samples with linear or branching pairs when sets are filtered to be comparable. Error bars indicate the 95% bootstrap interval. Samples are colored by tumor type and boxed (orange) when they present with pairs of both types. (E) Probabilities of observing a linear *vs.* branching relationship when picking two random subclones from TRACERx 100 trees (Jamal-Hanjani et al., 2017). Error bars indicate the 95% bootstrap interval.

We find that of 1,537 tumors with sufficient power and at least one phaseable pair in the correct context, 245 show discordant in-*cis* SNVs pairs, indicating parent-child relationships (**Figure 4A**). Annotating SNVs with their clone or subclone assignment from CCF clustering, the vast majority of these samples (233, 95.1%) show pairs supporting the expected clone-subclone relationship. In addition, there are 8 samples with pairs assigned to different subclones and 16 samples with pairs assigned to the same subclone, highlighting collinear evolution among subclones. Of 995 tumors, 51 carry discordant in-]*trans* SNVs pairs, with 32 and 27 of these samples having pairs assigned to the same or different subclones, respectively (8 have both), confirming the occurrence of two sibling subclones having expanded in parallel (**Figure 4B**).

The identification of subclones from phaseable SNV pairs can be considered largely independent of the consensus subclonal reconstruction. One can therefore use phasing results to assess the performance of subclonal reconstruction and *vice versa*. Indeed, tumors identified to contain higher numbers of subclones according to the consensus reconstruction are enriched for linear and branching pairs (p-value = 3.9×10^−15^, **Figure 4C**). Nevertheless, our identification of 16 and 32 samples with SNV pairs assigned to the same subclone but showing in-*cis* and in-*trans* discordance, respectively, confirms that our consensus approach identifies only a lower limit on the number of subclonal expansions. Interestingly, 8 of the 13 cutaneous melanomas showing linear or branching pairs had been deemed clonal by CCF clustering but had phasing evidence for 1–2 subclones (3 had linear, 7 had branching, and 3 had both linear and branching SNV pairs). This analysis suggests that, similarly to other cancer types, the large majority of cutaneous melanomas contain subclonal expansions. However, these might be obscured by the large numbers of clonal mutations in these extremely highly mutated tumors.

The frequency of branching versus linear evolution can be assessed directly by subsetting the phasing analysis to haploid regions and to pairs where both SNVs have been assigned to subclones, as both linear and branching relationships may be detected with similar power in this subset. Our results indicate that, in the pan-cancer setting, two subclones are 3.11 times more likely to be siblings than to have a parent-child relationship (bootstrapped 95% confidence interval [1.71; 7.50], **Figure 4D**). This result is consistent with the complex phylogenies obtained from multi-region sequencing efforts such as the TRACERx 100 non-small-cell lung cancer cohort (Jamal-Hanjani et al., 2017), where the odds of branching *vs.* linear evolution are 2.86 (bootstrapped 95% confidence interval [1.93; 5.07], **Figure 4E, STAR Methods)**. These results are also in line with observations of mutual exclusivity of subclonal drivers and extensive parallel evolution (Turajlic et al., 2018a; Turajlic et al., 2018b).

### Patterns of subclonal mutation signature activity changes across cancers

Mutation processes can differ in their activity between clonal and subclonal lineages (McGranahan et al., 2015). To explore the subclonal dynamics of mutation signatures in detail, we examined subclonal mutations for changes in signature activity. We reasoned that if a mutation process is activated during a specific subclonal expansion, only the post-]expansion mutations will carry the corresponding mutation signature. Signature activity change points can therefore be identified in SNVs that are rank-ordered by their CCFs estimates (Rubanova et al., 2020) (**STAR Methods**). Of the 2,552 samples with sufficient SNVs to perform this analysis, 1,944 (76%) have an activity change of at least 5% in at least one signature (a conservative threshold established *via* permutation and bootstrapping analyses, **STAR Methods**). We detect an average of 1.77 mutation signature activity clusters per sample.

Overall, mutation signature activity is remarkably stable. The most frequently changing signature (Signature SBS12, etiology unknown (Alexandrov et al., 2020), active in 198 of 326 (61%) liver cancers), is variable in approximately 60% of the cases in which it is active (**Figure 5A**). In addition, we find that the activity of Signature SBS9 (Pol η activity on AID lesions) decreases as a function of decreasing CCF in over half the tumors in which this signature is active (CLL and B-cell non-Hodgkin lymphoma). When only considering pairs of signatures that change in the same tumor, we see that 6 out of the top 10 pairs involve SBS5 (etiology unknown but hypothesized to reflect lower-fidelity DNA repair pathways (Kim et al., 2016)). Such changes in proportions are often anti-correlated, as the activity of one mutation process may be changing at the proportional expense of the activity of another.

**Figure 5.**
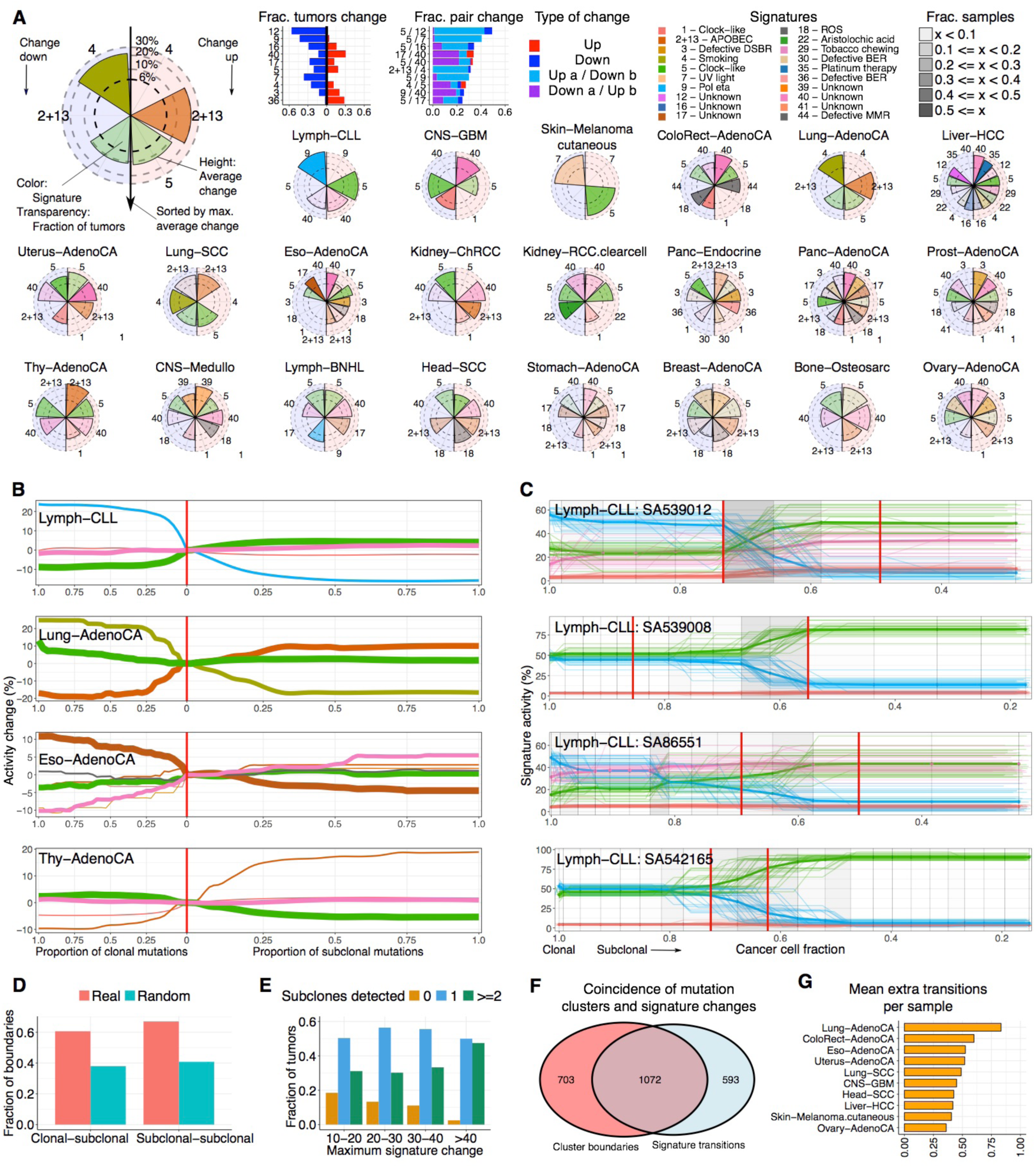
Subclonal boundaries are associated with changes in mutation signature activity. (A) Mutation signature changes across cancer types. Bar graphs show the proportion of tumors in which signature (pairs) change and radial plots provide a view per cancer type. Each radial plot contains the signatures that are active in at least 5 tumors and change (≥ 6%) in at least 3 tumors. The left and right side of the radial plot represent signatures that become less and more active, respectively. The height of a wedge represents the average activity change (log scale), the color denotes the signature and the transparency shows the fraction of tumors in which the signature changes (as a proportion of the tumors in which the signature is active). Signatures are sorted around the radial plot (top-to-bottom) by maximum average activity change. (B) Average signature trajectories for selected cancer types. Each line is colored by signature and corresponds to the average activity across tumors of this cancer type in which the signature is active. The width of the line represents the number of tumors that are represented. Mutations are split into clonal and subclonal, visually divided by a red vertical line. (C) Signature trajectories for selected individual CLL tumors. Each line corresponds to an activity trajectory derived from a bootstrap sample of SNVs. The grey vertical grid represents the mutation bins. These are colored grey when a significant change in signature activity is detected. Red vertical lines represent consensus subclonal mutation clusters. (D) The fraction of signature change points that coincide with boundaries between mutation clusters, as compared to what is expected when randomly placing change points. (E) The number of subclones detected in tumors grouped by the maximum detected signature activity change. (F) An overview of coinciding SNV cluster boundaries and signature activity change points. (G) The average number of additional signature change points detected per tumor.

We next evaluated signature trajectories per cancer type (**Figure 5A**). In CLL, SBS9 always decreases and SBS5 nearly always increases. In contrast, in ovarian cancers, most signature activity changes go both up and down in similar, relatively low proportions of tumors. On average, signature activity changes are modest in size, with the maximum average activity change recorded in CLL (33%, SBS9). Some changes are observed across many cancer types - *e.g.*, SBS5 and SBS40, of unknown etiology - while others are found in only one or a few cancer types. For example, in hepatocellular carcinomas, we observe an increase in SBS35 and a decrease in SBS12 (both etiology unknown), and in esophageal adenocarcinomas, we see an increase in SBS3 (double-strand break-repair) and a decrease in SBS17 (etiology unknown).

The average signature activity change across cancers of the same type is most often monotonic as a function of CCF. In other words, the activities of mutation processes consistently either decrease or increase (**Figures 5B and S4**). CLLs and lung adenocarcinomas initially exhibit a sharp change in signature activity when transitioning from clonal to subclonal mutations, but activity of the signatures appears to remain stable across multiple subclonal expansions (**Figure 5B**). In contrast, esophageal adenocarcinomas show a steady decrease in SBS17 activity, while thyroid adenocarcinomas often display a continuing increase in SBS2 and SBS13 (APOBEC) activity. These patterns observed across samples are also consistent at the single-sample level, for example in individual CLL samples (**Figure 5C**).

Interestingly, the SBS9 activity changes in CLL and B-cell non-Hodgkin lymphoma reflect the anatomical journey B cells have undergone in their evolution to cancers. Tumors showing SBS9 (pol η associated with AID activity) activity originate from post-germinal center B cells (Seifert et al., 2012). In these cases, SBS9 contributes to clonal but not subclonal mutations, because only the tumor-founding cell was exposed to somatic hypermutation in the germinal center. Later cells in this lineage have left the germinal center and are no longer exposed to AID. Similarly, the strong decrease of SBS7 (UV light) activity in cutaneous melanoma cases suggests these tumors have progressed to invade inner layers of the skin (Breslow, 1970), out of reach of damaging UVB exposure (Dupont et al., 2013). Finally, the co-occurring decrease of SBS4 (smoking) and increase of SBS2/13 (APOBEC) activity suggests that in lung cancers, cell-intrinsic mutation processes take over after early tumor evolution is fueled by external mutagens (Jamal-]Hanjani et al., 2017).

### Mutation signature activity changes mark subclonal boundaries

We next compared the mutation signature change points (shifts in activity) with the CCF of detected subclones, reasoning that these would correspond well if the emergence of subclones is associated with changes in mutation process activity. In such a scenario, we expect that the signature change points coincide with the CCF boundaries between subclones, assuming that clustering partitioned the SNVs accurately. In accordance with previous studies that highlight changes in signature activity between clonal and subclonal mutations (Jamal-Hanjani et al., 2017; McGranahan et al., 2015), we find that 34.5-54.7% of clone–subclone boundaries and 34.5%-57.3% of subclone-subclone boundaries coincide with a signature change point (**Figure 5D, STAR Methods**). This not only validates our clustering approach, but also demonstrates that subclonal expansions are often associated with changes in signature activity. It further suggests that increased ITH would correspond to greater activity change. Indeed, the samples with the largest changes in activity tend to be the most heterogeneous (**Figure 5E**). Conversely, an average of 0.49 changes per sample are not within a window of subclonal boundaries (**Figures 5F-G**), suggesting that some detected CCF clusters represent multiple subclonal lineages (**STAR Methods**), consistent with our mutation phasing results above.

### The landscape of subclonal driver mutations

We leveraged the comprehensive whole-genome view of driver events across these cancer genomes (Rheinbay et al., 2020) to gain insight into clonal *vs.* subclonal driver SNVs, indels and SVs. Out of 5,414 high-confidence SNV and indel driver mutations in 389 genes, we found 385 (7.1%) subclonal driver mutations across 147 distinct genes (**Figure 6A**). In total, 86% of samples with at least one subclone (1,576/1,831) contain no identified subclonal driver SNVs or indels, and only 11% of all detected subclones (280/2,542) were associated with acquisition of a clear subclonal driver SNV or indel. In contrast, clonal driver SNVs or indels were detected in 77% of samples (1,812/2,367).

**Figure 6.**
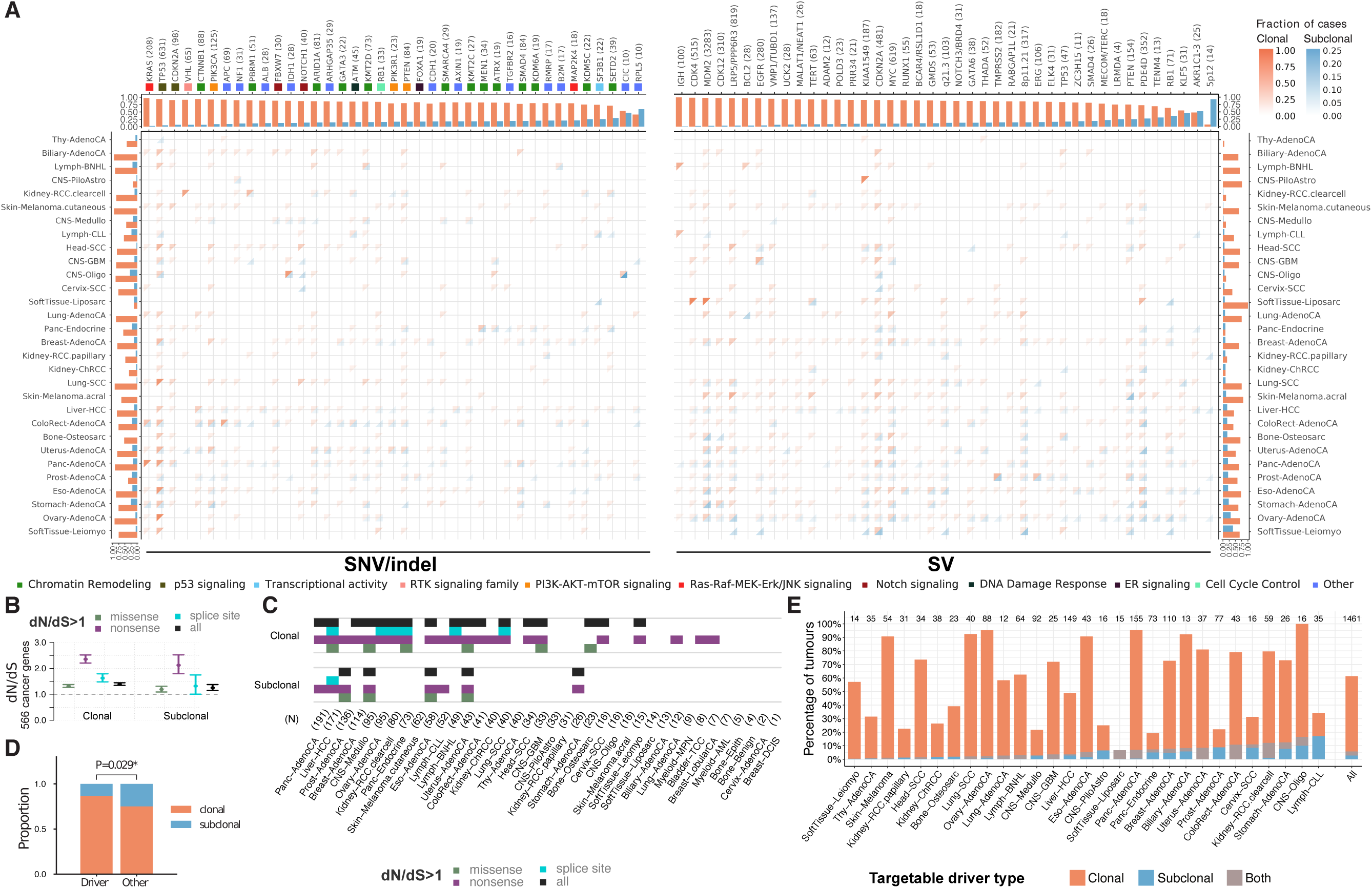
Driver mutations and subclonal selection. (A) Heatmap of the fraction of samples of the different cancer types with clonal (orange) and subclonal (blue) driver substitutions and indels (left panel) and structural variants (right panel). Marginal bar plots represent the fraction of clonal and subclonal driver mutations in each cancer type (side) and each driver gene or candidate region (top). Only genes with at least 4 subclonal driver mutations are shown. For SNVs and indel drivers (top left panel), gene set and pathway annotations highlight an enrichment of subclonally mutated drivers in chromatin remodeling. (B) dN/dS values for clonal and subclonal SNVs in 566 established cancer genes across all primary tumors. Values for missense, nonsense, splice site, and all mutations are shown, along with the 95% confidence intervals. (C) Cancer and mutation types for which dN/dS is significantly greater than 1 (95% confidence intervals>1) for clonal and subclonal mutations. Cancer types are ordered by the total number of samples. (D) Proportions of (sub)clonal driver gene fusions versus non-driver fusions. (E) Survey of targetable driver mutations across cancer types, stratified by clonal status.

As our whole-genome sequencing approach also allowed us to assess the clonality of SVs (Cmero et al., 2020), we next sought to examine the clonality of SV drivers. We considered an SV to be a driver if it was associated with a region of significantly recurrent breakpoints (Rheinbay et al., 2020) at non-fragile sites. By this analysis, 56.9% of samples analyzed (825/1,450) have a clonal SV driver, 14.7% (213/1,450) have at least one subclonal SV driver (**Figure 6A**), and 6.1% (89/1,450) have exclusively subclonal SV drivers. Pilocytic astrocytomas, non-Hodgkin’s lymphomas, biliary adenocarcinomas, and thyroid adenocarcinomas showed no evidence of subclonal SV drivers, while the remaining 26 of 30 cancer types analyzed all contained at least one subclonal SV driver in this cohort.

One explanation for the relative dearth of subclonal driver mutations is that subclonal driver mutations have a lower population prevalence than clonal ones. Specifically, driver identification depends on prevalence of the mutation within the cancer cohort. Our previous analysis demonstrated that the most prevalent drivers are also those that occur earliest in tumor development (Gerstung et al., 2020). This suggests that methods to annotate mutations (or genes) as drivers would be particularly prone to missing subclonal driver mutations. As such, we adapted a strategy from population genetics, to assess whether there was subclonal selection, even in the absence of discernible subclonal drivers. Selective pressures acting on the coding regions of cancer genomes can be quantified using the dN/dS ratio, which compares the rates of non-synonymous and synonymous mutations (Martincorena et al., 2017). A dN/dS ratio larger than 1 indicates positive selection, while smaller ratios characterize negative selection, and dN/dS ≈1 points towards neutral evolutionary dynamics. Previously, dN/dS > 1, evidence of positive selection, has been shown for cancer driver genes in all somatic mutations (Martincorena et al., 2017). When analyzing clonal mutations separately in our dataset, we confirm this signature of selection within a set of 566 well-established driver genes (**STAR Methods**). When specifically assaying our consensus subclonal mutations for the same set of drivers, we observe dN/dS > 1 for nonsense, missense and splice-site SNVs (**Figure 6B**). This indicates that selection for driver mutations, rather than neutral evolutionary dynamics (Williams et al., 2016), frequently shapes subclonal expansions, in agreement with our earlier study (Tarabichi et al., 2018). However, when considering dN/dS ratios for individual cancer types, we observe that in only a subset, the 95% confidence intervals exceed the threshold of positive selection (**Figure 6C**). The cancer types with evidence for selection had a significantly higher number of tumors sequenced (P = 1.6×10^−3^, Mann-Whitney U test), suggesting that the absence of conclusive signal in the remaining cancer types may be due to statistical power limitations.

The driver SNV and indel landscape indicates that specific genes recurrently harbor subclonal driver mutations across cancer types (**Figure 6A**). For example, the *SETD2* tumor suppressor is frequently subclonally mutated in clear cell renal cell carcinomas, as previously observed in multi-region sequencing experiments (Gerlinger et al., 2012), and in pancreatic neuroendocrine cancers. Interestingly, mutations in some driver genes that are exclusively clonal in most cancer types, are observed subclonally in others. For example, we find subclonal driver mutations in *MEN1* in pancreatic neuroendocrine tumors (6/30); *TP53* in prostate and breast cancers (4/12 and 5/59 respectively); and *CDKN2A* in pancreatic adenocarcinomas (5/42). Gene set analysis (**STAR Methods**) revealed an enrichment of subclonal mutations in genes responsible for chromatin remodeling, suggesting an important role of these processes in subclonal variegation. Indeed, we find that e.g. *ARID1A, PBRM1, KMT2C/D* and *SETD2* are enriched for subclonal driver mutations. Other genes often mutated in subclones are splicing factor *SF3B1* and, in breast and pancreatic adenocarcinomas, tumor suppressor *SMAD4*.

We similarly observed substantial variation in SV driver clonality across cancer types, implying cancer type-specific roles for SVs during tumor evolution (**Figure 6A**). Ten cancer types have a significant clonal bias for SV drivers (**Figure 6A**), when matched for power, suggesting that these cancers are driven by early SV events. These include SVs in the genomic region around *KIAA1549* in pilocytic astrocytomas, which likely result in the *BRAF-KIAA1549* fusion gene (Faulkner et al., 2015). Ovarian adenocarcinoma and soft-]tissue leiomyosarcoma show the highest rates of SV driver subclonality (33.7% and 40.0% respectively).

No significant subclonal enrichment was observed for SV drivers within a tumor type. However, enrichment was observed for specific SV drivers across cancer types (**Figures 6A and S5**). Clonally enriched SV drivers (**Figure 6A**, *q*-value < 0.05, rank-based permutation test) include those involving the *IGH* locus (97% of which occurred in lymphomas), or targeting *CDK12, TERT, MDM2, CDKN2A, LRP5/PPP6R3, MYC, EGFR* and gene poor region 8p11.21. In contrast, subclonally enriched SV drivers include those targeting *RB1, AKR1C1/2/3, KLF5, PTEN* and the gene poor 5p12 region. Interestingly, previous studies have linked *RB1* loss to tumor progression in liver (Bollard et al., 2017), liposarcoma (Schneider-Stock et al., 2002; Takahira et al., 2005), and breast cancer (Condorelli et al., 2017).

To further understand the clonality of gain-of-function driver SVs across cancer types, we specifically focused on previously known and curated oncogenic driver fusion SVs (**STAR Methods**). We found that known driver fusions are more likely to be clonal compared to other SVs (p = 0.0284, Fisher’s exact test, **Figure 6D**), with some recurrent fusions appearing exclusively clonal or highly enriched for clonal events (*CCDC6*-*RET, BRAF*-]*KIAA1549, TMPRSS2-ERG*), pointing to a model where gain-of-function SVs tend to appear early rather than late during tumor development.

Finally, to assess the potential impact of ITH on clinical decisions, we evaluated the clonality of actionable subclonal driver mutations, reasoning that targeting mutations that are not present in all tumor cells will likely result in ineffective treatment (Schmitt et al., 2016). Restricting our analysis to genes and mutations for which inhibitors are available, we find that 60.1% of tumors have at least one clinically actionable event (**Figure 6E**). Of these, 9.7% contain at least one subclonal actionable driver, and 4.7% show only subclonal actionable events. As our results represent conservative lower bound estimates of the subclonality at the level of the whole tumor, these results reinforce the importance of assessing the clonality of actionable mutations.

## DISCUSSION

We have developed consensus approaches to characterize genome-wide ITH for 38 cancer types, building on high quality SNVs, indels, SVs, CNAs, and curated driver mutations and mutation signatures, leveraging the largest set of whole-genome sequenced tumor samples compiled and analyzed to date. Remarkably, although these single region-based results are conservative and place a lower bound estimate on ITH, we detect subclonal tumor cell populations in 95.1% of 1,705 tumors. Individual subclones in the same tumor frequently exhibit differential activity of mutation signatures, implying that subclonal expansions can act as witnesses of temporally and spatially changing mutation processes. We extensively characterized the clonality of SNVs, indels, SVs, and CNAs. For SNVs and indels, we identified patterns of subclonal driver mutations in known cancer genes and average rates of subclonal driver events per tumor (Jamal-Hanjani et al., 2017; Landau et al., 2013; McGranahan et al., 2015; Yates et al., 2015). For SVs, we analyzed both candidate driver and passenger events, revealing how SVs influence tumor initiation and progression. Clonality estimates from CNAs suggest a complementary role of chromosomal instability and mutagenic processes in driving subclonal expansions. Finally, our results show rich subclonal architectures, with both linear and branching evolution in many cancers.

Analysis of dN/dS ratios in subclonal SNVs falling in exons of known cancer genes revealed clear signs of positive selection across the detected subclones and across cancer types. Although our analyses do not exclude the possibility that a small fraction of tumors evolve under weak or no selection, they show that selection is widespread across cancer types. Recent methodological advances to quantify selection in individual tumors from explicit tumor growth models have emerged and could shed further light on the evolutionary dynamics of individual tumors through single (Williams et al., 2018) and multiple (Sun et al., 2017) tumor biopsies. Our findings extend Peter Nowell’s model of clonal evolution (Nowell, 1976): as neoplastic cells proliferate under chromosomal and genetic instability, some of their daughter cells acquire mutations that convey further selective advantages, allowing them to become precursors for new subclonal lineages. Here, we have demonstrated that selection is ongoing up to and beyond diagnosis, in virtually all tumors and cancer types. The ubiquiotous presence of subclones provides evidence for ongoing selective sweeps, and akin to results of multi-region-based studies, we also detect widespread branching evolution, implying co-existence and competition of subclones.

Our observations highlight a considerable gap in knowledge about the drivers of subclonal expansions. Specifically, only 11% of the 2,542 detected subclones have a currently known SNV or indel driver mutation. Thus, late tumor development is either driven largely by different mechanisms (copy number alterations, genomic rearrangements (Jamal-Hanjani et al., 2017; Mamlouk et al., 2017), or epigenetic alterations), or most late driver mutations remain to be discovered. In support of the latter, our recent study (Gerstung et al., 2020) finds that late driver mutations occur in a more diverse set of genes than early drivers. For now, the landscape of subclonal driver mutations in localized cancer remains largely unexplored, in part due to limited resolution and statistical power to detect recurrence of subclonal drivers. Nonetheless, each tumor type has its own characteristic patterns of subclonal SNVs, indels, SVs and CNAs, revealing distinct evolutionary narratives. Tumor evolution does not end with the last complete clonal expansion, and it is therefore important to account for ITH and its drivers in clinical studies.

We show that regions of recurrent genomic rearrangements, harboring likely driver SVs, also exhibit subclonal rearrangements. This suggests that improved annotations must be sought for both SVs and SNVs, in order to comprehensively catalogue the drivers of subclonal expansion. By combining analysis of SV clonality with improved annotations of candidate SV drivers (Rheinbay et al., 2020), we highlight tumor types that would benefit from further characterization of subclonal SV drivers, such as pancreatic neuroendocrine cancers and leiomyosarcomas.

These observations have a number of promising clinical implications. For example, there is subclonal enrichment for SVs causing *RB1* loss across multiple cancer types, expanding on the known behavior of *RB1* mutations in breast cancer (Condorelli et al., 2017). These SVs may be linked to known resistance mechanisms to emerging treatments (*e.g.* CDK4/6 inhibitors in breast (Condorelli et al., 2017) and bladder (Pan et al., 2017) cancer). If profiled in a resistance setting, they may provide a pathway to second-line administration of cytotoxic therapies such as cisplatin or ionizing radiation, which show improved efficacy in tumors harboring *RB1* loss (Knudsen and Knudsen, 2008).

Our study builds upon a wealth of data of cancer whole-genome sequences generated under the auspices of the International Cancer Genome Consortium and The Cancer Genome Atlas, allowing detailed characterization of ITH from single tumor samples across 38 cancer types. It builds a consensus reconstruction of CNAs from 6 methods and consensus subclonal reconstruction from 11 methods. In establishing this reconstruction, we found that each individual method makes errors that are corrected by the consensus. Our consensus-building tools and techniques thus provide a set of best practices for future analyses of tumor whole-genome sequencing data. In addition, our high-quality curated consensus subclonal reconstructions on 2,658 tumor whole genomes spanning 38 cancer types constitute a rich resource for future studies.

## STAR METHODS SUMMARY

### Consensus copy number analysis

As the basis for our subclonal architecture reconstruction, we needed a confident copy number profile for each sample. To this end, we applied six copy number analysis methods (ABSOLUTE, ACEseq, Battenberg, cloneHD, JaBbA and Sclust) and combined their results into a robust consensus (see **STAR Methods** for details). In brief, each individual method segments the genome into regions with constant copy number, then calculates the copy number of both alleles for the genomic location. Some of the methods further distinguish between clonal and subclonal copy number states, *i.e.* a mixture of two or more copy number states within a genomic region. Disagreement between methods mostly stems from either difference in the segmentation step, or uncertainty on whole genome duplication (WGD) status. Both issues were resolved using our consensus strategy.

To identify a set of consensus breakpoints, we combined the breakpoints reported by the CNA methods with the consensus structural variants (SVs). If a hotspot of copy number breakpoints could be explained by an SV, we removed the copy number breakpoints in favor of the base-pair resolution SV. The remaining hotspots were merged into consensus calls to complement the SV-based breakpoints. This combined breakpoint set was then used as input to all methods in a second pass, where methods were required to strictly adhere to the provided breakpoints.

Allele-specific copy number states were resolved by assessing agreement between outputs of the individual callers. A consensus purity for each sample was obtained by combining the estimates of the copy number methods with the results of the subclonal architecture reconstruction methods that infer purity using only SNVs.

Each copy number segment of the consensus output was rated with a star-ranking representing confidence.

To create a subclonal copy number consensus, we used three of the copy number methods that predicted subclonal states for segments and flagged the segment as subclonal when at least two methods agreed the segment represented subclonal copy number.

### Consensus subclonal architecture clustering

We applied 11 subclonal reconstruction methods (BayClone-C, Ccube, CliP, cloneHD, CTPsingle, DPClust, PhylogicNDT, PhyloWGS, PyClone, Sclust, SVclone). Most were developed or further optimized during this study. Their outputs were combined into a robust consensus subclonal architecture (see **STAR Methods** for details). During this procedure, we used the PCAWG consensus SNVs and indels [Synapse ID syn7118450] and SVs [syn7596712].

The procedure to create consensus architectures consisted of three phases: a run of the 11 callers on a subset of SNVs that reside on copy number calls of high-confidence, merging of the output of the callers into a consensus and finally assignment of all SNVs, indels and SVs.

Each of the 11 subclonal reconstruction callers outputs the number of mutation clusters per tumor, the number of mutations in each cluster, and the clusters’ proportion of (tumor) cells (cancer cell fraction, CCF). These data were used as input to three orthogonal approaches to create a consensus: WeMe, CSR and CICC. The results reported in this paper are from the WeMe consensus method, but all three developed methods lead to similar results, and were used to validate each other (**STAR Methods**).

The consensus subclonal architecture was compared to the individual methods on two independent simulation sets, one 500-sample for training and one 965-sample for validation, and on the real PCAWG samples to evaluate robustness. The metrics by which methods were scored account for the fraction of clonal mutations, number of mutation clusters and the root mean square error (RMSE) of mutation assignments. To calculate the overall performance of a method, ranks of the three metrics were averaged per sample.

Across the two simulated datasets, the scores of the individual methods were variable, whereas the consensus methods were consistently among the best across the range of simulated number of subclones, tumor purity, tumor ploidy and sequencing depth. The highest similarities were observed among the consensus and the best individual methods in the simulation sets, and among the consensus methods in real data, suggesting stability of the consensus in the real set. Increasing the number of individual methods input to the consensus consistently improved performance and the highest performance was obtained for the consensus run on the full 11 individual methods, suggesting that each individual method has its own strengths that are successfully integrated by the consensus approaches (**STAR Methods**).

All SNVs, indels and SVs were assigned to the clusters that were determined by the consensus subclonal architecture using MutationTimer (Gerstung et al., 2020). Each mutation cluster is modelled by a beta-binomial distribution and probabilities for each mutation belonging to each cluster are calculated. This results in the final consensus subclonal architecture, and in addition, it also timed mutations relative to copy number gains (**STAR Methods**).

### SV clonality analysis

Due to the difficulty in determining SV VAFs from short-read sequence data, and subsequent CCF point estimation (Cmero et al., 2020), we elected to explore patterns of putative driver SV clonality using subclonal *probabilities*, allowing us to account for uncertainty in our observations of SV clonality (**STAR Methods**). After excluding unpowered samples, highly mutated samples, and cancer types with less than ten powered samples (**STAR Methods**), we analyzed 125,920 consensus SVs from 1,517 samples, across 28 cancer types. SVs were divided into candidate driver SVs and candidate passenger SVs using annotations from a companion paper (Rheinbay et al., 2020). SVs were considered candidate drivers if they were annotated as having significantly recurrent breakpoints (SRBs) at non-fragile sites, and candidate passenger SVs otherwise (**STAR Methods**).

Subclonal probabilities of driver and passenger SVs across tumor types were observed using weighted median and interquartile ranges (**STAR Methods**). Any tumor types with interquartile ranges exceeding subclonal probabilities of 0.5 were considered as having evidence of subclonal SVs. Permutation testing was used to determine significant differences in the weighted medians between driver and passenger SVs (**STAR Methods**). To test if any genomic loci were enriched for clonal or subclonal SVs across cancer types, we employed a GSEA-like (Subramanian et al., 2005) rank-based permutation test (**STAR Methods**).

### “Winner’s curse” correction

Because somatic mutation callers require a minimum coverage of supporting reads, in samples with low purity and/or small subclones, the reported CCF values and cluster sizes will be biased. As variants observed in a lower number of reads have a higher probability to be missed by somatic mutation callers, rare subclones will show lower apparent mutation numbers and higher apparent CCF values. We refer to this effect as the “Winner’s curse”. To adjust mutation clusters both in size and in CCF, we developed two methods, PhylogicCorrectBias and SpoilSport. Results from both methods were integrated to produce a consensus correction, and our correction approach was validated on simulated data (**STAR Methods**).

### Mutation signatures trajectory analysis

Given the mutation signatures obtained from PCAWG [syn8366024], we used TrackSig (Rubanova et al., 2020) to fit the evolutionary trajectories of signature activities. Mutations were ordered by their approximate relative temporal order in the tumor, by calculating a pseudo-time ordering using CCF and copy number. Time-ordered mutations were subsequently binned to create time points on a pseudo-timeline to which signature trajectories can be mapped.

At each time point, mutations were classified into 96 classes based on their trinucleotide context and a mixture of multinomial distributions was fitted, each component describing the distribution of one active signature. Derived mixture component coefficients correspond to mutation signature activity values, reflecting the proportion of mutations in a sample that were generated by a mutation process. By applying this approach to every time point along the evolutionary timeline of a sample, a trajectory of the activity of signatures over time was obtained.

We applied likelihood maximization and the Bayesian Information Criterion to simulations to establish the optimal threshold at which signature activity changes can be detected. This threshold was determined to be 6%. Subsequently, a pair of adjacent mutation bins was marked as constituting a change in activity if the absolute difference in activity between the bins of a at least one signature was greater than the threshold.

Signature trajectories were mapped to our subclonal reconstruction architectures by dividing the CCF space according to the proportion of mutations per time point belonging to a mutation cluster determined by the consensus reconstruction. By comparing distances in pseudo-time between trajectory change points and cluster boundaries, change points were classified as “supporting” a boundary if they are no more than three bins apart.

## Supplementary Figure Legends

**Figure S1.**
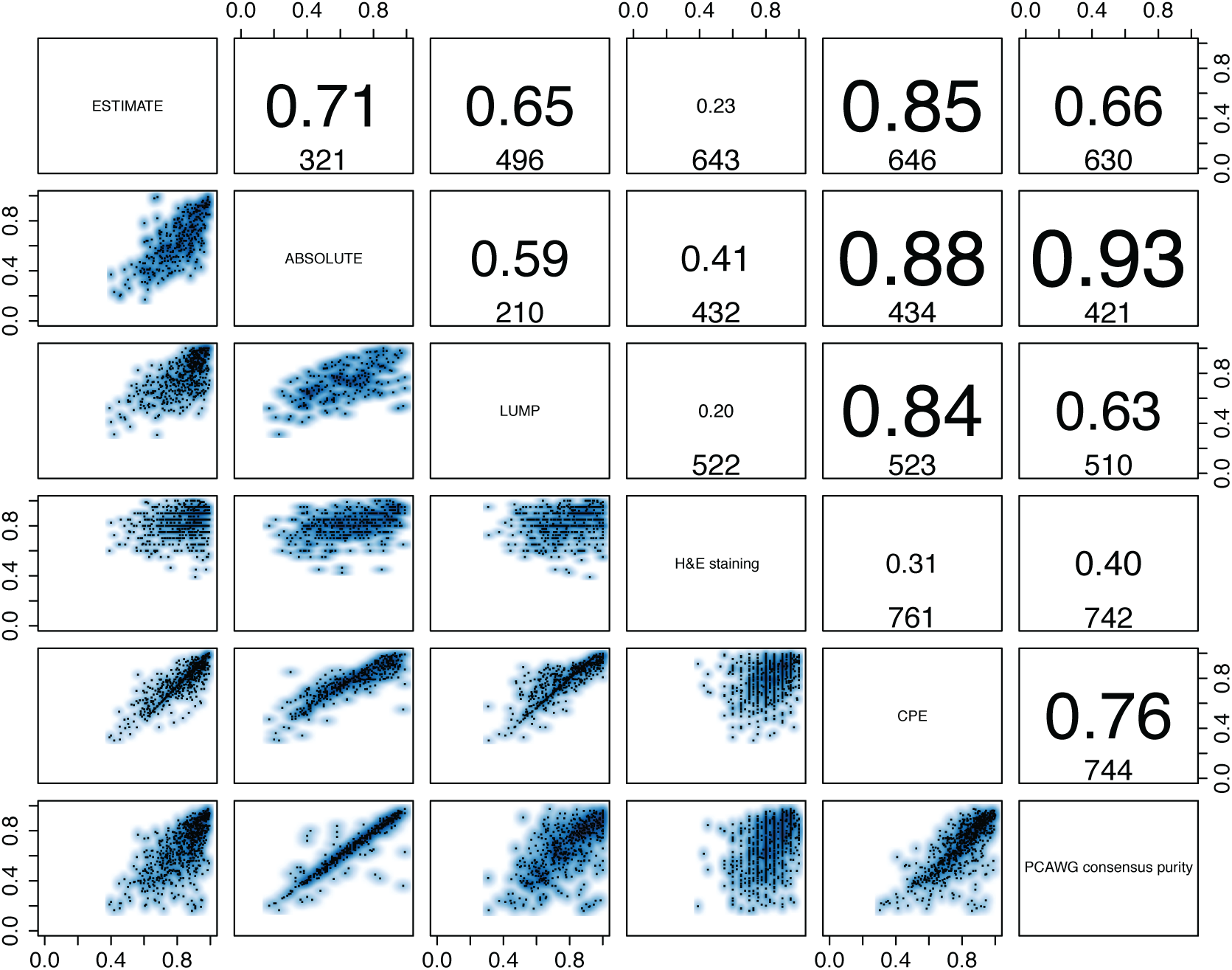
Validation of consensus purity values. The lower triangle shows pairwise scatterplots of the purities obtained through expression profiles of a panel of immune and stromal genes (ESTIMATE), somatic copy number data (ABSOLUTE), leukocyte unmethylation (LUMP), image analysis by hematoxylin and eosin staining (H&E staining), and consensus purity as derived by Aran *et al*. (Aran et al., 2015) (CPE). The top triangle shows the respective Pearson correlation coefficients and the number of samples that have both purity estimates available.

**Figure S2.**
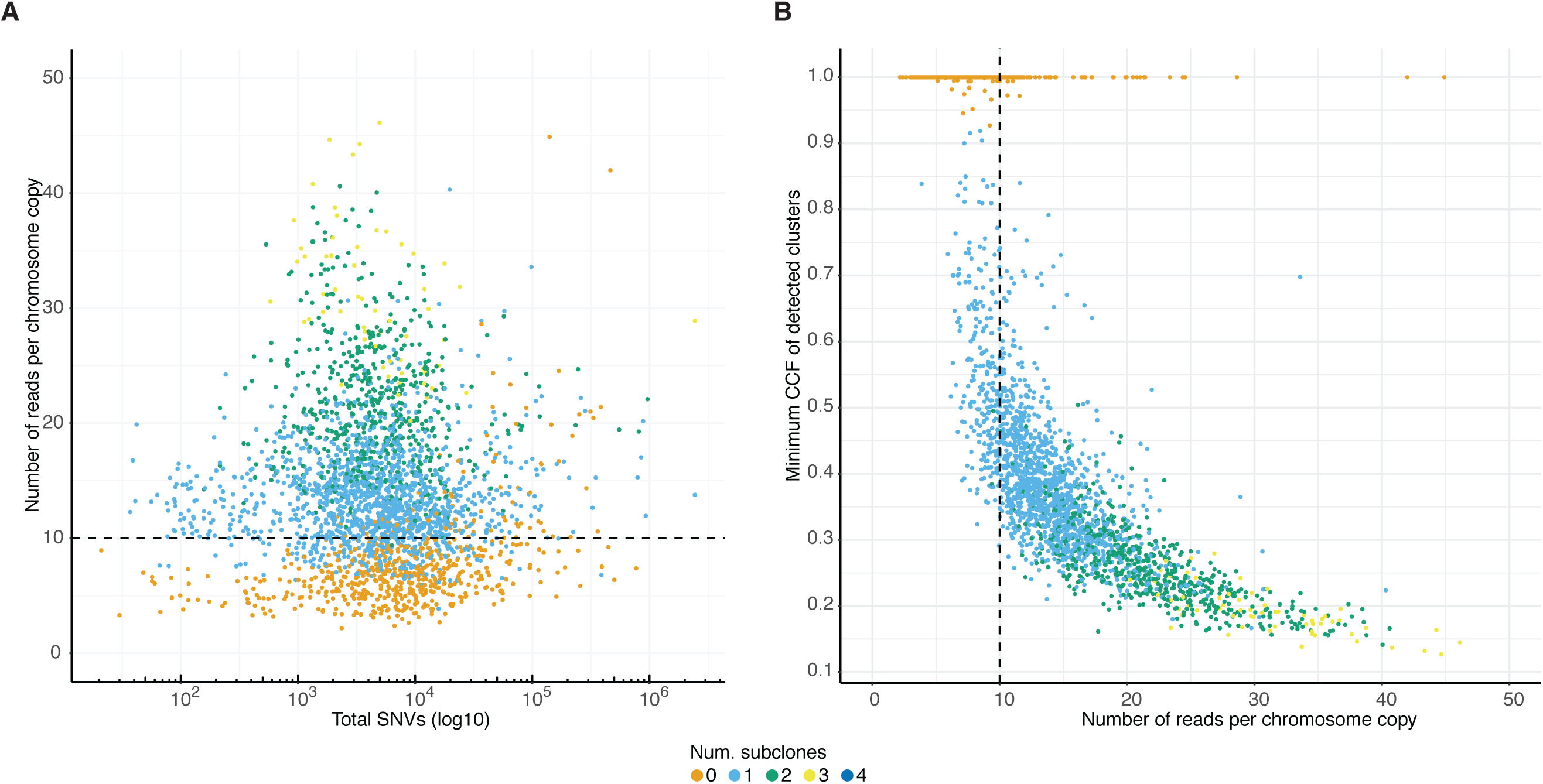
Power analysis of the consensus subclonal architecture approach. (A) Our ability to detect subclones depends, not on the number of detected SNVs, but on the number of reads per clonal copy (nrpcc) available. This metric takes tumor purity, ploidy and sequencing coverage into account (see **STAR Methods**). We control for this effect by including only tumors with nrpcc ≥ 10. In these tumors, we should be sufficiently powered to detect a subclone at a CCF as low as 30% (see **STAR Methods**). This becomes clear from (B) which shows the minimum CCF of the detected clusters in each tumor against the number of reads per chromosome copy.

**Figure S3.**
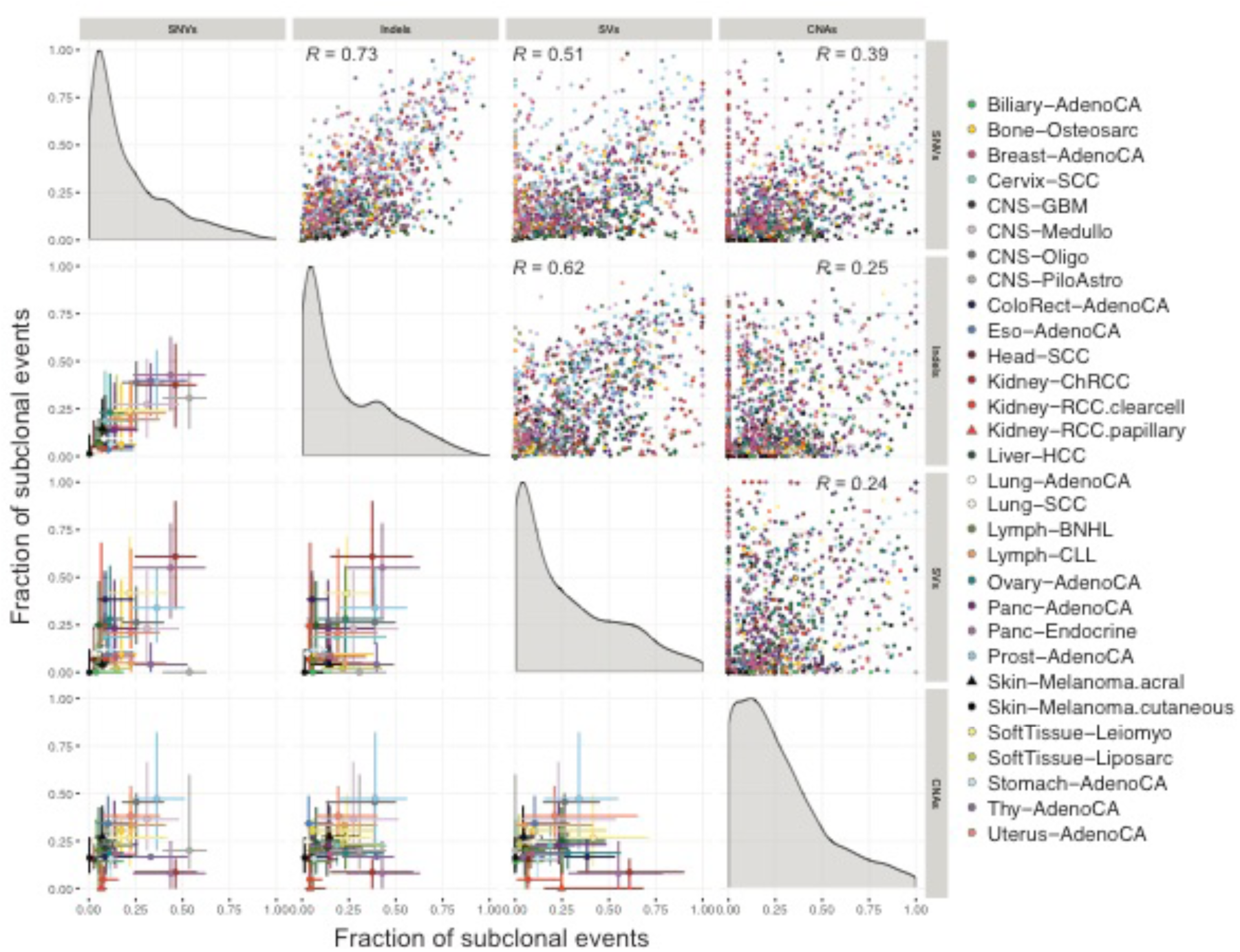
Correlation in ITH between SNVs, indels, CNAs and SVs by cancer type. Evidence of ITH is shown for 1,705 samples with sufficient power to detect subclones above 30% CCF (see **STAR Methods**), as in **Figure 3**. Pairwise scatter plots in the upper triangle show the fractions of subclonal SNVs, indels, CNAs and SVs per tumor sample. Pearson’s correlation coefficient, *R*, is separately computed for each panel across all samples. Panels on the diagonal show the kernel density estimate of the distribution of subclonal fractions. In the lower triangle, each point shows the median subclonal fraction per cancer type and intervals indicate the interquartile range. Panels only include samples with at least 5 arm-level CNAs (1,238 / 1,705) and at least 5 SVs (1,609 / 1,705).

**Figure S4.**
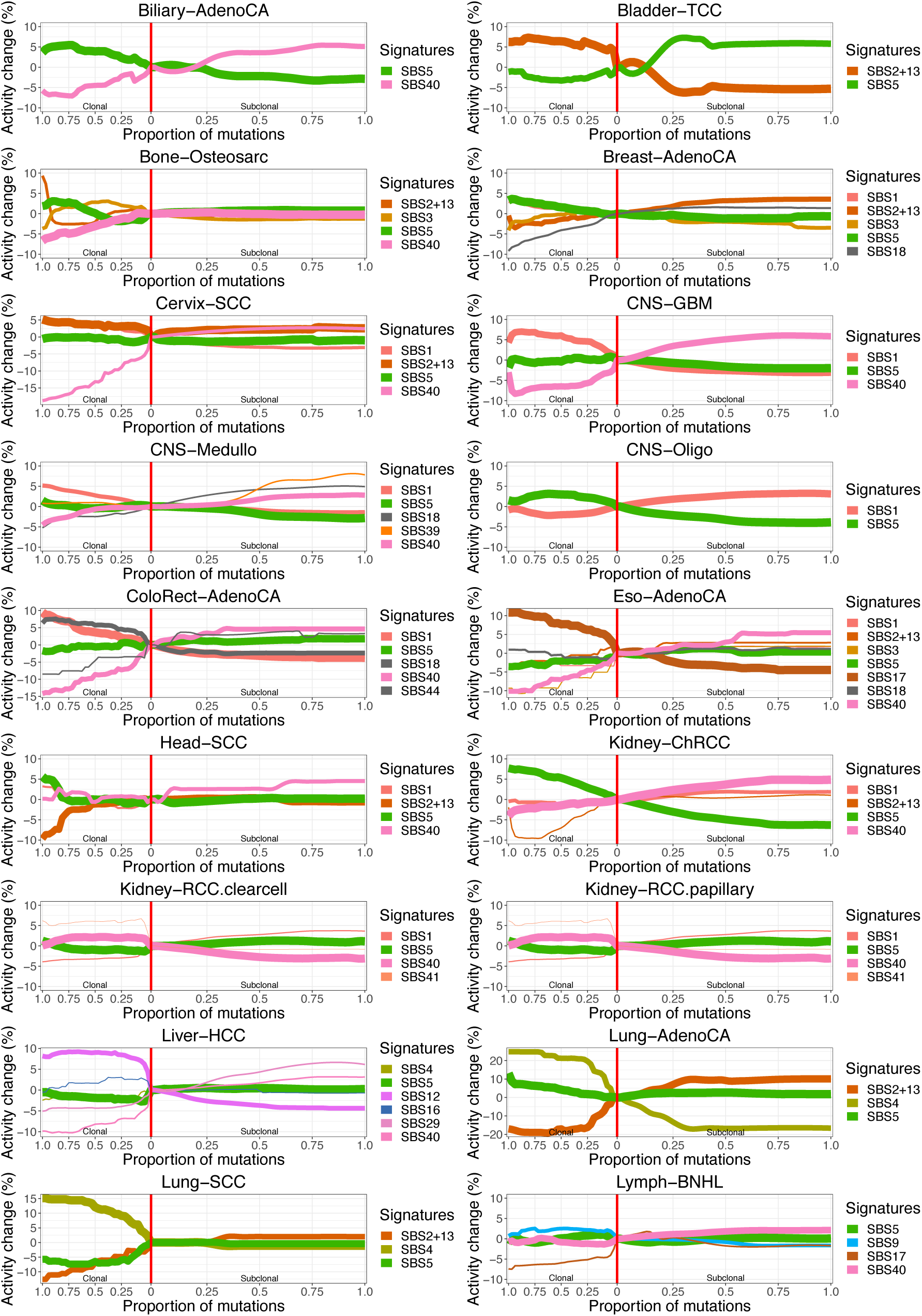
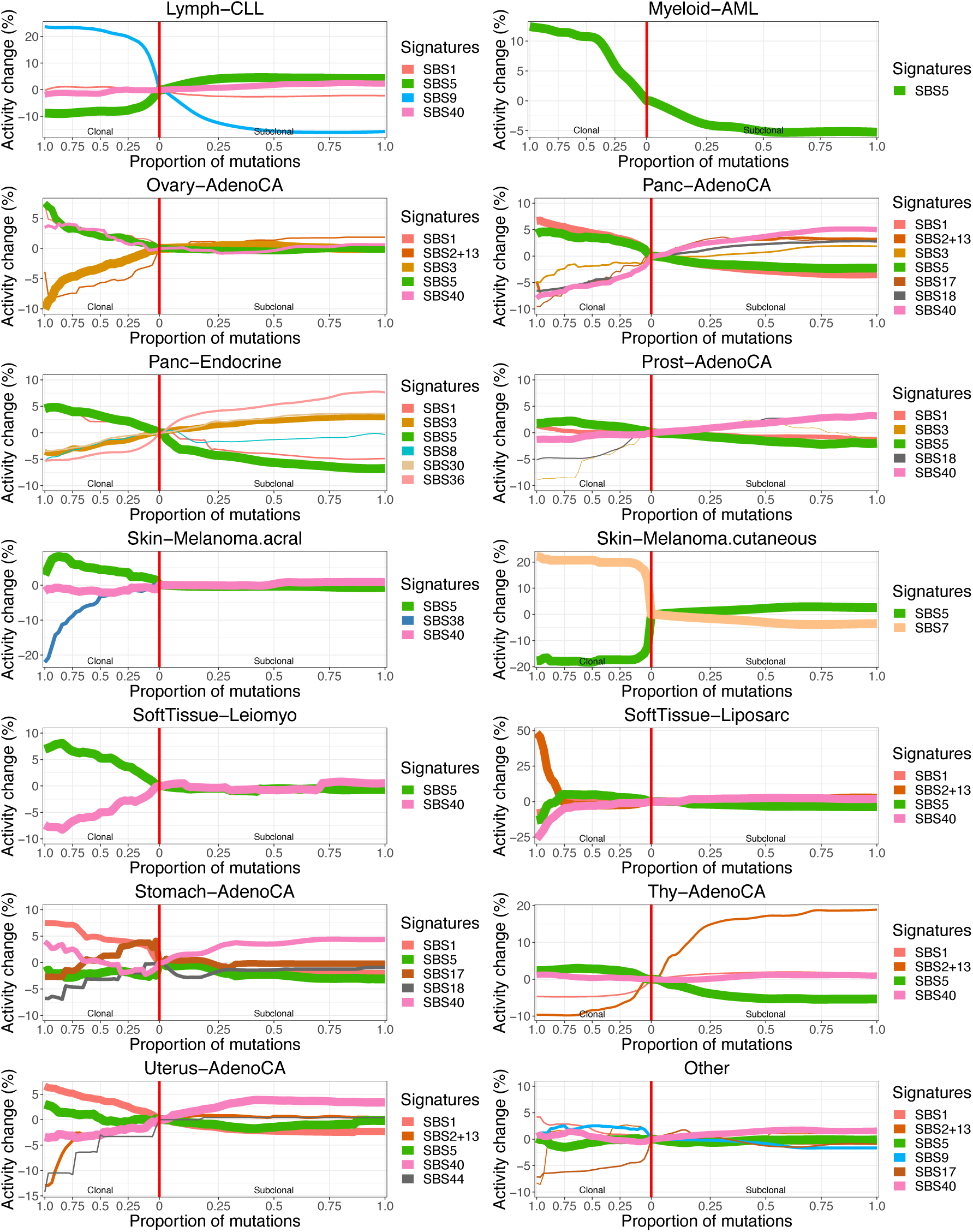
Summary signature trajectories per cancer type. The average trajectories for mutation signatures were calculated across tumors of the same cancer type. The color of the line denotes the signature and its width reflects the number of contributing tumors. The trajectories have been centered around the activity at the boundary between clonal and subclonal mutations in order to highlight relative changes in signature activity.

**Figure S5.**
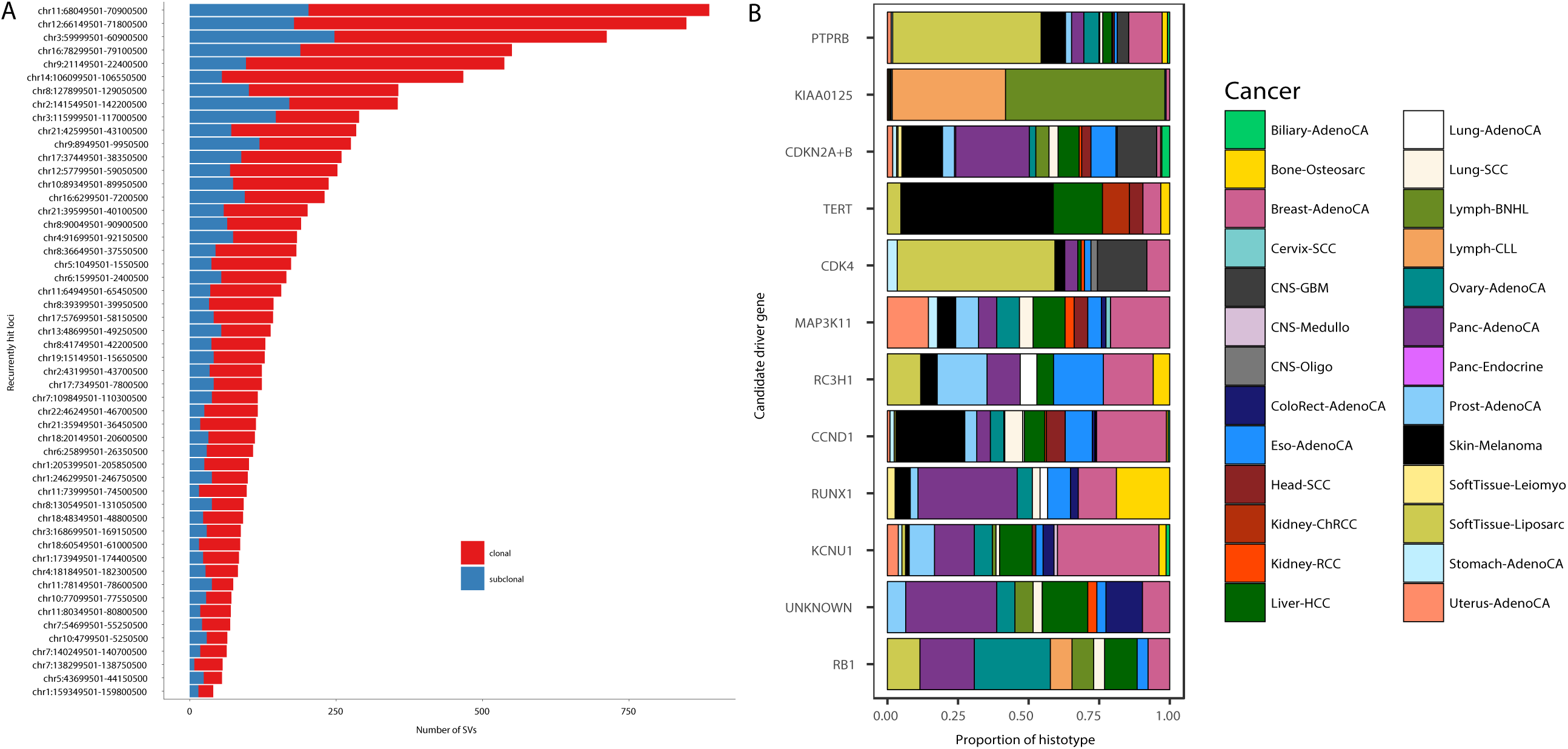
Clonality analysis of significantly recurrent breakpoints. (A) Number and clonality of SVs observed at 52 loci with significantly recurrent breakpoints (SRBs) (Rheinbay et al., 2020). SVs with a subclonal probability larger than 50% were considered subclonal and clonal otherwise. (B) Proportion of cancer types contributing to the enrichment of clonal or subclonal SVs in a locus (see **Figure 6A**). The genes on the y-axis represent the most likely driver gene for each locus (Rheinbay et al., 2020).

## ACKNOWLEDGMENTS

This work was supported by the Francis Crick Institute, which receives its core funding from Cancer Research UK (FC001202), the UK Medical Research Council (FC001202), and the Wellcome Trust (FC001202). This project was enabled through the Crick Scientific Computing STP and through access to the MRC eMedLab Medical Bioinformatics infrastructure, supported by the Medical Research Council (grant number MR/L016311/1). MT and JD are postdoctoral fellows supported by the European Union’s Horizon 2020 research and innovation program (Marie Skłodowska-Curie Grant Agreement No. 747852-SIOMICS and 703594-DECODE). JD is a postdoctoral fellow of the Research Foundation – Flanders (FWO). IVG is supported by a Wellcome Trust PhD fellowship (WT097678) and the Ann and Sol Schreiber Mentored Investigator Award of the Ovarian Cancer Research Alliance. SM is funded by a Vanier Canada Graduate Scholarship. SCS is supported by the NSERC Discovery Frontiers Project, “The Cancer Genome Collaboratory” and by NIH GM108308. DJA is supported by Cancer Research UK. FM, GM and KeY would like to acknowledge the support of the University of Cambridge, Cancer Research UK and Hutchison Whampoa Limited. GM, KeY and FM are funded by CRUK core grants C14303/A17197 and A19274. KeY is further supported by EPSRC EP/R018634/1. SSe and YJ are supported by NIH R01 CA132897. HZ is supported by grant NIMH086633 and an endowed Bao-Shan Jing Professorship in Diagnostic Imaging. PTS is supported by U24CA210957 and 1U24CA143799. WW is supported by the U.S. National Cancer Institute (1R01 CA183793 and P30 CA016672). DCW is funded by the Li Ka Shing foundation. PVL is a Winton Group Leader in recognition of the Winton Charitable Foundation’s support towards the establishment of The Francis Crick Institute. We acknowledge the contributions of the many clinical networks across ICGC and TCGA who provided samples and data to the PCAWG Consortium, and the contributions of the Technical Working Group and the Germline Working Group of the PCAWG Consortium for collation, realignment and harmonized variant calling of the cancer genomes used in this study. We thank the patients and their families for their participation in the individual ICGC and TCGA projects. We gratefully acknowledge Nicholas McGranahan and Charles Swanton for valuable comments on our manuscript.

## MEMBERS OF THE PCAWG EVOLUTION AND HETEROGENEITY WORKING GROUP

Stefan C. Dentro^1,2,3,*^, Ignaty Leshchiner^4,*^, Moritz Gerstung^5,*^, Clemency Jolly^1,*^, Kerstin Haase^1,*^, Maxime Tarabichi^1,2,*^, Jeff Wintersinger^6,7,*^, Amit G. Deshwar^6,7,*^, Kaixian Yu^8,*^, Santiago Gonzalez^5,*^, Yulia Rubanova^6,7,*^, Geoff Macintyre^9,*^, Jonas Demeulemeester^1,10,*^, David J. Adams^2^, Pavana Anur^11^, Rameen Beroukhim^4,12^, Paul C. Boutros^6,13^, David D. Bowtell^14^, Peter J. Campbell^2^, Shaolong Cao^8^, Elizabeth L. Christie^14,15^, Marek Cmero^15,16^, Yupeng Cun^17^, Kevin J. Dawson^2^, Nilgun Donmez^18,19^, Ruben M. Drews^9^, Roland Eils^20,21^, Yu Fan^8^, Matthew Fittall^1^, Dale W. Garsed^14,15^, Gad Getz^4,22,23,24^, Gavin Ha^4^, Marcin Imielinski^25,26^, Lara Jerman^5,27^, Yuan Ji^28,29^, Kortine Kleinheinz^20,21^, Juhee Lee^30^, Henry Lee-Six^2^, Dimitri G. Livitz^4^, Salem Malikic^18,19^, Florian Markowetz^9^, Inigo Martincorena^2^, Thomas J. Mitchell^2,31^, Ville Mustonen^32^, Layla Oesper^33^, Martin Peifer^17^, Myron Peto^11^, Benjamin J. Raphael^34^, Daniel Rosebrock^4^, S. Cenk Sahinalp^19,35^, Adriana Salcedo^36^, Matthias Schlesner^20^, Steven Schumacher^4^, Subhajit Sengupta^28^, Ruian Shi^6^, Seung Jun Shin^8,37^, Lincoln D. Stein^36^, Oliver Spiro^4^, Ignacio Vázquez-García^2,31,38,39^, Shankar Vembu^6^, David A. Wheeler^40^, Tsun-Po Yang^17^, Xiaotong Yao^25,26^, Ke Yuan^9,41^, Hongtu Zhu^8^, Wenyi Wang^8,#^, Quaid D. Morris^6,7,#^, Paul T. Spellman^11,#^, David C. Wedge^3,42,#^, Peter Van Loo^1,#^

^1^The Francis Crick Institute, London NW1 1AT, United Kingdom; ^2^Wellcome Trust Sanger Institute, Cambridge CB10 1SA, United Kingdom; ^3^Big Data Institute, University of Oxford, Oxford OX3 7LF, United Kingdom; ^4^Broad Institute of MIT and Harvard, Cambridge, MA 02142, USA; ^5^European Molecular Biology Laboratory, European Bioinformatics Institute, Cambridge CB10 1SD, United Kingdom; ^6^University of Toronto, Toronto, ON M5S 3E1, Canada; ^7^Vector Institute, Toronto, ON M5G 1L7, Canada; ^8^The University of Texas MD Anderson Cancer Center, Houston, TX 77030, USA; ^9^Cancer Research UK Cambridge Institute, University of Cambridge, Cambridge CB2 0RE, United Kingdom; ^10^Department of Human Genetics, University of Leuven, B-3000 Leuven, Belgium; ^11^Molecular and Medical Genetics, Oregon Health & Science University, Portland, OR 97231, USA; ^12^Dana-Farber Cancer Institute, Boston, MA 02215, USA; ^13^University of California, Los Angeles, Los Angeles, CA 90095, USA; ^14^Peter MacCallum Cancer Centre, Melbourne, VIC 3000, Australia; ^15^University of Melbourne, Melbourne, VIC 3010, Australia; ^16^Walter + Eliza Hall Institute, Melbourne, VIC 3000, Australia; ^17^University of Cologne, 50931 Cologne, Germany; ^18^Simon Fraser University, Burnaby, BC V5A 1S6, Canada; ^19^Vancouver Prostate Centre, Vancouver, BC V6H 3Z6, Canada; ^20^German Cancer Research Center (DKFZ), 69120 Heidelberg, Germany; ^21^Heidelberg University, 69120 Heidelberg, Germany; ^22^Massachusetts General Hospital Center for Cancer Research, Charlestown, Massachusetts 02129, USA; ^23^Massachusetts General Hospital, Department of Pathology, Boston, Massachusetts 02114, USA; ^24^Harvard Medical School, Boston, 02215, USA; ^25^Weill Cornell Medicine, New York, NY 10065, USA; ^26^New York Genome Center, New York, NY 10013, USA; ^27^University of Ljubljana, 1000 Ljubljana, Slovenia; ^28^NorthShore University HealthSystem, Evanston, IL 60201, USA; ^29^The University of Chicago, Chicago, IL 60637, USA; ^30^University of California Santa Cruz, Santa Cruz, CA 95064, USA; ^31^University of Cambridge, Cambridge CB2 0QQ, United Kingdom; ^32^Organismal and Evolutionary Biology Research Programme, Department of Computer Science, Institute of Biotechnology, University of Helsinki, 00014 Helsinki, Finland; ^33^Carleton College, Northfield, MN 55057, USA; ^34^Princeton University, Princeton, NJ 08540, USA; ^35^Indiana University, Bloomington, IN 47405, USA; ^36^Ontario Institute for Cancer Research, Toronto, ON M5G 0A3, Canada; ^37^Korea University, Seoul, 02481, Republic of Korea; ^38^Computational Oncology, Memorial Sloan Kettering Cancer Center, New York, NY 10065, USA; ^39^Irving Institute for Cancer Dynamics, Columbia University, New York, NY 10027, USA; ^40^Human Genome Sequencing Center, Baylor College of Medicine, Houston, TX 77030, USA; ^41^School of Computing Science, University of Glasgow, Glasgow G12 8RZ, United Kingdom; ^42^Oxford NIHR Biomedical Research Centre, Oxford OX4 2PG, United Kingdom.

*: These authors contributed equally

^#^: These authors jointly directed the work

