## Supplementary Methods for "Characterizing genetic intra-tumor heterogeneity across 2,658 human cancer genomes"

### Table of contents

**Glossary** **3**

**0. Brief overview of the data set** **4**

**1. Copy number consensus** **7**
*1.1 Overview* *7*
*1.2 Description of copy number methods* *7*
1.2.1 ABSOLUTE 7
1.2.2 ACEseq 8
1.2.3 Battenberg 8
1.2.4 cloneHD 9
1.2.5 JaBbA 10
1.2.6 Sclust 12
*1.3 Copy number consensus approach* *13*
1.3.1 Consensus copy number procedure 13
1.3.2 Copy-number-calling methods differ in genome segmentation 13
1.3.3 Method for determining consensus segment breakpoints 14
1.3.4 Resolving ploidy disagreement between copy number 19
callers 19
1.3.5 Consensus copy number 22
1.3.6 Subclonal copy number consensus 25
*1.4 Validation* *25*
1.4.1 Comparison of purity against TCGA 25

**2. Subclonal architecture methods** **26**
*2.1 Description of reconstruction methods* *26*
2.1.1 BayClone-C 26
2.1.2 Ccube 27
2.1.3 CliP (Clonal structure identification through Penalizing on pairwise difference) 29
2.1.4 cloneHD 30
2.1.5 CTPsingle 32
2.1.6 DPClust 34
2.1.7 PhylogicNDT 36
2.1.8 PhyloWGS 38
2.1.9 PyClone 40

2.1.10 Sclust 41
2.1.11 SVclone 41
*2.2 Subclonal architecture consensus approaches* *43*
2.2.1 Consensus approaches 45
2.2.2 Evaluation metrics 53
2.2.3 Results 55
*2.3 "Winner's curse" correction* *62*
2.3.1 PhylogicCorrectBias 62
2.3.2 SpoilSport 63
2.3.3 "Winner's curse" correction consensus 64
*2.4 Validation* *65*
2.4.1 Simulation of subclonally heterogeneous samples – PhylogicSim500 65
2.4.2 A grid approach to simulations of tumors – SimClone1000 68

**3. Results of reconstructing the subclonal architecture of tumors** **73**
*3.1 Power analysis* *73*
***3.2 Results by individual methods are consistent with consensus results*** *73*
*3.3 Correlation between fraction of subclonal mutations and mutation burden* *75*

**4. Selection and driver genes** **76**
*4.1 Selection and dN/dS* *76*
*4.2 Gene set analysis of subclonally mutated genes* *76*

**5. Evidence of additional ITH** **78**
*5.1 Analysis of phaseable mutation pairs* *78*
5.1.1 Infinite Sites Assumption 79
5.1.2 Analysis of TRACERx 100 phylogenetic trees 80
*5.2 Tracking signature activities across cancer timelines* *80*

**6. Analysis of drivers in clinically targetable genes** **83**

**7. SV analysis and fusion clonality detection** **85**
*7.1 Clonality analysis of recurrent structural variants* *85*
*7.2 Fusion clonality analysis* *87*

**References** **88**

### Glossary

| | |
|---|---|
| **BAF** | B-allele frequency |
| **CCF** | Cancer cell fraction, an estimate of the proportion of tumor cells that a subclone consists of or the estimated fraction of tumor cells that carry a mutation |
| **CNA** | Copy number alteration |
| **ICGC** | International Cancer Genome Consortium |
| **indel** | Small insertion or deletion |
| **logR** | The log of the ratio of relative read coverage in the tumor over that in the matched normal |
| **LOH** | Loss of heterozygosity |
| **nrpcc** | Number of reads per chromosome copy, a combination of tumor purity, ploidy and sequencing coverage |
| **PCAWG** | The Pan-Cancer Analysis of Whole Genomes project |
| **SNP** | Single nucleotide polymorphism |
| **SNV** | Single nucleotide variant |
| **SSM** | Simple somatic variant (SNV or indel) |
| **subclone** | A subpopulation of tumor cells that have arisen after the most recent common ancestor. The mutations carried by the cells in this subpopulation are sequenced and can be used to detect the presence of the subpopulation. |
| **WGD** | Whole genome duplication |

### 0. Brief overview of the data set

This manuscript describes analyses based on the ICGC-TCGA Pan-Cancer Analysis of Whole Genomes (PCAWG) dataset, which make use of the output from various PCAWG working groups (Alexandrov et al., 2020; Gerstung et al., 2020; Rheinbay et al., 2020; The ICGC/TCGA Pan-Cancer Analysis of Whole Genomes Consortium, 2020). To aid the reading of this manuscript, we briefly describe the dataset here.
The ICGC-PCAWG dataset comprises samples selected from individual ICGC and TCGA projects for which completion was imminent in 2015, as detailed in (The ICGC/TCGA Pan-Cancer Analysis of Whole Genomes Consortium, 2020). Donors were included if a tumor and matched normal were sequenced to a minimum per-base coverage of 30x and 25x respectively, on the Illumina HiSeq platform with 100-150 bp paired-end reads. 2,834 donors passed these criteria, which were reduced to 2,658 after an extensive quality control procedure (also described in (The ICGC/TCGA Pan-Cancer Analysis of Whole Genomes Consortium, 2020)). In total, 2,778 cancer samples from these 2,658 distinct donors were included in the final dataset, comprising 2,605 primary tumors and 173 metastases or recurrences. Purity and ploidy values (Fig. 1) as well as whole genome duplication status (Fig. 2), were estimated via a consensus approach based on 6 CNA callers, as described further below.

All sequencing data was collected and analyzed through a series of standardized primary analysis pipelines to first realign reads to the same reference genome and subsequently call SNVs, indels and SVs, using a homogenized procedure, as is detailed in (The ICGC/TCGA Pan-Cancer Analysis of Whole Genomes Consortium, 2020). High quality somatic SNV and indel calls were established via a robust consensus strategy based on multiple methods to achieve greater accuracy. Eighteen different callers were considered and asked to produce calls on 63 tumors selected across 23 cancer types and 26 contributing projects. For 50 tumors, there was sufficient DNA to perform deep sequencing via DNA hybridization capture. Around 250,000 SNVs and indels were selected, stratified by the number of methods by which they were called, followed by uniform sampling across the overlaps. A consensus approach was then defined to maximize precision and sensitivity, based on the three core pipelines supplied by the Broad, DKFZ-EMBL and Sanger. Two additional callers (one SNV caller from MD-Anderson and an additional indel caller from the Barcelona Supercomputing Center) were added to improve the ability to detect low-allele-frequency variants. Variant allele frequency was taken into account when sampling variants for the validation; the precision obtained on SNVs at different allele frequencies is shown in Fig. 3.

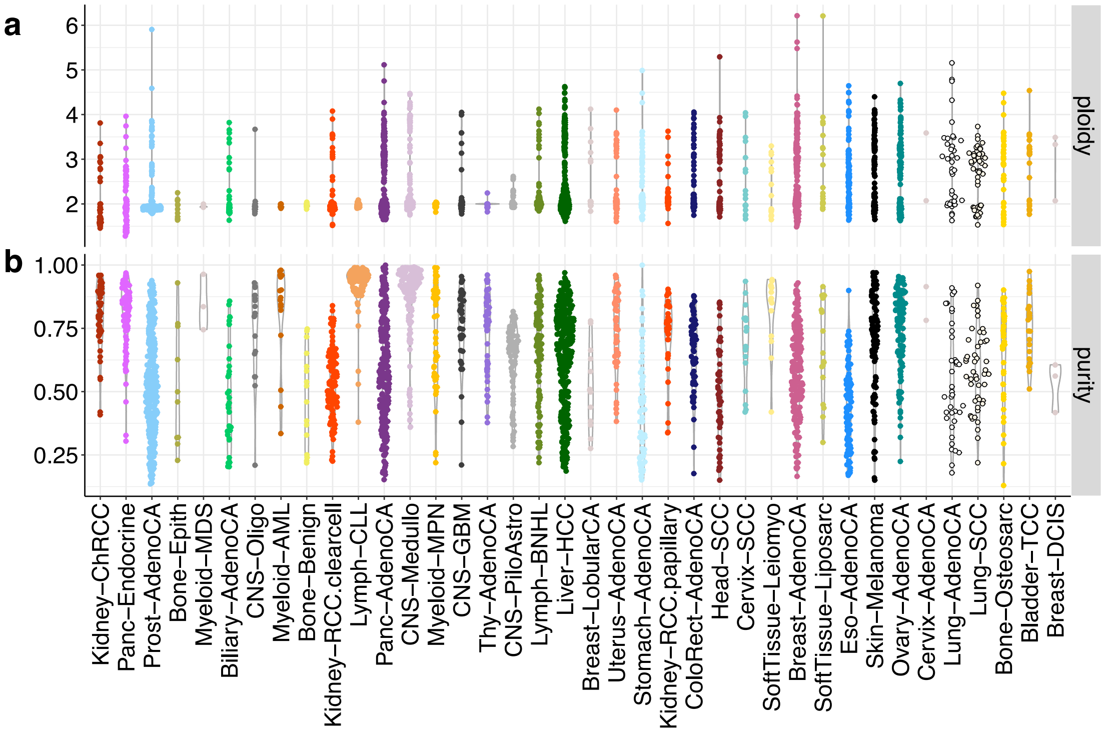


*Fig. 1 Ploidy (a) and purity (b) values across cancer types, sorted by median ploidy.*

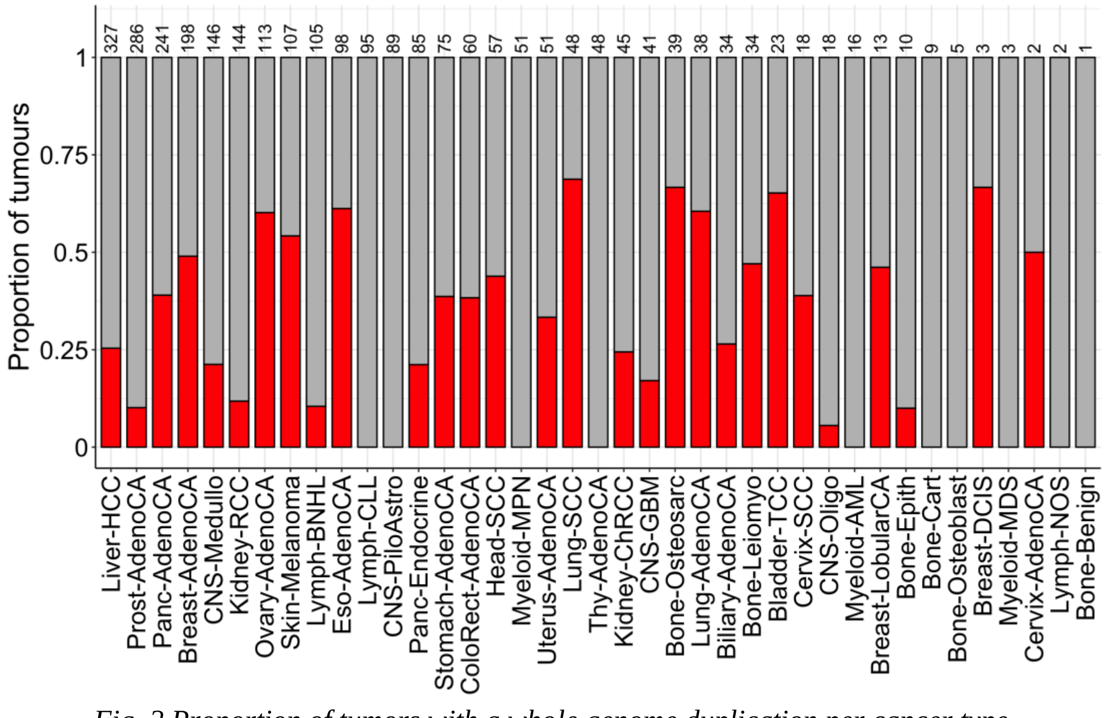


*Fig. 2 Proportion of tumors with a whole genome duplication per cancer type.*

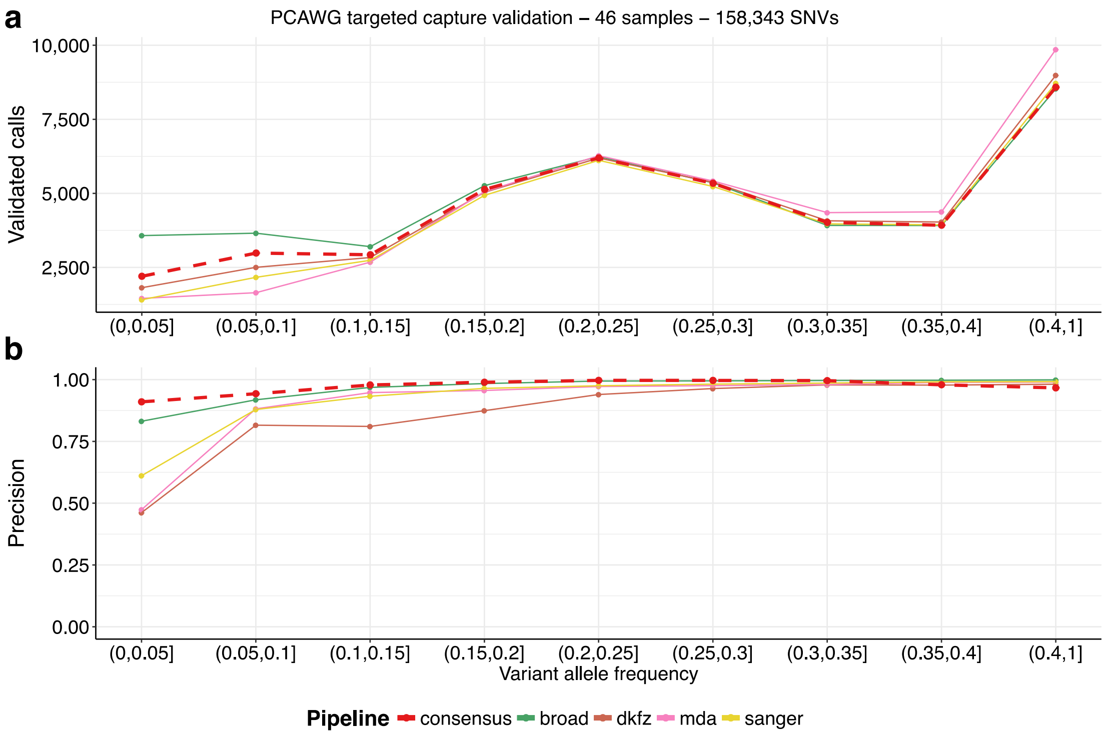


*Fig. 3* ***SNV validation results.*** *The figure shows the number of validated calls for each variant allele frequency bin (a) and the obtained precision (b). The consensus achieves the highest precision of all pipelines with a minimum of 90% of positive calls (lowest VAF bin) and over 94% in all other VAF bins.*

Driver calls have been made available throughout the PCAWG project; the full findings and methods are described in (Rheinbay et al., 2020). Somatic driver SNVs and indels were discovered by combining the outputs from 16 different discovery methods, including factors such as mutational burden, functional impact and mutation hotspots. To combine calls, their approach integrated p-values assigned to each event by the 16 callers, taking into account autocorrelation between methods based on similar principles, and applying multiple-testing correction. This was applied to protein-coding genes, promoters, untranslated regions (UTRs), distal enhancers and non-coding RNAs. Mutational signatures, and estimates of their activity in each sample, have been produced by the PCAWG signatures group, with findings and methods detailed in (Alexandrov et al., 2020). They applied two different computational approaches based on non-negative matrix factorization to 4,645 genome and 19,184 exome sequences, to extract 49 single base substitution, 11 doublet base substitution and 17 indel signatures. The doublet base and indel signatures are completely novel, whilst the single base signatures are broadly concordant with the published signatures in COSMIC, with some additional signatures, and some signatures now being split. The authors report a high concordance between the two approaches on all PCAWG samples, except in the case of hyper-mutators (5.6% of samples). Both reference sets of signatures and the per-sample quantifications of activity have been made available to the PCAWG project. In this manuscript, for simplicity we make use of the signatures output from one of the approaches only (SigProfiler).

### 1. Copy number consensus

#### 1.1 Overview

The copy number consensus procedure builds on top of the output from six different copy number callers. Upon inspection of profiles produced by the individual callers, we observed that profiles differ depending on their segmentation and that disagreement on copy number states of large proportions of the genome is the result of disagreement on whether a whole genome duplication had occurred. We therefore first constructed a complete set of breakpoints from an initial pass of five of the six methods (the JaBbA output was not yet available). The consensus breakpoints were then fed back in to the methods to produce copy number calls from all six methods. After resolving ploidy disagreements, we applied a procedure to every consensus segment that looks for agreement in major and minor allele states between the output of the six callers. Finally, we produced consensus purity calls for every tumor by combining purity calls from the six copy number callers with those from subclonal architecture reconstruction methods that produce purity calls from SNV data. The full procedure results in a purity estimate for every tumor in the PCAWG dataset and a copy number call for every consensus segment, where every call is assigned a confidence level and a quality star.

#### 1.2 Description of copy number methods

##### 1.2.1 ABSOLUTE

Ignaty Leshchiner, Dimitri Livitz, Gad Getz

We used the ABSOLUTE algorithm to calculate the purity, ploidy, and absolute DNA copy number of each sample (Carter et al., 2012). Fragment based coverage (derived from full read template span) was collected over the genome and corrected for GC and mappability biases. Tangent-normalization of the tumor copy number profile was performed by using a panel built from PCAWG normal samples. Further, allele-specific copy number was computed based on heterozygous sites identified in the paired normal and segmentation was performed by using a circular binary segmentation algorithm. The Nelder–Mead algorithm was used to search the space of possible purity and ploidy solutions and prioritize them. Dirichlet Process clustering was performed on subclonal copy number segments, to annotate identical subclonal copy-number cluster states. Somatic SNVs were imputed on the computed absolute copy number profile and the cancer cell fraction and multiplicities were inferred for each mutation independently prior to running subclonal architecture reconstruction.

##### 1.2.2 ACEseq

Kortine Kleinheinz, Roland Eils, Matthias Schlesner

ACEseq (allele-specific copy number estimation from sequencing) (Kleinheinz et al., 2017) determines absolute copy number from a combination of the coverage ratio of tumor and matched normal in genomic windows and the B-allele frequencies (BAF) of the corresponding SNPs. In addition to the copy number, tumor ploidy and tumor cell content are estimated.

During pre-processing of the data, allele frequencies are obtained for all single nucleotide polymorphism (SNP) positions recorded in dbSNP version 135 (Levy et al., 2015). To improve sensitivity for the detection of genomic imbalances, SNPs in the matched normal are phased with IMPUTE2 (Howie et al., 2009) . The coverage for 10 kb windows with sufficient mapping quality and read depth (maximum mappability over at least 50 % of the window; average coverage of at least 5 reads per base) is determined and corrected for GC content-dependent as well as replication timing-dependent coverage bias. Subsequently the genome is segmented with the PSCBS package (Olshen et al., 2011) into regions of equal coverage and imbalance state. Prior to segmentation, structural variant (SV) breakpoints defined by the consensus SV set were incorporated as segment borders. The resulting segments were submitted for consensus breakpoint estimation.

The segments obtained with the final consensus breakpoints were annotated with coverage and BAF values to estimate the tumor cell content and ploidy of the sample. All ploidy / tumor cell content combinations in the range of 1 and 6.5 for ploidy and 15% and 100% for tumor cell content were modelled, and the combination which best explains the data was chosen. Balanced segments were constrained to even-numbered copy number states during the fitting, and solutions which required a BAF value > 1 or negative copy numbers for at least one segment were excluded.

##### 1.2.3 Battenberg

Stefan Dentro, Kevin Dawson, Henry Lee-Six, David Wedge, Peter Van Loo

We applied the previously described (Dentro et al., 2017; Nik-Zainal et al., 2012) Battenberg algorithm to the PCAWG data. Briefly, the Battenberg pipeline starts with collecting read count information for all 1000 genomes phase 3 SNPs from the tumor and its matched normal. B-allele frequency (BAF) and relative coverage ratio (logR) are calculated for each SNP, after which the logR is corrected for GC content. The matched normal is used to obtain germline heterozygous SNPs, which are subsequently phased using IMPUTE2 to obtain haplotype blocks. Outlier blocks are switched because two consecutive haplotype blocks can have a different allele chosen as allele *b*. The phased SNPs result in precise BAF estimates that allow for detection of subclonal copy number.

The data is segmented using piecewise constant fitting (PCF), with structural variants (SVs) taken in as prior established breakpoints. A clonal copy number profile is fit by a grid search over purity and ploidy combinations. The purity/ploidy combination that yields the largest proportion of the genome with clonal copy number is picked. Subclonal copy number is fit by first testing whether the BAF is significantly different from the currently fit clonal copy number state, allowing for at most a 1% deviation from the clonal BAF. If the BAF is significantly different, then there are four options under the assumption that there are at most two cellular populations with only a single allele copy number difference: Allele A and B are rounded up, A and B are rounded down, A is rounded up and B is rounded down, or A is rounded down and B is rounded up. The option that explains the observed BAF best is picked and used to fit the final copy number states.

The algorithm used for this analysis is different from the original version in three ways: Correction for GC content, inclusion of SVs during segmentation and merging of adjacent segments that are assigned the same copy number states. Sequencing data can be affected by inconsistent coverage that correlates with GC content (Benjamini and Speed, 2012) and coverage is contained within the LogR calculations. A correction step is therefore required. We correct the LogR by fitting a linear model that explains the data as being affected by two types of GC artefacts: A high frequency wave, encoded in a small window size and a low frequency wave corresponding to a large window size. We pre-calculated the GC content in various windows centered on each 1000 genomes SNP. For each window size we calculate the GC content correlation and pick the highest correlating window size below 1MB and of 1MB or larger. The residuals of the linear model are saved as the corrected LogR.

Inclusion of SVs during segmentation increases the accuracy of the segmentation and therefore the copy number calls. We first create SV segments by sorting the breakpoints by chromosome and position. Then, for every SV segment, PCF is run separately to obtain the final segmentation. After copy number has been established, we merge adjacent segments with equal copy number states. This step removes SV supported segments that do not constitute a copy number change (inversions).

##### 1.2.4 cloneHD

Ignacio Vázquez García, Ville Mustonen

We used the cloneHD algorithm for copy number calling (Fischer et al., 2014). Briefly, cloneHD implements a Hidden Markov Model (HMM) that describes the (hidden) copy number state of the sample. The hidden states emit observations that correspond to the read depth and B-allele frequency counts (BAF) of each sample. The emission models implemented in cloneHD are Poisson for read depth and Binomial for BAF, and their over-dispersed counterparts. Here we used cloneHD in read depth + BAF mode and exclusively the Poisson and Binomial emission models.

The first step in the cloneHD pipeline is the filterHD algorithm, which is a continuous state space HMM with a jump-diffusion state transition model. The diffusion component is useful to allow for the hidden state to change smoothly and models sources of (non-biological) bias. The jump component corresponds to a discrete change in the hidden state above the noise level, such as a copy number gain or loss. The result of filterHD is a posterior probability of the hidden state, e.g., the posterior read depth (both for the tumor and its matched normal) or the posterior B-allele frequency for BAF, and per-locus jump probabilities. Importantly, filterHD does not seek to explain the subclonal structure in the data and is a generic algorithm for fuzzy segmentation. It plays a similar role to the segmentation step carried out by most copy number callers.

To find out a biologically interpretable model of the data we used cloneHD. This was done by applying cloneHD to both read depth and BAF data of each sample, the per-locus jump probability for these, and the mean of the posterior read depth for read depth of the matched normal sample (to correct for bias). The HMM implemented in cloneHD aims to explain the data as a mixture of copy number states weighted by the corresponding cellular fractions of normal and cancerous cells, while allowing for the hidden state to change using the per-locus jump probability. Copy number calling using single samples from a tumor suffers from degenerate solutions that can be difficult to resolve. Most importantly, a correct baseline ploidy of the sample has to be inferred. cloneHD allows the user to specify a penalty for leaving a normal baseline (diploid) copy number. A penalty value of 1.0 does not penalize leaving the normal baseline. Without this penalty, higher copy number baselines are typically preferred over lower ones. Biologically, this would mean a higher number of inferred whole genome duplications. By default, we used 0.95 for this penalty. This setting was used to provide cloneHD copy number calls for the consensus breakpoint calling. After consensus breakpoints were available we used them to obtain per-locus jump probabilities, allowing for transitions only at these set locations. We then executed cloneHD as before, using the baseline penalty values 0.80, 0.95 and 0.99. Based on the consensus average ploidy for each sample we then selected the closest cloneHD solution. We note that for most samples all three solutions are essentially the same, however, for a subset the penalty is decisive whether to call a whole genome duplication or not.

##### 1.2.5 JaBbA

Xiaotong Yao, Steven Schumacher, Rameen Beroukhim, Marcin Imielinski

We used Junction Balance Analysis (JaBbA) to integrate paired-end and read depth signals and infer copy numbers on genomic intervals and rearrangement junctions.

JaBbA exploits the principle that copy number alterations (CNA) always result from rearrangements or whole chromosome gains / losses, previously explored in several publications on germline and cancer genome structural variant analysis (Li et al., 2016; Medvedev et al., 2010; Oesper et al., 2012). This results in a junction balance constraint (JBC), namely that the copy number of every genomic segment is consistent with its neighbors. Applied to a genomic graph whose nodes represent (+ and – strands of) genomic segments and edges represent alternate and reference junctions (corresponding to 3-5' phospho-diester bonds), the JBC forces the copy number of every node to equal the sum of the copy numbers of all incoming (similarly, outgoing) edges.

To fit this model to data, we formulate a mixed integer quadratic program (MIQP) to match node copy numbers with estimates of read depth measurements parameterized as posterior means and variances at $n$ genomic segments. The summed deviation of the model fits (i.e. segment copy numbers) from the observed data (mean estimate on segment read depths) is captured in a weighted sum of squares quadratic objective function that incorporates purity, ploidy, and inverse variances as weights. To allow for missing junctions, which frequently result from rearrangements occurring at hard to map genomic regions (e.g. centromeres) and force local violations of junction balance, we add a slack parameter to the JBC. These slack parameters represent "loose ends" in the structural variation model, which can either represent missing junctions or "biological loose ends" that could correspond to neo-telomeres in the cancer karyotype. We then add a linear slack penalty to the objective function, which penalizes the number of slack (loose end copies). We provide a user-defined hyper-parameter, which balances the contribution of (linear) slack and quadratic (read-depth) penalties to the analysis. Model fitting yields estimated integer value copy numbers on graph nodes, edges, and loose ends and numeric value purity / ploidy.

For the PCAWG consensus copy number analysis, we applied JaBbA in two rounds, in concordance with the other copy number methods. In practice, JaBbA begins with a .bam file, a junction call set (e.g. a BND VCF file, a .bedpe), and (optionally) a preliminary segmentation and purity / ploidy input. We compute high density binned read depth (200bp bins) by calculating tumor / normal ratios of GC and mappability corrected coverage of "proper" read pair centroids in the .bam file (fragCounter). In the absence of a preliminary segmentation (pre-consensus run), we use circular binary segmentation (CBS) to segment high-density binned coverage (~15M intervals) into a lower dimensional collection (e.g. 10-100K regions) of regions of constant copy number. In the subsequent "consensus" run, we use the "initial copy number consensus" as the preliminary segmentation. For the junction input, we used the full PCAWG-6 call set during the pre-consensus run. In subsequent runs, we used a strict intersection of this junction set with the end points of consensus PCAWG-11 segmentation.

The union of the endpoints in this segmentation and the breakpoints in the junction call set induce a partition of the reference genome, from which we build a genomic graph of $n$ segments and $m$ edges. We use the high-density coverage to compute posterior means and variances on fragment density at each of these $n$ segments. We solve a preliminary least squares problem (corresponding to our full MIQP objective function, but without JBC, i.e. with zero slack penalty) to infer an affine transformation between read depth and integer copy number, and hence purity and ploidy. This least squares inference implements principles described in by Van Loo and colleagues (Van Loo et al., 2010). We find that in practice, the JBC does not alter purity and ploidy estimation, and dividing the computation into these two phases dramatically improves convergence and speed of the MIQP. Following purity and ploidy inference, application of the full MIQP using CPLEX (https://www.ibm.com/software/commerce/optimization/cplex-optimizer/) fits $n$ nodes and $m$ edges in the graph with integer copy numbers. After fitting integer (total) copy numbers, JaBbA uses allelic read counts at germline heterozygotic sites (obtained via samtools pileup at HapMap v3 sites) to infer likely allelic copy numbers at genomic segments. See the pipeline on Fig. 4.

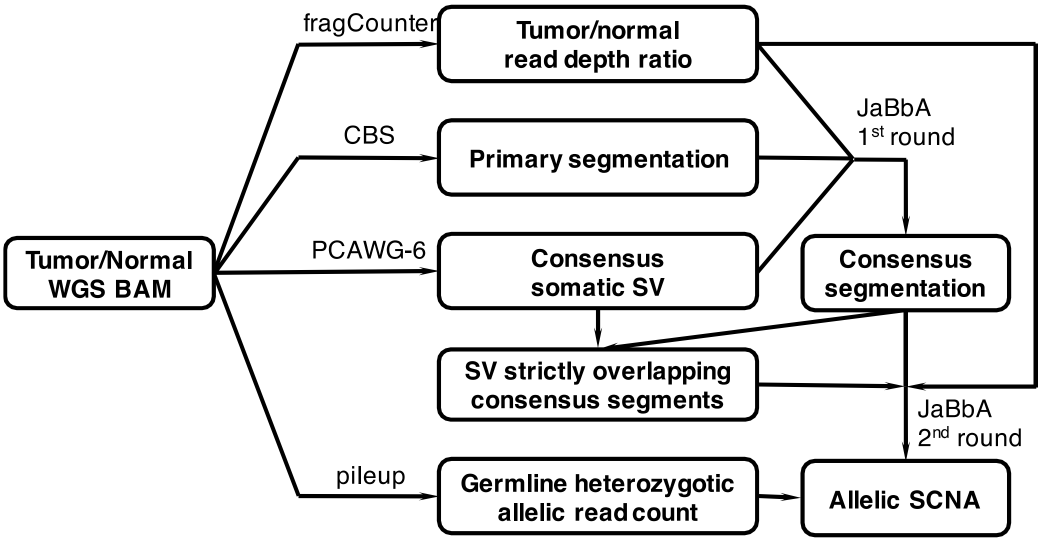


*Fig. 4 The full JaBbA pipeline depicted graphically.*

#### 1.2.6 Sclust

Martin Peifer, Yupeng Cun, Tsun-Po Yang

The copy number module of Sclust has extensively been used in our recent large-scale sequencing efforts of small cell lung cancer and neuroblastoma (George et al., 2015; Peifer et al., 2015). In total, Sclust performs copy number segmentation and determines purity, ploidy, and allele-specific copy number (both clonal and subclonal). As input, the method requires read counts from tumor and matched normal sample in a sufficiently partitioned genome. Within the ICGC pan-cancer analysis, we chose a partitioning into non-overlapping 1kb windows. The read counts are then used to generate GC-corrected read ratios between the tumor and the normal. Secondly, Sclust requires base counts of common single nucleotide polymorphisms (SNPs) in both the tumor and the matched normal sample. This data is used to compute B allele frequencies at all heterozygous sites of the normal. Given this as input data, the algorithm first preforms segmentation based on the read ratios by detecting significant jumps in the data. Purity and ploidy estimates are then computed using a conditional maximum likelihood approach. In particular, the likelihood of B allele frequencies is optimized to obtain purity estimates for a fixed expected ploidy. This yields purity estimates in function of the expected ploidy, which is plugged into the likelihood of read ratios and optimized in order to obtain an estimate of the expected ploidy. Here, the search domain can be adjusted to select a more suitable solution in case of whole genome duplications. Along the optimization, allele-specific copy numbers are computed. To derive subclonal copy number changes, we test which of the B allele frequencies disagree with the model predictions (here a significance threshold of 1% is chosen). For all segments assigned to be subclonal, we select the best combination of copy number states that are just one copy apart. Finally, ploidy estimates are computed directly from the determined copy number states.

### 1.3 Copy number consensus approach

#### 1.3.1 Consensus copy number procedure

ICGC PCAWG relied on a consensus strategy for SNVs, SVs, and indels, as calls for each on which different algorithms agreed were understood to be high-confidence predictions. For copy number calls, we relied on a similar consensus approach, which combined results from six individual copy number callers: ABSOLUTE, ACEseq, Battenberg, CloneHD, JaBbA and Sclust.

Each copy number caller uses a two-step process, first segmenting the genome into regions assumed to have constant copy number state, then determining the clonal and subclonal copy number states of each segment. Disagreement amongst copy number callers arises primarily from two factors: differences in genome segmentation, and uncertainty concerning whether a whole-genome duplication (WGD) occurred. Thus, our consensus strategy resolved both factors for each sample, allowing us to determine a consensus copy number state for much of the genome across samples.

#### 1.3.2 Copy-number-calling methods differ in genome segmentation

Jeff Wintersinger, Quaid D. Morris

Copy number callers segment a sample's genome into regions assumed to have constant copy number. To delineate these segments, they find breakpoints at which the boundaries between segments where copy number state changes. Once these breakpoints are established, callers then determine the mixture of copy number states within each segment, encompassing the number of major allele copies, the number of minor allele copies, and the proportion of cells with that state.

Copy number methods differed substantially in the number of breakpoints they defined through their genome segmentations (Fig. 5), with some methods calling an order of magnitude more breakpoints than others. Broadly speaking, these can be broken into two classes: "liberal" methods (ACEseq and cloneHD) called, on average, a great many more breakpoints than "conservative" methods (ABSOLUTE, Battenberg, JaBbA, and Sclust). To resolve this disagreement between different algorithms concerning the proper genome segmentation, we established a set of consensus breakpoints, taking into account this “liberal” vs. “conservative” distinction. All methods subsequently used that consensus segmentation in determining copy number states.

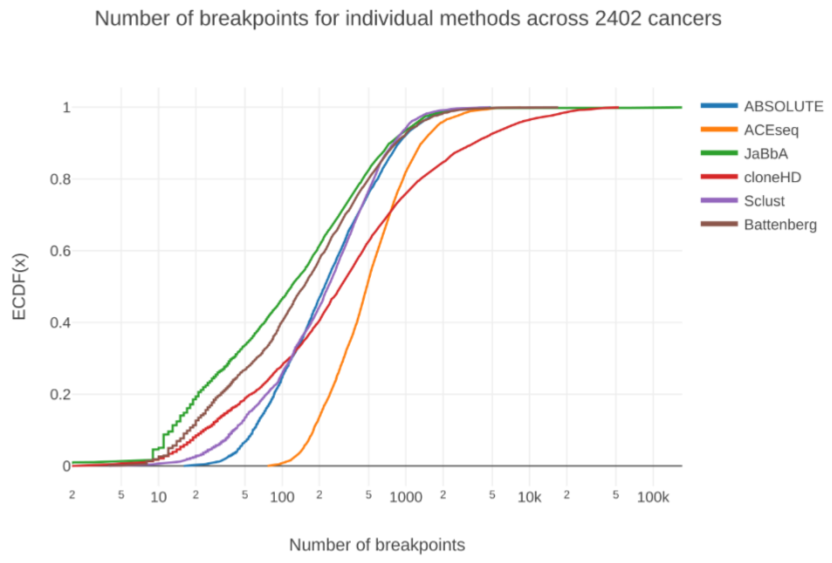


*Fig. 5 Cumulative distribution of the number of samples with the indicated number of breakpoints predicted for each tumor by the six copy-number-calling methods. ACEseq and cloneHD (the "liberal" methods) often predicted an order of magnitude more breakpoints per tumor than ABSOLUTE, JaBbA, Sclust, and Battenberg (the "conservative" methods). While the four conservative methods characterized only approximately 7% of cancers as having more than 1000 breakpoints, ACEseq and cloneHD found 17% and 24%, respectively, crossing this threshold.*

#### 1.3.3 Method for determining consensus segment breakpoints

Jeff Wintersinger, Quaid D. Morris

As the six copy-number-calling methods differed substantially in the segmentation they calculated, we increased agreement in our consensus copy number calls by deciding upon a set of consensus breakpoints for each genome. We developed a consensus strategy that favored "true positive" breakpoints at the potential cost of increasing "false negatives" to create a complete set of breakpoints. Orthogonal evidence of copy number breakpoints from structural variants was used to quantify the "true positive" and "false negative" rate of our consensus approach (further detailed below). Copy number methods were required to take the consensus segmentation as input, but were permitted to merge adjacent segments they judged as having the same copy number. They were however not allowed to introduce additional breakpoints that would create new segments. The cost of introducing spurious breakpoints was therefore less than that of missing breakpoints where the underlying copy number state did indeed change.

The algorithm we developed for determining consensus breakpoints draws on the insight that regions between adjacent segments indicate a method's uncertainty in precisely where the breakpoint delineating change in copy number state should lie. The segmentation released by each method consists of a set of regions defined by the genomic loci $S_i$ and $E_i$, indicating the start and end of each region, with the interval ($S_i$, $E_i$) representing a region of constant copy number. On a given chromosome, however, we need not have each region immediately following its predecessor such that there is no gap, which would imply $S_i = E_{i-1} + 1$. Instead, the region ($E_{i-1}$, $S_i$) has undefined copy number—the segmentation method inferred that CN status changed at some point within this interval, but could not pinpoint the location because of the noisy signal.

Our consensus breakpoint algorithm leverages this information concerning uncertainty (Fig. 6).

1. For each copy number segmentation method $M_s$, take each reported segment ($S_i$, $E_i$), and generate an interval spanning ($E_i - \delta$, $S_{i+1} + \delta$). This interval represents a plausible region over which a breakpoint may lie according to $M_s$, permitting the breakpoint to move $\delta$ bases upstream or downstream beyond the reported boundaries. Here, we set $\delta$ = 50 kb, which we selected after comparing the breakpoints generated by a range of $\delta$ values to the underlying read depth and B allele frequency signals in the data. $\delta$ = 50 kb achieved a reasonable balance between false-positive consensus breakpoints (when $\delta$ was too large) and false-negative consensus breakpoints (when $\delta$ was too small).
2. Compute the intersection of intervals between each method. Scanning from the start of the chromosome, find the first intersection $I_s$ supported by the threshold method set $T$. Here, we defined $T$ to be any combination of at least three of the six copy number methods, or any combination of two of the "conservative" methods (i.e., ABSOLUTE, Battenberg, JaBbA, and Sclust). We called this strategy *any3_any2_conservative*. This avoided calling consensus breakpoints supported by only the two "liberal" methods (ACEseq and cloneHD), while correcting false-negative cases where a breakpoint was supported by only two of the input methods despite clear evidence in the underlying data. Relative to *any3*, a stricter criterion requiring that at least three of six methods support a breakpoint, *any3_any2_conservative* added only a small number of breakpoints (Fig. 7).
3. Select the start and end interval loci from each method $m$ falling within the interval, corresponding to $S_m$ and $E_m$, respectively. Score each locus according to the size of the associated gap $G_m = \text{rank}(S_m - E_m)$, where $G_m$ corresponds to the rank in the empirical cumulative distribution of all gaps generated by the given method. Thus, if a method assigns a large gap between two segments relative to the other segments it generates, its uncertainty in breakpoint placement is understood to be relatively large; conversely, a relatively small gap indicates high certainty. Select the breakpoint from within the intersection that has the smallest $G_m$ value. In the case that two breakpoints in the intersection have the same $G_m$ (which occurs, e.g., because both $E_m$ and $S_m$ fell in the intersection), arbitrarily prefer end loci to start loci. Record this as the single consensus breakpoint associated with this intersection. Otherwise, in the rare case that no input start and end loci fall in the intersection, report the upstream-most end of the intersection as the consensus breakpoint. Such cases arise when only the $\delta$ bases padding each input segment intersect, meaning that the intersection as a whole is relatively small, and that either end of the intersection can be taken as a reasonable representation of a breakpoint's position.
4. Remove all intervals that contributed to the intersection. Return to step 2. Repeat until no intersections passing the threshold remain on the chromosome.
5. Add PCAWG consensus structural variants to the consensus breakpoint set. To do so, find all consensus breakpoints within 100 kb of a consensus SV. Replace the consensus BP with the consensus SV, as the SV presumably represents the same mutational event, but with greater precision concerning position. For any

SVs lacking a consensus BP within 100 kb, add the SV as an additional consensus breakpoint.

6. Add breakpoints at centromeres and telomeres as necessary, as copy number states cannot be called across these boundaries. Use the chromosome lengths and centromere start and end locations reported in the hg19 human reference genome. If any centromere start or end lacks a consensus breakpoint within 1 Mb, add an additional consensus BP at that location. If a consensus breakpoint occurs within the centromere, move it to the start or end of the centromere, according to whichever point is closer. Likewise, if no breakpoint occurs within 1 Mb of a chromosome start or end position (representing telomere locations), add an additional breakpoint at the chromosome start or end.

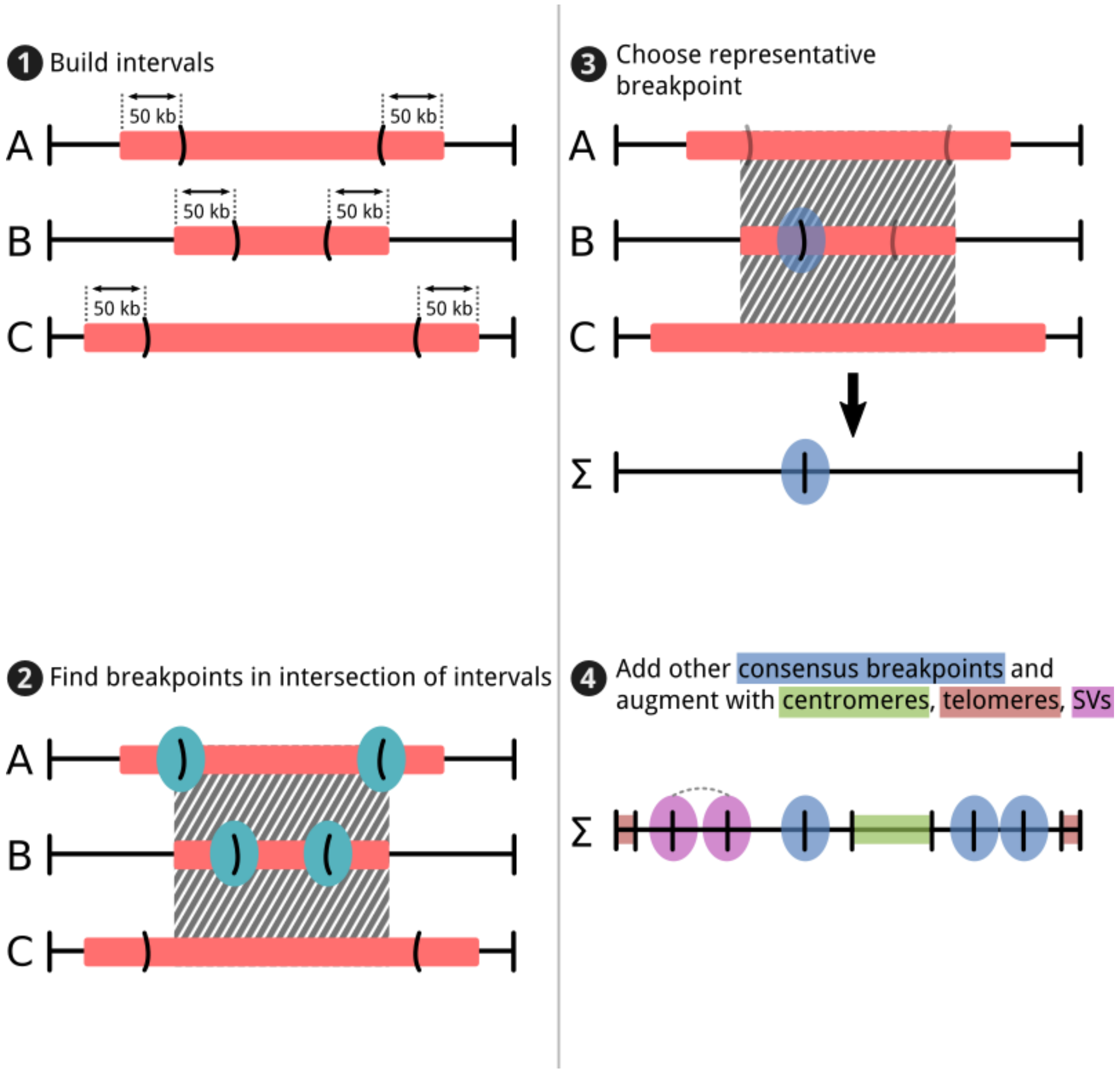


*Fig. 6 Individual genome segmentation methods A, B, and C report segments of constant copy number. In this example, we determine the consensus segmentation Σ, requiring that all three methods A, B, and C contribute to the intersection that yields consensus breakpoints. Each segment from methods A, B, and C is composed of a start locus $S_i$ and end locus $E_i$. Depicted here for each method is the end point of one segment, $E_i$, and the start point of the next segment, $S_{i+1}$. Note that the methods may differ in the copy number state they assign to each segment, as the consensus breakpoint method is concerned only with where segments occur, not what the status of each segment is. By proceeding through steps one to three, the consensus breakpoint algorithm selects a single breakpoint supported by the input segments from the individual methods. In step four, the breakpoints supported by the individual*

*segmentation methods are refined and augmented using the PCAWG consensus structural variants, as well as knowledge of where centromeres and telomeres occur on each chromosome.*

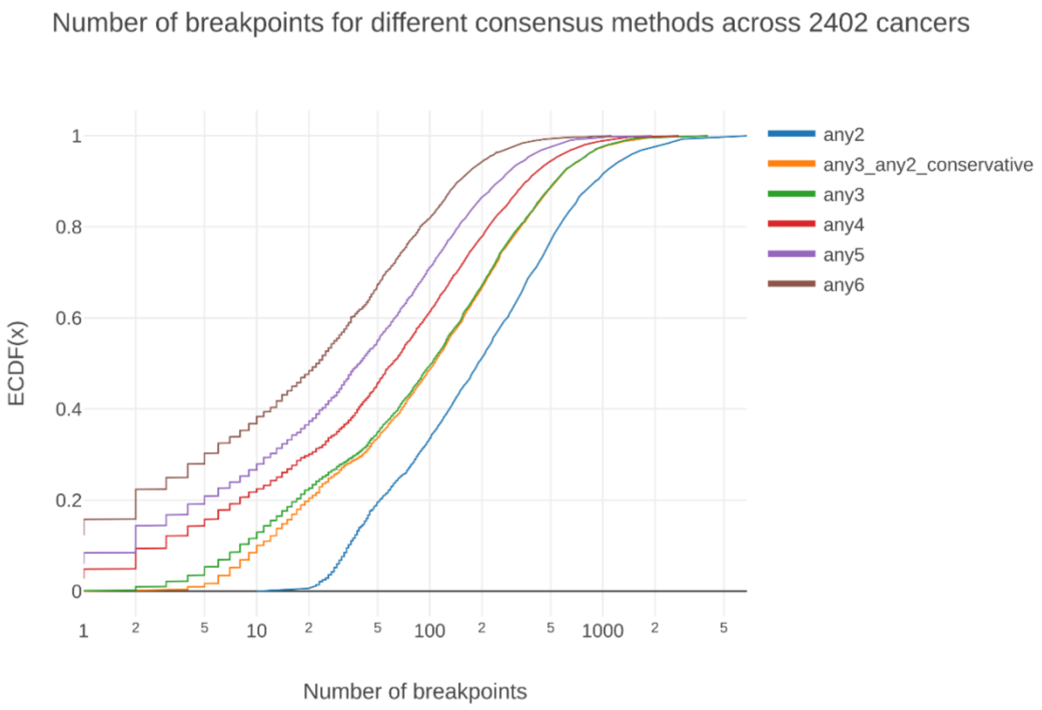


*Fig. 7 Cumulative distribution of the number of samples with the indicated number of breakpoints produced for each cancer by different consensus strategies. Given the six copy-number-calling methods, the anyN strategies required N of six methods to agree on a breakpoint's placement to establish a consensus breakpoint at that location. The strategy we selected, any3_any2_conservative, required agreement between any three methods, or agreement between any two of the four "conservative" methods (i.e., ABSOLUTE, Battenberg, JaBbA, and Sclust). This avoided calling consensus breakpoints supported by only the two "liberal" methods (ACEseq and cloneHD). Relative to any3, the any3_any2_conservative strategy introduced only a few extra breakpoints, but corrected false negatives where we failed to obtain support for a breakpoint from three methods despite clear evidence of its existence in the underlying data.*

#### Most consensus breakpoints obtain support from SVs

Every copy number event should generate associated structural variants (barring whole chromosome events or breakpoints that fall within regions where difficulties in the alignment of sequencing reads hinder SV calling, like centromeres). As such, each breakpoint should have associated with it a nearby structural variant. On a per-tumor basis, we compared the number of consensus breakpoints with support from the PCAWG consensus SVs, to the number of consensus breakpoints lacking such support (Fig. 8). Under the *any3_any2_conservative* strategy, 80% of cancers had more SV-supported than SV-unsupported breakpoints, which helped validate our consensus threshold. In examining the proportion of breakpoints with SV support as a function of number of breakpoints, we found that the average cancer received SV support for 77% of its breakpoints (Fig. 9A), with a slight positive correlation between total number of breakpoints and the proportion finding SV support. Furthermore, in each tumor, 83% of consensus SVs supported nearby consensus breakpoints (Fig. 9B), with no correlation between number of SVs and the fraction that supported breakpoints.

In theory, all non-centromeric, non-telomeric copy number events should have associated structural variants, but not all structural variants should correspond to copy number changes (e.g., balanced translocations do not affect copy number). Consequently, finding on average SV support for 77% of our consensus breakpoints assured us that our consensus breakpoints were largely recapitulating the breakpoints found by orthogonal structural variant detection algorithms, increasing confidence in our results. Manual inspection of consensus breakpoints without associated consensus SVs suggested that there was sufficient evidence in the underlying B allele frequency and read depth signals to call the breakpoints, implying that simply taking the set of consensus SVs as our breakpoints would have missed legitimate copy number events. Given our preference for false positives over false negatives, the non-SV-supported breakpoints were a worthwhile addition to our breakpoint set, even if some were false.

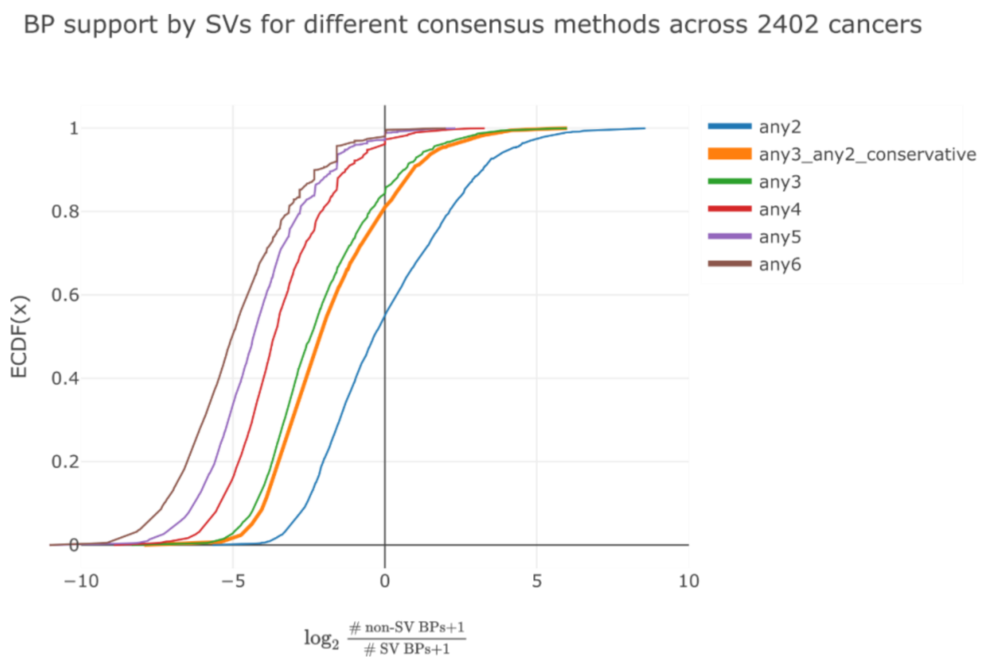


*Fig. 8 Cumulative distribution across cancers of the log-ratio of the number of breakpoints without support from SVs, to the number of breakpoints with SV support. As virtually all copy number events should generate associated structural variants, most consensus breakpoints should be able to find a nearby supporting SV. Under the any3_any2_conservative consensus breakpoint strategy, only 20% of cancers had more unsupported breakpoints than SV-supported breakpoints (i.e., a log ratio greater than zero).*

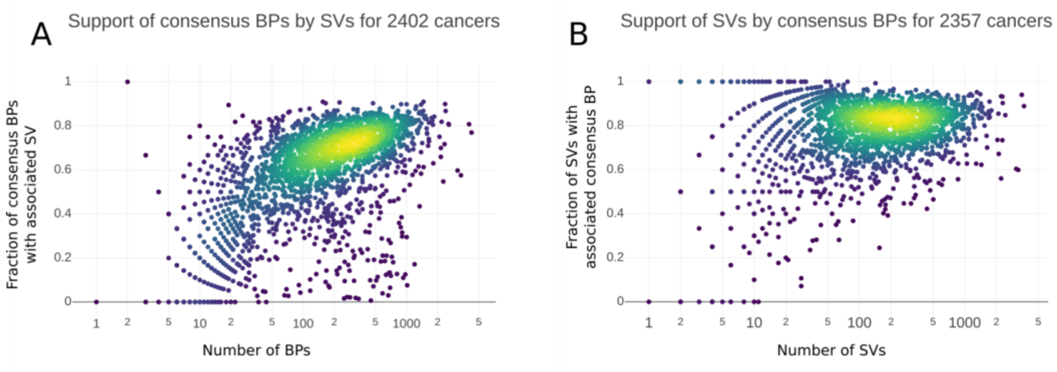


*Fig. 9* ***A)*** *Support of consensus breakpoints (BPs) from SVs as a function of number of BPs. Color indicates density. As the number of BPs increased, so too did the fraction of BPs that found a nearby supporting SV. The mean fraction of BPs with SV support was 77%.* ***B)*** *Support of SVs from consensus BPs as a function of number of SVs. Color indicates density. There was no correlation between the number of SVs and the fraction with support from BPs. The mean fraction of SVs with BP support was 83%.*

#### 1.3.4 Resolving ploidy disagreement between copy number callers

Jeff Wintersinger, Stefan Dentro, Ignaty Leshchiner, Kortine Kleinheinz

After the six copy number callers produced copy number profiles on the provided consensus segmentation, we focused on disagreement between the callers on ploidy calls. An automated procedure was used to identify these tumors, which were subsequently subjected to a rigorous review procedure with the aim to resolve the discrepancy.

##### Finding outlier profiles

Jeff Wintersinger

CNA callers often disagreed on whether cancers had undergone whole-genome duplications (WGDs). To determine which cancers suffered from this disagreement, we ran all CNA callers across the full dataset, then declared each cancer “WGD-certain” if all callers agreed on whether it was diploid or tetraploid, and “WGD-uncertain” otherwise. We performed this characterization as follows:

1. For each caller, compute the proportion of the genome with a clonal diploid copy-number state of (major allele, minor allele) = (1, 1). Also compute the proportion in clonal tetraploid (2, 2). If either proportion exceeded 20% of the genome, declare the caller’s result on that cancer “diploid” or “tetraploid”, respectively. Otherwise, if neither state accounted for at least 20% of the genome, report the caller’s result as “unknown”.
2. If all callers agreed on “diploid” or “tetraploid”, declare the cancer as “WGD-certain”, meaning that the callers agree it is diploid (no WGD occurred) or tetraploid (a WGD occurred).
3. Otherwise, if any callers disagree with the others on “diploid” or “tetraploid”, or if any caller reports “unknown”, declare the cancer as “WGD-uncertain”.

After this procedure, 461 of 2778 cancers were deemed WGD-uncertain.

A preliminary consensus copy number procedure revealed an additional 315 tumors with agreement on less than 20% of the genome between the CNA callers. These 315 cases were also reviewed.

#### Resolving whole genome duplication uncertainty

Ignaty Leshchiner, Stefan Dentro, Kortine Kleinheinz

The above described copy number reconstruction methods differ in how copy number is estimated and perform different pre-processing steps. It was therefore expected that the methods could deviate on the called profiles. The samples identified through the above procedure have been put through a review by a panel of experts. Initially to understand where the discrepancy lay between the profiles, and later to resolve the differences. The expert panel consisted of three core and five alternating members and sat down for four afternoons. Each member prepared a figure per sample with all possibly interesting information (raw data, SNV multiplicity states, subclonal architecture calls, etc.). A central figure (see Fig. 10 and Fig. 11 for examples) was used to feed the discussion that contained: Copy number profiles from all methods and raw BAF, copy ratio (LogR) and multiplicity values from ABSOLUTE. During the review, a sample was marked as *WGD* or *no_WGD*, and it was only marked on unanimous agreement amongst the panel. When the panel could not agree the sample was marked as *unknown*. The panel operated on the basis that sufficient evidence of a genome doubling is required to mark a tumor as *WGD*.

Key identifiers of a missed whole genome doubling that were considered: Subclonal copy number called at 0.5 CCF (these segments become clonal when doubling), a SNV cluster at 0.5 CCF (these mutations become clonal when doubling the ploidy), there should be a clonal SNV cluster at around 1.0 CCF and no superclonal clusters beyond 1.0 CCF, while very large homozygous deletions in general should not occur (whole chromosome(s), or whole chromosome arm(s), in general we worked with a threshold of 20Mb, which is often just visible on whole genome figures). Finally, when the tumor is called as genome doubled one expects some SNVs on one chromosome copy, except in the unlikely scenario that the whole genome duplication was the last event to occur in the tumor before resection and it became clonal without the tumor acquiring further SNVs.

Over time, we observed recurrent scenario's, which allowed for quick identification of the outlier. For example, one method did not allow for a purity below 0.3. It would call a higher purity, pushing the copy number profile upwards and leaving a purity/ploidy discrepancy that does not allow clean inference of the subclonal architecture inference from SNV data. Another example are samples with very heavy wave artefacts in the coverage. In this case, some methods would over-segment the genome to fit the noise, leading to a discrepant ploidy that our automated approach could not resolve (see Fig. 11). Methods have been adapted since the review to address the scenarios uncovered.

The panel did not agree on the WGD status of 38 cases, due to the high complexity of their copy number profiles. All segments of these genomes have been marked with level *i* accordingly (see further).

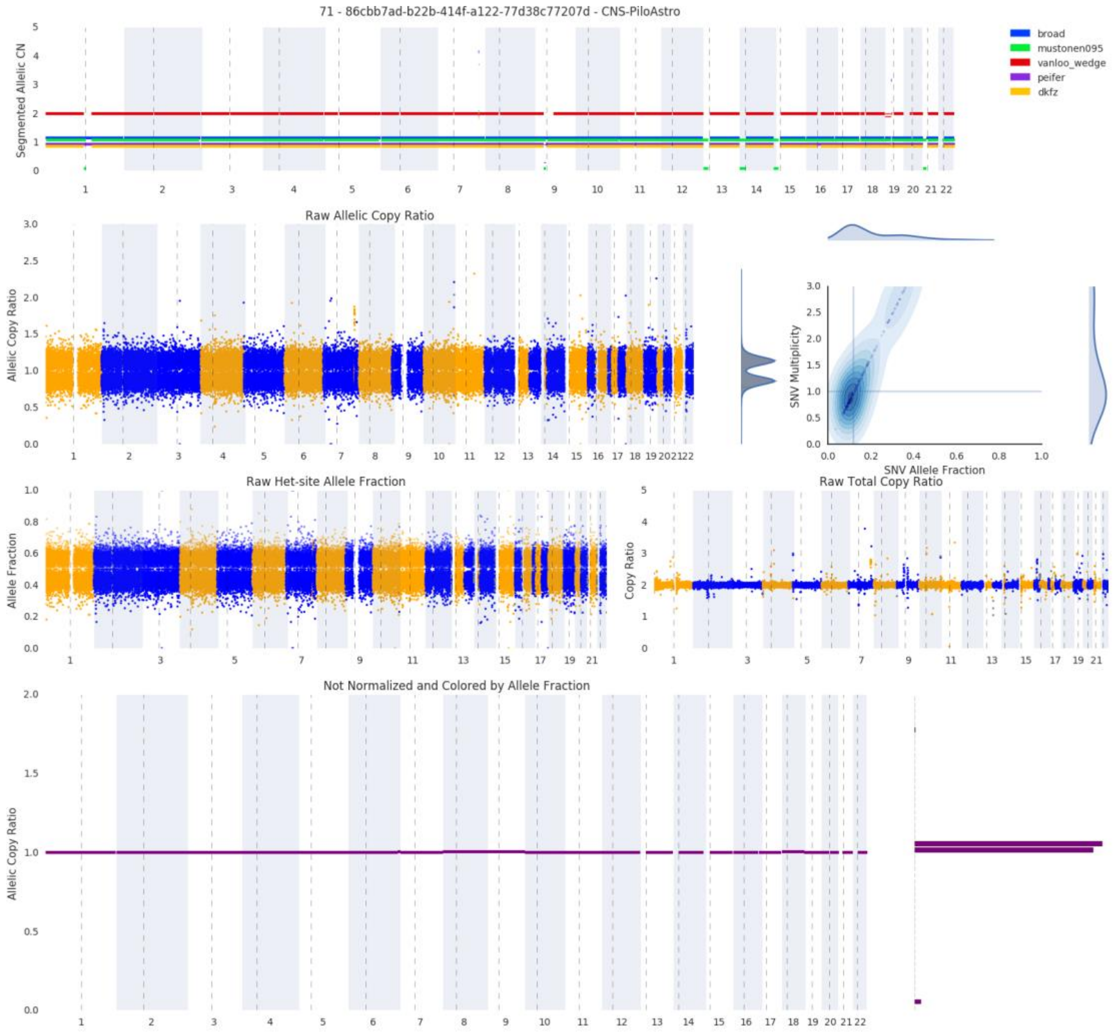


*Fig. 10 Copy number review figure for sample SA414133. The figure contains copy number profiles for five methods (top), raw allele ratio's for SNPs and multiplicity values (second row left and right respectively), raw B allele frequencies and copy ratio's for SNPs (third row left and right) and normalized allele ratios (bottom). In this case the raw data shows there are no copy number alterations of note. Battenberg however (red copy number profile) calls a whole genome duplication. Adding the duplication does not allow for an increase in the proportion of the genome called with clonal copy number. In this scenario Battenberg was considered the outlier and the profile is flagged as no_WGD. This figure shows copy number profiles on the methods' own segmentation.*

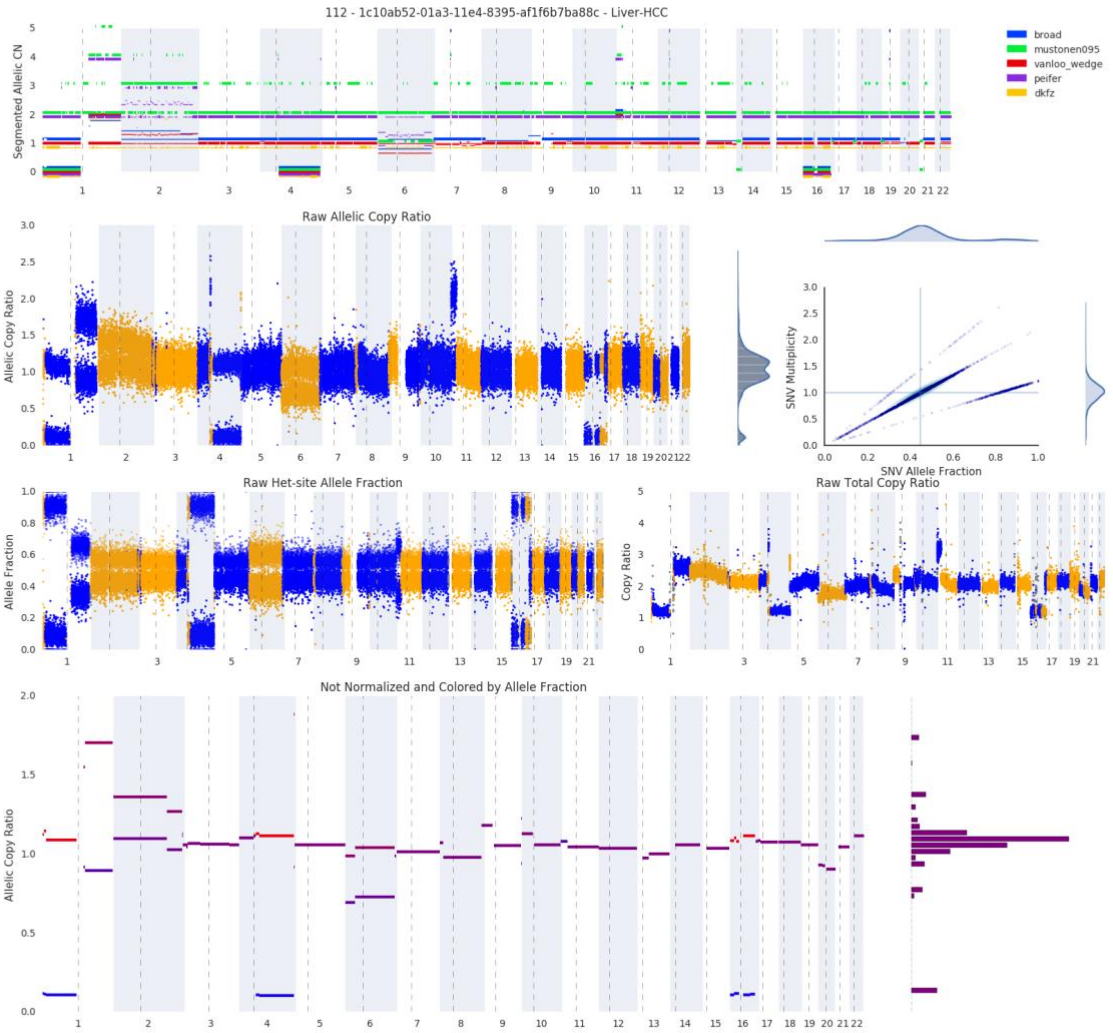


*Fig. 11 Example of copy number profile where noise affects the fit and causes disagreement on the ploidy. The copy number profiles called by cloneHD (green), Sclust (purple) and ACEseq (yellow) contain large numbers of small segments, compared to ABSOLUTE (blue) and Battenberg (red). The methods affected by the noise interpret it as signal, which cause it to fit the segments with a higher or lower copy number state, resulting in a ploidy discrepancy. This figure shows copy number profiles on the methods' own segmentation. The call for this sample (SA529805) is no_WGD.*

#### 1.3.5 Consensus copy number

Stefan Dentro, David Wedge, Peter Van Loo

With a consensus segmentation established and whole genome duplication uncertainty resolved we could build the consensus copy number profiles. We aimed for every profile to contain a call for every segment where there are reliable calls. To do so we first identified 6 ways of extracting agreement between the CNA callers:

a. All methods agree on a clonal copy number call (both major and minor alleles).
b. A single method disagrees on the copy number state of a single segment, leaving the call out creates agreement.
c. A single method disagrees on the ploidy of a sample, leaving the profile out creates agreement.

d. The majority of available methods agree on clonal copy number.
e. Complete or leave-one-out agreement may be achieved by rounding subclonal copy number.
f. Majority vote may be achieved after rounding subclonal copy numbers.

For each sample, every segment goes through the list starting at *a*, until agreement is reached. On average, we obtain consensus on 93% (median 95%) of the genome after reaching level *f*. The segments that remain without a consensus call go through a second approach that is designed to find a call from a single method.
We first calculate, for every CNA method, what proportion of the consensus profile it agrees with after reaching level *f*. This allows ranking of the methods, where an overruled profile due to ploidy is not included (see filtering below). We then devised the following additional levels:

g. Take the call from the best method. If there is consensus for the copy number state of one of the alleles we require the best method to agree with it (see rounding below).
h. Take the call from another method, iterating from the best to the worst performing method.

A special level was added to distinguish samples where the expert panel did not reach consensus on the ploidy of a sample. These cases went through the full consensus procedure, but afterwards all their segments were re-marked as level *i*.
We then devised a star rating system that denotes the amount of confidence in each of the calls. Levels *a*, *b* and *c* are the strictest and require all methods but one to agree, at the least. These segments are therefore assigned 3 stars. Segments for which a majority of the methods agree on either clonal or rounded clonal copy number are assigned 2 stars (levels *d*, *e*, *f*). The remaining levels (*g*, *h*, *i*) receive 1 star to denote the lowest confidence.

#### Rounding subclonal copy number

Subclonal copy number is reported in three different ways across the 6 methods. ABSOLUTE reports up to 3 different copy number states (a combination of a major and a minor allele) per segment, of which 1 is termed the ancestral state. Battenberg and Sclust report subclonal copy number as a mixture of two states, while ACEseq returns a single non-integer state (i.e. a mixture). Both cloneHD and JaBbA provided clonal calls only.

When subclonal copy number is reported as individual states found in proportions of cells (ABSOLUTE, Battenberg and Sclust), we selected the reported states individually. For ABSOLUTE that corresponds to selection of the copy number states corresponding to the ancestral state, the highest CCF state and the lowest CCF state separately. For Battenberg and Sclust that meant selecting the states from the highest CCF and lowest CCF calls separately. As ACEseq reports two non-integer states we opted for rounding both alleles up and down.

To create a consensus call, we first obtain an inventory of the available copy number states across all possible rounded states, including the clonal calls from cloneHD and JaBbA. If there is a major/minor allele combination that satisfies the minimum number of methods criterion (either leave-one-out or majority vote) we select that state as the consensus.

If no agreement is reached we attempt to establish consensus by voting for the major and minor allele separately. An allele is accepted if it passes the minimum number of methods threshold. In some cases this leads to consensus on one of the alleles. The state of that allele is saved and fed into levels *g* and *h*, where a call is selected where one of the alleles agrees with the established consensus allele.

#### Chromosomes X and Y

Fewer methods report on X and Y chromosomes:

- X: ABSOLUTE, Battenberg (females), ACEseq, cloneHD and JaBbA
- Y: ABSOLUTE, ACEseq and JaBbA

The number of methods required to agree for the separate levels is adjusted accordingly.

#### Consensus purity

For consensus purity, we have calls from the 6 CNA methods and a number of SNV-based methods: CliP, CTPsingle, PhyloWGS, cloneHD (on SNVs) and Ccube (discussed further). Outlier calls are first removed for CNA and SNV methods separately (see filtering below). For each sample, we establish a density over the combined data. Analogous to taking the mode, we select the call that is closest to the highest peak in the density as the consensus.

There is a larger discrepancy in purity calls from CNA methods on samples with few copy number alterations. For tumors where less than 8% of the genome is altered, we therefore calculate the density over the calls from SNV based methods only.

Finally, the median absolute deviation is calculated to capture the amount of agreement between the calls and is used as a measure of confidence.

#### Filtering

After the expert panel review of ploidy-uncertain cases, a reference ploidy can be obtained for almost all samples. The methods either all agreed on large portions of the genome or certain ploidy estimates were overruled by the expert panel. We therefore calculated a reference ploidy that serves to overrule calls from individual CNA callers. To allow for larger variations on high ploidy profiles we set the threshold at 0.25 times the reference ploidy. If a profile differed by more than this threshold, it was automatically overruled and excluded from the procedure for both copy number and purity.

Further filtering on the purity calls was performed to remove outliers. A purity call was filtered out if it differed from all other non-ploidy-overruled purity calls by more than 0.2. This method was applied separately to SNV-based purity values.

We also excluded calls on complex regions by removing segments that start at the first base of the acrocentric chromosomes 13, 14, 15 and 21.

##### 1.3.6 Subclonal copy number consensus

Ignaty Leshchiner, Daniel Rosebrock

To produce subclonal consensus copy number, three callers that had the highest concordance in the subclonal segments were chosen: ABSOLUTE, Battenberg, and Sclust. For 3-star segments, consensus copy number was kept. For 1- and 2-star segments, if 2 out of 3 callers agreed on the allelic copy number state, then this copy number was used. If at least 2 callers were in agreement of subclonality of one allele's copy number state, then the segment was called as subclonal. If ABSOLUTE produced one of the subclonal calls, then the consensus copy number value reported by ABSOLUTE was used (due to its internal Dirichlet Process clustering of states) if downstream merging of copy number segments was required (e.g. for calling arm level events). Segments for which all callers disagreed on copy number state were called as NA.

#### 1.4 Validation

##### 1.4.1 Comparison of purity against TCGA

Kerstin Haase

As a means to validate our consensus inferred purity values, we compared them to the published purities from (Aran et al., 2015). The authors provide consensus purities for 9,364 samples across 21 cancer types from the TCGA cohort. They analyzed the samples with four orthogonal methods and have integrated them into a consensus. The different utilized approaches include expression profiles of a panel of immune and stromal genes (ESTIMATE), somatic copy number data (ABSOLUTE), leukocyte unmethylation (LUMP) and image analysis by heaematoxylin and eosin staining (H&E staining). Consensus purity is determined by normalizing all methods to give them equal means and standard deviations (CPE).

Even though the methods applied by Aran et al. (2015) are quite different to our consensus approach using WGS, our consensus has a high correlation with the one reported by them (R=0.76) (**Supp. Fig. 1**).

### 2. Subclonal architecture methods

#### 2.1 Description of reconstruction methods

##### 2.1.1 BayClone-C

Subhajit Sengupta, Juhee Lee, Yuan Ji

BayClone-C is an ultra-fast method to estimate the cancer cell fraction (CCF) for each single nucleotide variant (SNV) and group SNVs into clonal or subclonal clusters with different CCF values.

**Preprocessing**

Using the consensus tumor purity ($\rho$), the consensus copy number calls and total and variant read counts for each single nucleotide variant (SNV), the algorithm estimates multiplicity and cancer cell fraction (CCF) for each SNV. Variant allele frequency (VAF) of SNV $s$ is given by the ratio $n_s/N_s$ where $N_s$ is the total number of reads and $n_s$ is the number of reads with variant sequence. We assume

$$\text{VAF}_s = \frac{CN_s^M * \rho}{\rho * CN_s^T + (1-\rho) * CN_s^N}$$

where $CN_s^T$ and $CN_s^N$ are the total copy number at the SNV $s$ locus in tumor cells and normal cells, respectively. $CN_s^M$ is the mutation copy number of SNV $s$, i.e. the average number of copies of the mutant allele across subclones. We consider the case where SNV $s$ lies inside a clonal copy number segment. Denote the allele-specific copy numbers at the SNV $s$ locus by $q_{1s}$ and $q_{2s}$ (with $q_{2s} \geq q_{1s}$). Therefore, the possible integer multiplicity values are $\{1, \cdots, q_{2s}\}$. Multiplicity for an SNV is estimated as the minimizer $m_s = argmin_{t\in\{1,\cdots,q_{2s}\}} \left|1 - \frac{CN_s^M}{t}\right|$. We then estimate CCF for SNV $s$ as $\widehat{CCF} = \frac{CN_s^M}{m_s}$.

**Algorithm**

We fit a finite mixture of Gaussian distributions to the estimated CCFs. We use the expectation-maximization (EM) algorithm (Dempster et al., 1977) to obtain the maximum likelihood estimate (MLE) of the mixture distribution. We run EM with different numbers of mixture components starting from one to seven and select the optimal number of components based on the Bayesian information criterion (BIC)(Schwarz, 1978). The R package *mclust* (Fraley, 2016; Scrucca et al., 2016) is used for implementation.

**Postprocessing**

A postprocessing step of merging the initial CCF clusters is conducted where each CCF cluster corresponds to a Gaussian distribution in the previous step. The merger is needed because it is possible that Gaussian mixture components are not sufficiently separated from each other to be interpreted as different CCF values (Hennig, 2010), especially when a set of CCF values can be described by unimodal distributions that are not Gaussian. We use one of the modality-based merging methods known as the ridgeline unimodal method (Ray and Lindsay, 2005). This method aims to find a partition of mixture components such that it would create unimodal clusters but further merging would generate a non-unimodal cluster. See Fig. 12 for an example. After merging all the Gaussian mixture components, we obtain a set of CCF clusters. Then as a final step, we collapse all the CCF clusters with cluster means greater than 0.9 into a single clonal cluster with the new cluster mean equal to 1.0. SNVs in the final CCF cluster with mean equal to 1 or less than 1 are considered clonal or subclonal, respectively. Clustering membership gives the assignment of SNVs to each cluster. We merge the Gaussian mixture components using the *mergenormals* function in the R package *fpc* (Hennig, 2010).

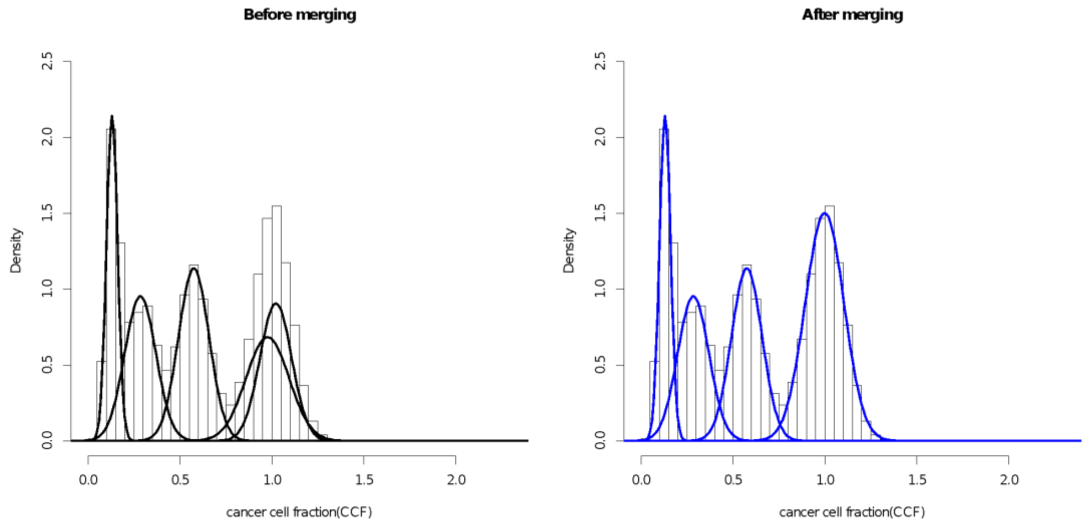


*Fig. 12 An example of merging using a PCAWG sample SA6164. Left and right panels are CCF clusters before and after merging of Gaussian components.*

#### 2.1.2 Ccube

Ke Yuan, Geoff Macintyre, Florian Markowetz

**Preprocessing**

The Ccube pipeline uses the consensus copy number, consensus purity and variant and reference allele read counts from consensus somatic point mutation calls.

**Purity**

The Ccube pipeline also produces an independent purity estimate using mutations from balanced copy number regions. For samples without whole-genome duplication, only mutations in normal copy number regions are included. For samples with whole-genome duplication, all mutations in balanced copy number regions are included. We then convert the allele frequencies to cellular prevalence estimates using the following equation:

$$f_i = \frac{\eta_i n_{i,tot_t}}{2((1-\eta_i)n_{i,tot_n} + \eta_i n_{i,tot_t}} \quad (1)$$

Where $f_i$ is the VAF of the $i$th mutation obtained as the ratio between variant and wild-type allele counts, $\eta_i$ is the cellular prevalence of the $i$th mutation, $n_{i,tot_n}, n_{i,tot_t}$ are the total copy number of normal and tumor populations respectively. We then cluster the $\eta_i$ using student's-t mixture model. The model is fitted with the variational Bayes approach described in (Archambeau and Verleysen, 2007). The purity corresponds to the component with the largest mean. In addition, the eligible component must have at least more than 1.5% of mutation assigned to it.

**Algorithm**

Ccube is a Bayesian mixture model for clustering cancer cell fractions. Each mutation is assumed to be a sample from a mixture of Binomial distributions, each of which is parameterized by the total number of reads covering the position and expected VAF. The expected VAF, denote as $\mathbb{E}[f_i]$, is parameterized as the following:

$$\mathbb{E}[f_i] = \frac{\rho(m_i(1-\epsilon) - n_{i,tot_t}\epsilon)}{(1-\rho)n_{i,tot_n} + \rho n_{i,tot_t}} CCF_k + \epsilon$$

Where $CCF_k$ is the CCF of the $k$th cluster, $\rho$ is the purity of the sample, $\epsilon$ is the sequencing error. The multiplicity is determined during clustering by the following:

$$m_i = \underset{m_i \in 1,\dots,n_{i,maj}}{\operatorname{argmax}} \sum_k \mathbb{E}_{q(z_{i,k},CCF_k)}[\log p(b_i | d_i, \mathbb{E}[f_i], z_{i,k} = 1)]$$

Where $n_{i,maj}$ is the copy number of the major allele at the $i$th mutation, $z_{i,k}$ is 1 if the $i$th mutation is assigned to the $k$th cluster and 0 otherwise. $b_i$ and $d_i$ denote the variant and total read counts, respectively. $p(.|.)$ is a conditional distribution, $\mathbb{E}_{q(.)}[.]$ is an expectation function with respect to $q(.)$, and $q(.)$ is an approximation to the posterior distribution of the arguments. This approximation is obtained by the variational Bayes approach. The number of cluster is handled by a truncated Dirichlet Process prior for cluster-wise parameter $CCF_k$. Full details on the Ccube model will be presented elsewhere.

**Postprocessing**

The solution with the best lower bound to the marginal likelihood of the model is selected. The solution consists of 1) posterior distributions of $CCF_{1,\dots,K}$ in terms of means and variances; 2) posterior probabilities of each mutation to be assigned to all clusters and final assignments; 3) multiplicities and observed CCFs based multiplicities. Clusters with less than 1% of mutations assigned are removed. The mutations are re-assigned in a variational expectation step. Clusters with mean CCF less than 10% apart are merged by re-running the fitting algorithm with the merged cluster configuration. A typical graphical summary can be found in Fig. 13.

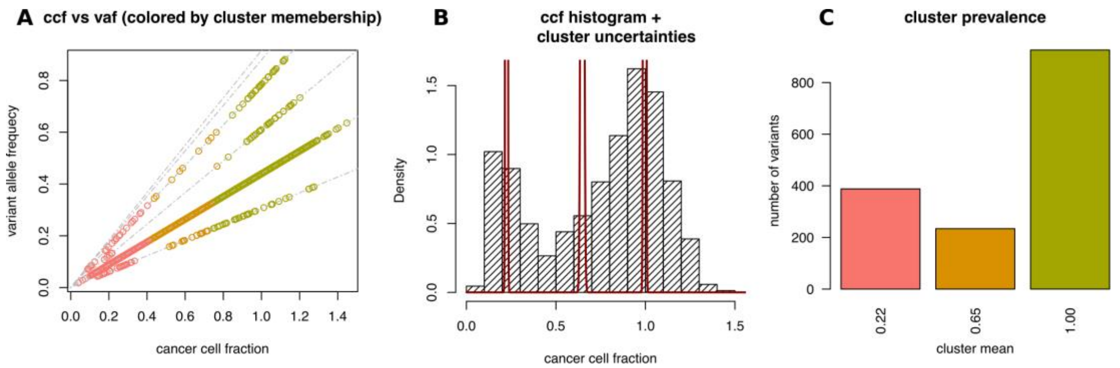


*Fig. 13 Ccube results summary for sample SA518422.* ***A)*** *Scatterplot of VAF and CCF. Each point in the figure is a mutation color coded by its cluster membership. The grey dashed lines are all possible linear mappings (eq. 1) determined by copy number and multiplicity configurations in the sample.* ***B)*** *Histogram of observed CCFs. The red solid line shows the approximated posterior distribution of CCF cluster centers. The peak at CCF=1 corresponds to the clonal cluster.* ***C)*** *The number of variants assigned to each CCF cluster. Each CCF cluster is labelled by its cluster center.*

#### 2.1.3 CliP (Clonal structure identification through Penalizing on pairwise difference)

Kaixian Yu, Hongtu Zhu, Wenyi Wang

**Preprocessing**

CliP is flexible on inputs: it requires either total copy number or allele-specific copy number, a rough estimation of purity, and single nucleotide variant read counts. However, the ideal set of inputs are allele-specific copy number profile, purity estimates, and single nucleotide variant read counts (counts supporting the variant and reference alleles), as are available in this study. The multiplicity of each SNV $i$ is estimated by,

$$m_i = [\frac{r_i}{n_i \rho}(\rho C_i^V + 2(1-\rho))]_0,$$

where $r_i$ is the variant allele read count, $n_i$ is total read count, $\rho$ is purity, and $C_i^V$ is the clonal total copy number. The operator $[x]_0$ is defined as: $[x]_0 = max\{1, [x]\}$, where $[x]$ represents the closest integer from $x$.

**Algorithm**

CliP assumes each variant read $r_i$ comes from a binomial distribution $Binom(n_i, \theta_i)$, where $\theta_i$ can be expressed as,

$$\theta_i = \frac{\phi_i m_i}{2(1-\rho) + \rho C_i^V}$$

where $\phi_i$ is the cellular prevalence (CP), $C_i^V$ the clonal total copy number, and $m_i$ the multiplicity of the SNV. Here, $\phi_i$ is our main interest. To ensure $\phi_i's$ fall in [0,1], a logit transformation is necessary; therefore,

$$\phi_i = \frac{e^{\omega_i}}{1 + e^{\omega_i}}$$

and,

$$\theta_i = \frac{e^{\omega_i} m_i}{(2(1-\rho) \ + \ \rho C_i^V)(1+e^{\omega_i})}$$

Then we formulate the problem as minimizing a penalized profile likelihood:

$$argmin_{\omega_i}\{ \frac{1}{2}\sum_{i=1}^{N} \frac{n_i(\theta_i - \hat{\theta}_i)^2}{\theta_i(1-\theta_i)} + \sum_{1\le i\le j\le N} p_\lambda(|\omega_i - \omega_j|) \},$$

Where $p_\lambda(x)$ is a penalizing function, one may use a LASSO (Tibshirani, 1996) type penalty (L1), MCP (Zhang, 2010), SCAD (Fan and Li, 2001), etc.

The minimization problem is solved by Alternating Direction Method of Multipliers (ADMM). Distinct $\phi_i's$ $(\omega_i's)$ represent different clusters.

**Postprocessing**

A series of $\lambda's$ are used to generate solutions, selecting an appropriate $\lambda$ can be achieved by information criterions (such as BIC, AIC). We also propose a bootstrap log-likelihood ratio test to select the penalizing parameter $\lambda$, which is much more computationally intensive. When an appropriate $\lambda$ is selected, we further conduct a filtering step in which a cluster with fewer than 1% SNVs is merged with the closest larger cluster.

The purity can be estimated if we further make the assumption that the clonal cancer cell frequency (CCF) is 1. Under such an assumption, the purity is set to be the largest $\phi_i$. If the estimated purity is different from the initial input purity, i.e. larger than 0.01 in actual implementation, a rerun of the whole pipeline is performed with this newly estimated purity until the difference between two consecutive estimates is less than 0.01.

##### 2.1.4 cloneHD

Ignacio Vázquez García, Ville Mustonen

Given a mixed sample of the genomes of clones that have accrued single-nucleotide variants (SNVs), cloneHD reconstructs their evolutionary history from bulk DNA sequencing. This problem consists of three parts: (i) identification of subclones, (ii) reconstruction of subclone-specific profiles, and (iii) inference of evolutionary relationships between subclones.

**Preprocessing**

We used consensus copy number and consensus purity as priors for cloneHD SNV clustering. The priors that cloneHD can use are the mean total copy number and the available copy number per locus. The mean total copy number is the mean copy number for each SNV locus of the entire sample, including both tumor and normal parts. The available copy number prior depends on the major allele state of the locus and allows mutations to be in states from zero up to the major allele state. For example, if a mutation was present before a chromosome underwent copy number gains, it would be present in all gained copies, whereas if it happened after the gain it would be present in fewer copies.

**Algorithm**

The cloneHD algorithm described in (Fischer et al., 2014)) identifies subclones and infers a subclone-specific posterior probability of SNV genotypes using a discrete state space HMM. The algorithm uses a genotype-based description, where the SNV genotypes of each subclone are encoded as hidden variables. Each subclone has an identity across the whole genome without a tree constraint on the relationship of the subclones. Mapping between subclones and clusters implies that when the model dimensions are increased by one subclone, two possible subclonal clusters are added at once (Fig. 14). As a result, alternative subclonal architectures can be problematic to resolve based on single samples from a tumor. To alleviate this problem, we extended cloneHD to allow for an SNV prior (‘tree prior’) to be learned from the data. This optionally enforces a tree structure on the SNVs and addresses the relationships between subclones. The other priors used in SNV mode reflect copy number and are described in the Supplementary Information of the cloneHD paper, equations 34-35 (Fischer et al., 2014). They are computed from copy number and B allele frequency data, as described above.

To build an informative SNV prior, we take into account the lineage relations between subclones. Therefore, we treat the relation between subclones as a rooted tree, where the normal tissue is taken as the root, from which the clonal mutations of the sample are derived. Each node in the tree represents a cluster of mutations. On the way from this root to the terminal nodes, we assume that a cell may acquire a new mutation exactly once. Each of the connecting edges then represents mutations that distinguish the child from the parent node. In the subtree below this state switch, the genotype always remains mutated. This is commonly referred to as the 'infinite sites' assumption, because it is unlikely for the same mutation to occur twice.

We can describe the tree by the SNV genotype prior. The constraints on the tree phylogeny restrict the SNV genotype prior and thus the family of lineage trees that are possible. Here we implemented a set of tree priors shown in Fig. 14. Resolving the complete space of tree topologies is currently limited by the resolution of the data, but it may be possible to implement a generative model of tree structures, instead of constraints on the prior, which would require an integration over all possible trees.

**Postprocessing**

Finally, we carried out model selection to determine the number of subclones that best explain the data. We required an improvement of 50 units of log-likelihood (for details of the scoring see the Supplementary Information in (Fischer et al., 2014). We then converted from a representation of subclone genotypes to mutation clusters. To remove clusters that are not supported by sufficient high-confidence mutations, we required that the total sum of posterior probability of mutations to belong to a given cluster at 0.9 or higher probability was required to be >10. This corresponds to a minimum of ~10 high confidence mutations assigned to each cluster.

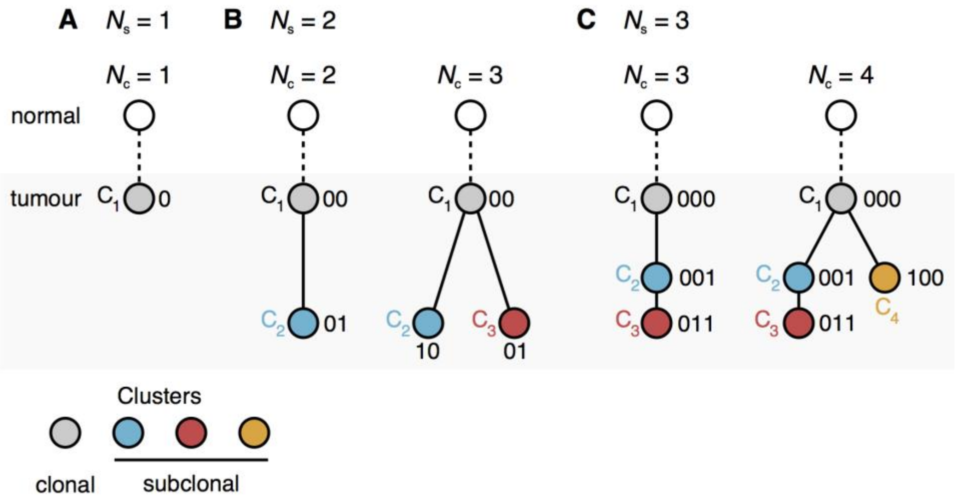


*Fig. 14 Mapping between subclone genotypes and clusters. There is a correspondence between subclonal genotypes and mutation clusters. Each node in a tree is a cluster, and mutations are acquired along the edges of the tree, from the root (top) to the leaves (bottom). Each node is labelled by the cluster genotype. The ancestral node linking to the clonal cluster is also shown. SNV genotype priors that are compliant with the infinite sites assumption constrain the set of tree topologies. The examples show families of trees compliant with this assumption relating* ***(A)*** $N_s = 1$, ***(B)*** $N_s = 2$ *and,* ***(C)*** $N_s = 3$ *genotypes, each of which contains* $N_c$ *mutation clusters.* $N_s = 1$ *corresponds to a scenario where there are no subclones in the sample, so all mutations belong to the clonal cluster denoted as a grey node. The* $N_s = 2$ *scenario has a single subclone corresponding to one clonal and one or several subclonal clusters and* $N_s = 3$ *more than one subclone describing linear or branched evolution. For example, with* $N_s = 2$ *subclones, genotype* $g = 00$ *denotes all clonal mutations,* $g = 10$ *mutations that are private to the blue cluster and* $g = 01$ *mutations that are private to the red cluster. The tree prior extends cloneHD by inferring a prior for a mutation to belong to one of the clusters shown. Assuming a uniform mutation rate the prior weights can be interpreted to relate to the branch lengths of the tree leading to each node.*

##### 2.1.5 CTPsingle

Salem Malikic, Nilgun Donmez, Cenk Sahinalp

**Preprocessing**

As input, CTPsingle (Donmez et al., 2017) takes reference and variant read counts for single nucleotide variant (SNV) calls. CTPsingle expects only somatic SNVs in its input; all germline SNPs should be discarded in advance. It also expects the mutations to reside in copy number neutral regions of the genome. In other words, CTPsingle assumes that all mutations in the input are heterozygous SNVs belonging to the regions that have not been affected by copy number aberrations. Mutations on non-autosomal (i.e. X and Y) chromosomes can be included if the gender of the patient is given. Tri-allelic variants are also discarded from the input. For each SNV, the numbers of reads supporting variant and reference allele are used to obtain a clustering of mutations as described in the next section.

**Algorithm**

For the given set of SNVs, CTPsingle uses a beta-binomial model to perform clustering of mutations in read count space. Denoting the number of variant reads and the total number of reads covering an arbitrary mutation $i$ as $y_i$ and $n_i$, respectively, we assume that $y_i$ is binomially distributed with unknown probability of success $p_i$:

$$y_i \mid (n_i, p_i) \sim Binom(n_i, p_i).$$

The probability parameter $p_i$ is assumed to be generated from a Dirichlet process as given below:

$$p_i \mid G \sim G$$
$$G \mid (\alpha, G_0) \sim Dir(\alpha, G_0)$$

The baseline distribution $G_0$ is taken to be a Beta-binomial distribution with parameters $a_1$ and $b_1$. Since the Beta prior is conjugate to the Binomial distribution, resulting in a Beta-binomial posterior, inference can be performed using a standard Markov Chain Monte Carlo (MCMC) method as described in (MacEachern, 1998). Based on our earlier observations on simulated and real data, parameters $\alpha$, $a_1$ and $b_1$ were set to 0.001, 5.0, 5.0, respectively. In order to mitigate overclustering due to a fraction of noisy mutation calls, we weakly constrained the minimum cluster size to at least 2% of the total number of mutations.

**Postprocessing**

From the clustering step, we obtain the number of subclones (i.e. clusters) $k$ and the assignment of mutations to subclones. After the clustering step, the mean allelic frequency $s_j$ for each subclone $j = 1, 2, \dots, k$ is calculated as the average of allelic frequencies $y_v/n_v$ of all mutations $v$ belonging to the cluster $j$. As the clustering is performed using variant read counts of heterozygous SNVs, cellular prevalence $s_j^*$ of subclone $j$ is obtained as $s_j^* = 2s_j$. Given the cellular prevalence of each subclone, the purity of the sample is inferred as $p = \max(s_j^*)$ where the maximum is taken over all indices $j \in \{1, 2, \dots, k\}$. The output of the clustering step is illustrated in Fig. 15 where mutations are merged together and their counts are shown based on the values of $2y_v/n_v$ that reflect their cellular prevalence.

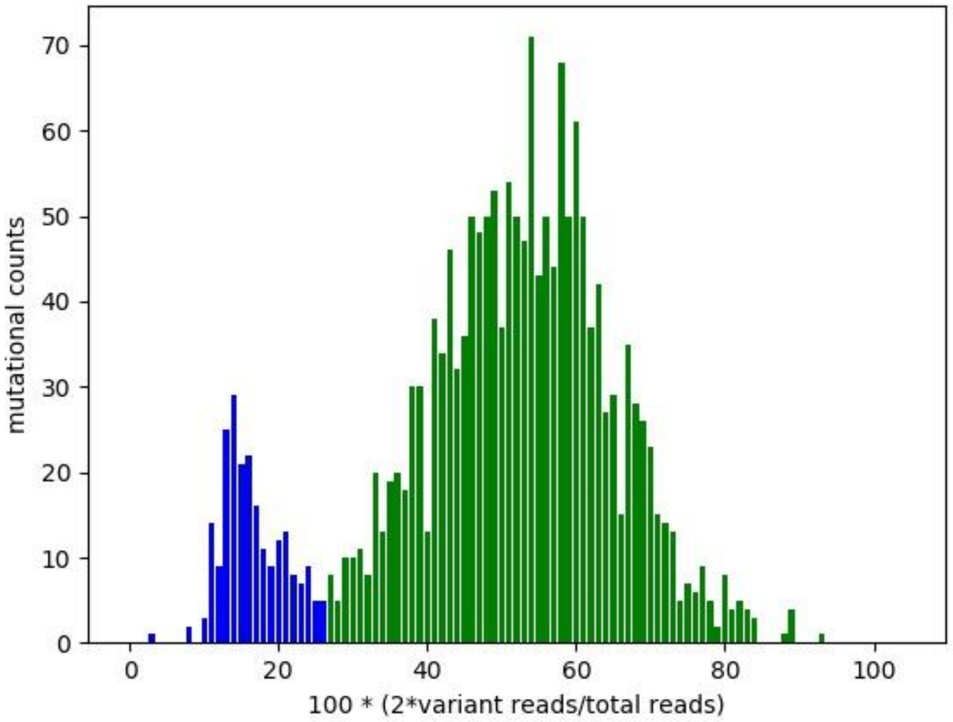


*Fig. 15 Barplot of CTPsingle clustering result for sample SA530652. Different colors represent different subclones. Although clustering is performed in read counts space, for the sake of visualization, mutations are merged based on the ratios of variant and total reads shown on the x-axis and scaled to the interval [0,100]. For each integral value on the x-axis the number of mutations having the considered ratio is shown on the y-axis.*

#### 2.1.6 DPClust

Stefan Dentro, David Wedge, Kevin Dawson, Peter Campbell, Peter Van Loo

**Preprocessing**

The DPClust pipeline takes as input an allele-specific (subclonal) copy number profile, a purity estimate and single nucleotide variants (SNVs) reported as chromosome, position and allele counts. The variant allele frequency (VAF) of each SNV is calculated through the reported allele counts using only the reads reporting the variant and wild-type alleles. With the VAF established, we obtain an estimate of the cancer cell fraction (CCF) of each SNV in three steps: First we approximate the copy number state of the SNV, its multiplicity, through:

$$u_i = \frac{f_i}{\rho}\left[\rho n_{tot_t} + (1-\rho) n_{tot_n}\right]$$

Where $f_i$ is the VAF of SNV $i$, $\rho$ represents the tumor purity and $n_{tot\_t}$ and $n_{tot\_n}$ is the total copy number at the locus of SNV $i$ in tumor and normal cells respectively. The multiplicity is established by rounding the approximate mutation copy number state $u_i$:

$$m_i = \begin{cases} 1, & u_i < 1 \\ |u_i|, & u_i \geq 1 \end{cases}$$

Finally, the CCF estimate of each SNV is obtained through

$$CCF_i = \frac{u_i}{m_i}$$

**Algorithm**

DPClust is based on a Dirichlet Process (DP) that models the SNVs as drawn from a statistical distribution that consists of an unknown number of distributions. A posterior estimate of the number of internal distributions is obtained through Markov Chain Monte Carlo (MCMC) using stick-breaking. To account for read sampling variation on the number of observed reads, each variant is modelled as a draw from a binomial distribution. A description of the algorithm's characteristics is provided in (Bolli et al., 2014).

**Postprocessing**

After completing the MCMC iterations, we aim to obtain three estimates: (1) An estimate of the finite number of distributions (cell populations), $K$, that are present in the input data, (2) the proportion of tumor cells that each population consists of ($CCF_k$) and (3) likelihoods of each SNV belonging to each population. The number of cell populations $K$ is determined by finding peaks in the posterior weight density (Fig. 16). In each iteration $j$, the stick-breaking procedure assigns a weight $\omega_{k\_j}$ to each cluster that represents its size and the cluster has a $CCF_{k\_j}$. Over many iterations, weight accumulates in the CCF space, where a large amount of weight corresponds to a high likelihood of the existence of a mutation cluster. We then obtain an estimate of $K$ by obtaining all local maxima in the weight density.
With the $K$ clusters and their locations ($CCF_k$) established, SNVs can be assigned to clusters. We first establish the CCF area covered by each $k \in K$ by finding the CCF location between each pair of neighbouring clusters that corresponds to the minimum density. The minimum density on either side of a cluster represents its upper and lower CCF bound. Probabilities of a mutation belonging to a cluster are then established by accounting how often a SNV would have been assigned to each $k$ throughout the MCMC iterations. Finally, small clusters are removed: Clusters smaller than 1% of total mutations in tumors with fewer than 150 SNVs and clusters with fewer than 30 SNVs otherwise.

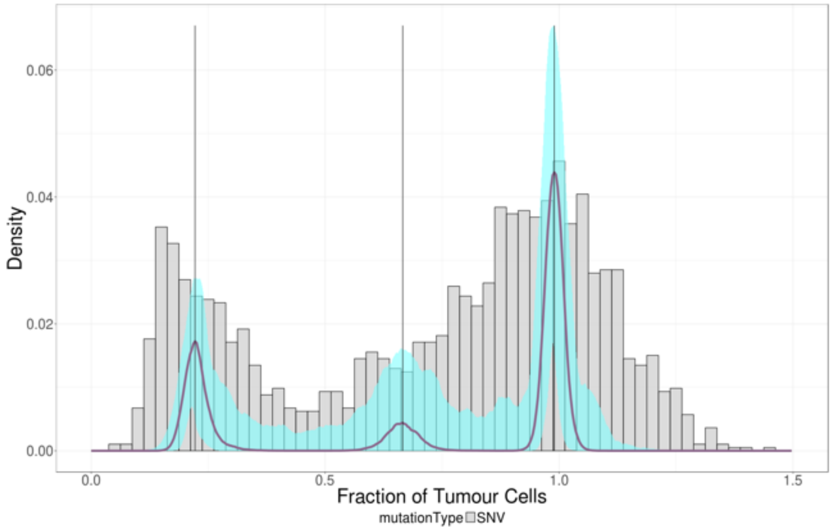


*Fig. 16 Posterior density (purple) and confidence interval (cyan) of cluster locations in cancer cell fraction (CCF) space for sample SA518422. The number of mutation clusters is obtained by finding peaks in the density. The peak at CCF=1 represents the clonal SNVs carried by all tumor cells, the other two peaks represent two subclones.*

#### 2.1.7 PhylogicNDT

Ignaty Leshchiner, Dimitri Livitz, Daniel Rosebrock, Gad Getz

ABSOLUTE computes a probability distribution for the cancer cell fraction (CCF) of every mutation (independently). These distributions factor in sample purity and local absolute copy number.

PhylogicNDT utilizes called somatic variants and absolute copy number to perform Dirichlet Process clustering on the calculated CCF of the somatic variants in order to learn the underlying number of clusters from the data (Brastianos et al., 2017; Landau et al., 2013; Shaw et al., 2016).

In order to learn the underlying subclonal structure, we applied the Dirichlet Process as described by (Escobar and West, 1995)) to the discretized CCF distributions of the variants. The mixing parameter was resampled iteratively from a prior gamma distribution as follows:

1) Sample eta: $\eta = B(\alpha + 1, n)$ where $n$ is the number of variants and $B$ is the beta function.
2) Sample alpha:
$$\alpha = \pi_\eta G(a + k, b - log(\eta)) + (1 - \pi_\eta)\ G(a + k - 1, b - log(\eta))$$
[equation 13 (Escobar and West, 1995)] where $\eta$ is given above, $k$ is the number of clusters, $a$ and $b$ are given by the prior, G(a,b) is a sample from the Gamma Distribution with shape and scale a and 1/b respectively, and $\pi_\eta$ is given by:
$$\frac{\pi_\eta}{1 - \pi_\eta} = \frac{a + k - 1}{n(b - log(\eta))}$$

In each iteration, the probability of opening a new cluster is proportional to:

$$\frac{\alpha}{n - 1 + \alpha}$$

Within each iteration a Gibbs sampler over the mutations is run, where the probability of a mutation joining a particular cluster is proportional to:

$$\frac{m}{n - 1 + \alpha}$$

where $m$ is the size of each cluster and is dictated by the likelihood of a particular mutation CCF distribution to be joined with other mutations present in the cluster at a given iteration.

Once the Dirichlet Process was completed for 10,000 iterations, we integrated the mutation probability distributions across all iterations and variants into a posterior density.
The prior mutational CCF distributions and the posterior cluster densities can be visualized with the distribution of clustered mutations across chromosomes (expected to be uniform) for quality control.

See the following Fig. 17 for an illustration of PhylogicNDT clustering.

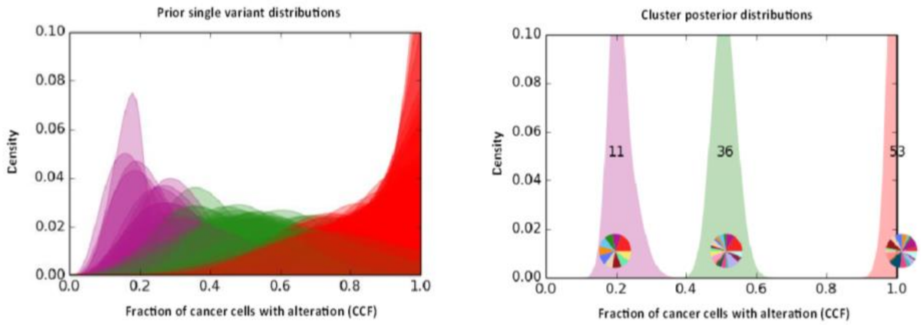


*Fig. 17* ***PhylogicNDT clustering.*** *The prior mutational CCF distributions (left) and posterior cluster densities (right). Colors represent the final subclonal assignment of mutations. Pie charts show distribution of clustered mutations across chromosomes.*

The number of clusters, $k$, was then computed from this distribution by identifying all local maxima corresponding to peaks with sufficient density (1%). Mutations were then assigned to the best fitting detected cluster based on the similarity of their MCMC trace to the peak.

#### 2.1.8 PhyloWGS

Jeff Wintersinger, Amit Deshwar, Shankar Vembu, Quaid D. Morris

**Preprocessing**
We subsampled simple somatic mutations (SSMs, also referred to as SNVs) to a maximum of 5,000 per sample, selecting SSMs according to the provided priority list used across methods. We selected only SSMs located in genomic regions where the consensus copy-number state was known, allowing the method to correct for copy-number changes' effects on observed variant allele frequencies.

PhyloWGS places copy-number aberration (CNA) events as mutations in its inferred phylogenetic tree based on their reported cellular prevalence. Although PhyloWGS can utilize subclonal CNAs, only clonal consensus CNAs were available for these analyzes, and so only those were provided to PhyloWGS. We modified PhyloWGS so that all CNAs were assigned to the same cluster, but did not require them to be assigned to the clonal cluster, thus permitting clonal clusters defined by only SSMs.

To input a CNA into PhyloWGS, we must know not only the cellular prevalence $p$ of the CNA, set here to the consensus purity, but also the uncertainty in the cellular prevalence estimate (i.e., the standard error $e$). In the original PhyloWGS, we communicated these values to our Bayesian model by generating a "pseudo-SSM" for each CNA, and setting its variant reads $v$ and total reads $d$ to the values PhyloWGS would expect for an SSM with the same cellular prevalence and standard error.

Given the read depth $d$ for the pseudo-SSM, we set $v$ to represent the reported cellular prevalence by setting $v = 0.5*pd$, meaning that $v$ represents the expected number of variant reads for an SSM in a diploid region with a cellular prevalence of $p$. In general, we recommend setting the read depth $d$ so that the standard error of the pseudo-SSM's implied cellular prevalence would be equal to the standard error in the estimate of the CNA's cellular prevalence. As PhyloWGS uses a binomial likelihood, we can approximate the standard error of the pseudo-SSM's implied cellular prevalence as

$$e = \sqrt{\frac{p(1-p)}{(d+1)}}$$

and then solve for $d$. Note that $e$ scales as the inverse of $d$, meaning that higher read depths produce smaller errors.

As standard errors for the CNAs were not available, we estimated them using a rule of thumb that let us compute $d$ directly. Because most CNA-calling methods use heterozygous SNPs to determine cellular prevalence, and there are on average seven heterozygous SNPs per 10 kb of genomic sequence, we assumed that the standard error in the estimate of the cellular prevalence for a 10 kb CNA would be approximately equal to the cellular prevalence estimate error for an SSM cluster containing seven heterozygous SNPs. As such a cluster's $d$ value would be the sum of read depths for all seven SNPs, we set the $d$ value for the pseudo-SSM representing a CNA $i$ to be $d_i = L_i * R * D$, such that $L_i$ indicates the length (in kb) of the genomic region affected by the CNA, $R$ indicates the heterozygous SNP rate used to infer the CNA (i.e., 7 heterozygous SNPs per 10 kb), and $D$ indicates the average read depth, taken as the median number of total reads across all SSMs. Next, we merged clonal CNAs into single large CNA events by summing their $d_i$ values, preventing them from being assigned independently of one another to different (subclonal) populations in the inferred tree, producing $d_S = \Sigma d_i$. Finally, we determined the weight $d_C$ for the single clonal CNA event $C$ as $d_C = \min([d_S], A * D)$, such that $A$ indicates the average number of SSMs per tumor (set to $A = 3000$, as this was approximately the average number of clonal SSMs in the PCAWG dataset). $A$ represents the maximum confidence level we would have in the cellular prevalence estimate of a CNA in terms of equivalent number of SSMs. Thus, we inform the model that the standard error in the cellular prevalence estimate for a CNA must be no smaller than the standard error associated with a cluster of 3000 SSMs all possessing the sample's average read depth. That is, we constrained the confidence of cellular prevalence estimates from SSMs and from clonal CNAs to be approximately the same. Consequently, for an average read depth of 50 and a purity of 80%, which are typical values seen in PCAWG datasets, this minimum standard error is:

$$e_{min} = \sqrt{\frac{p(1-p)}{(d+1)}} = \sqrt{\frac{0.8(1-0.8)}{(3000 * 50 + 1)}} \approx 0.001$$

**Algorithm**
The PhyloWGS algorithm is fully described in (Deshwar et al., 2015). In brief, we build upon the tree-structured stick-breaking process (TSSB) prior, using Gibbs sampling to sample trees, population frequencies, and SSM assignments via MCMC. After discarding the initial 1000 burn-in trees, we recorded 2500 trees for each dataset. The population frequencies were determined via 500 iterations of Metropolis-Hastings for each Gibbs sampling iteration.

**Postprocessing**
PhyloWGS generates clone trees, while the PCAWG consensus efforts required only mutation clusters. As such, we discarded all tree-structure information, retaining only mutation clusters and their assigned frequencies from each MCMC sample. Clusters bearing less than 1% of total mutations were removed, with their SSMs reassigned on a per-mutation basis to whichever cluster offered the highest binomial likelihood assignment. We discarded polyclonal trees (i.e., trees bearing multiple independent clonal cancerous populations), rerunning PhyloWGS with a different random seed if more than 80% of sampled trees for a given dataset were polyclonal. On the remaining monoclonal trees, we removed "superclonal" populations. Superclonal populations were deemed to occur when the clonal population had a single child population bearing at least three times as many SSMs as the purported clonal population, with the child possessing no more than one child population itself (but any number of descendant populations). After removing superclonal populations, we assigned their SSMs to their single descendant, then set the descendant's cellular prevalence to the mean of its own cellular prevalence and that of the former clonal population, weighted by the number of SSMs originally in each.

To generate a single solution for each dataset, we first determined the mode number of cancerous populations $K$ across all MCMC samples, restricting further steps to sampled trees that also had $K$ populations. To match populations between trees, we sorted the $K$ populations from each sampled tree in order of descending cellular prevalence. Next, to establish SSM assignments to populations, we counted the number of times a given

SSM was assigned to each of the $K$ possible populations, reporting the mode assignment as our answer. To determine the per-population number of mutations, rather than simply report the count of summarized SSM assignments for each population, we better summarized the posterior consensus concerning mutation count by calculating the mean number of SSMs per population across selected trees. As the per-population values could then take non-integer values, we rounded these values to the nearest integers, distributing rounding errors by adjusting mutation counts upward or downward by one in the population with the most mutations, and continuing to modify mutation counts in populations ordered by descending number of mutations until the rounding error was fully distributed. Thus, though the per-population mutation counts may not match the reported SSM assignments, the sum of counts will match the total number of mutations assigned. Finally, we determined summary cellular prevalences for each population simply by calculating the mean cellular prevalences of each population across selected trees.

##### 2.1.9 PyClone

Ke Yuan, Geoff Macintyre, Florian Markowetz

**Preprocessing**

The PyClone pipeline uses consensus copy number, consensus purity and variant and reference allele read counts from consensus somatic point mutation calls (SNV).

**Algorithm**

The description of PyClone is in (Roth et al., 2014). The model assumes that each mutation is sampled from a mixture of Beta-Binomial model. The cluster-wise parameter is a draw from a Dirichlet Process prior. For copy number, we use the parental copy number prior. The prior assumes the composition cells in terms of normal, reference and variant populations. The reference and variant populations are cancer cells. We set the two to have the same copy number profiles. The multiplicities are chosen among $[1, n_{i,min}, n_{i,maj}]$. The multiplicities are marginalized out with a prior, enforcing equal probabilities. The model is fitted to data with MCMC.

**Postprocessing**

After discarding burn-in samples, we use MPEAR implemented in the R package mcclust to compute consensus assignments from the assignment traces. CCF cluster centers are obtained as median of CCF traces from the mutations assigned to the same cluster. The results consisted of 1) CCF centers; 2) final assignments; 3) multiplicities and observed CCFs. Clusters with less than 1% of mutations assigned are removed. Clusters with their CCF centers less than 10% apart are merged.

#### 2.1.10 Sclust

Martin Peifer, Yupeng Cun, Tsun-Po Yang

From copy number states, estimated purity, and variant allele frequency we derive the cancer cell fraction of each mutation similar to DPClust. This serves as input for the mutational clustering module of Sclust. The general idea of our mutational clustering approach is that the histogram of SNV cancer cell fractions can be decomposed into a distribution of clonal populations and a distribution of sampling error due to the finite coverage. The distribution of the sampling error is approximated by a Gaussian with zero mean and average standard distribution derived from the cancer cell fractions. Next, the distribution of the clonal populations is derived by a deconvolution of the histogram of cancer cell fractions with the distribution of sampling error. To solve this ill-posed inverse problem, we use smoothing splines with a fixed smoothing parameter. Peaks in the estimated distribution of clonal populations constitute the cluster location of subclonal populations. Finally, mutations are assigned to the most probable peak using a binomial model.

#### 2.1.11 SVclone

Geoff Macintyre, Ke Yuan, Florian Markowetz

For a detailed explanation of the SVclone algorithm and pipeline please see (Cmero et al., 2020) Here we provide further details regarding the application of SVclone to PCAWG samples.

**Preprocessing**
Using the consensus SV vcf, consensus SNV vcf, final consensus copy number profile, consensus purity estimate, and indexed WGS miniBAM file for each sample, the following preprocessing steps were carried out using SVclone:

- annotate - each of the SV pairs was annotated as coming from a deletion, inversion, translocation, or duplication
- count - normal and variant read counts for each SV were extracted from the miniBAM file and adjusted based on SV event type
- filter - the following filters were applied:
  - any SV or SNV that was not matched to a valid copy number segment was removed
  - any SV that lacked split or spanning supporting reads was removed
  - any SV less than 1000bp was removed
  - samples with greater than 4000 SNVs had their SNVs randomly downsampled to 4000 SNVs for clustering

**Clustering**
The co-clustering mode of SVclone was used to simultaneously cluster and determine the CCF of SNVs and SVs. MCMC was carried out for 25,000 iterations (12,500 burn-in) to approximate the posterior distributions over model parameters. During clustering SVclone's cluster merging procedure was used to merge clusters with overlapping CCF 95% confidence intervals. This procedure was repeated 8 times for each sample and the optimal run was chosen based on SVclone's BIC-like measure, svc_IC. alpha and beta

values for the gamma distribution prior over the Dirichlet concentration were set to 0.1 and 0.5 respectively.

**Postprocessing**

Cluster membership for the variants used during clustering, and cluster mean CCFs, were determined as per SVclone's default approach. Variants not used in clustering were retroactively assigned to the most likely cluster using SVclone's post_assign procedure. Low-count clusters were filtered out (< 10 variants or cluster contains < 1% of variants) and variants from these clusters were re-assigned to the second most likely cluster.

#### 2.2 Subclonal architecture consensus approaches

Kaixian Yu, Maxime Tarabichi, Amit Deshwar, Stefan Dentro, Ignaty Leshchiner, David C. Wedge, Quaid D. Morris, Peter Van Loo, Wenyi Wang

The 11 subclonal reconstructions described above were applied to the 2,778 cancer genomes in the PCAWG data set and we aimed to combine the output into a robust consensus subclonal architecture.

We applied the 11 individual callers, described in the previous subsection, to a subset of ultra-confident copy number segments from the copy number consensus (see 1. Copy number consensus). Segments were ordered by their confidence level and selected up until at least 75% of the genome was covered. This ensured that the most confident copy number segments were used and that those with lower confidence levels could not introduce spurious mutation clusters.

To find mutation clusters, the methods were run on the provided consensus SNVs in the selected copy number regions only. This yielded 11 subclonal architectures that are described by four essential outputs:

- Number of mutation clusters identified,
- Number of mutations in each cluster,
- Proportion of tumor cells that each cluster represents,
- Mutation assignments to each cluster - either probabilistic or hard assignments of all SNVs to the clusters.

A fifth optional output is the mutation multiplicities, i.e. the number of physical DNA copies bearing the mutation, which is not provided by all individual methods but is used by CICC, one of the consensus approaches, to recalculate the proportion of tumor cells represented by the new consensus clusters.

The output of the 11 callers was used as input to the consensus procedure. We developed three orthogonal approaches (described below **2.2.1 Consensus approaches**) that utilize different representations of the 11 subclonal reconstructions: WeMe uses cluster sizes and locations only, CSR uses mutation co-assignment probabilities, while CICC uses mutation-to-cluster hard assignments.

Each of these three methods returns output for three of the key subclonal architecture descriptors: the number of mutation clusters, the number of mutations in each cluster, and their cellular prevalence. The results reported in this paper are from the WeMe consensus method, but as we show below, we could have chosen CSR or CICC, which lead to similar results.

We ran individual methods together with the consensus approaches on two simulated datasets and on the PCAWG data. We derived a metric system in which we could score the results and evaluate robustness. The metrics and simulations are described in the following sections.

Importantly, individual methods perform downsampling when there are more than a given number of SNVs $N$ to be clustered, each method $m$ using its own $N{:=}N_m$ to consider downsampling to $N_m$ SNVs. Because CICC and CSR rely on mutation assignment to produce a consensus, we asked individual methods to downsample from a pre-randomized list of SNVs, taking the $N_{i,downsampled}$ first SNVs from that list as input, where $N_{i,downsampled}$ is the $i_{th}$ method's threshold for downsampling. This ensured a minimal overlap of $min_i(N_i)$ across methods to produce the consensus from.

Finally, to assign all SNVs and also indels and SVs, we provided the consensus subclonal architecture description, the full consensus copy number profile and all consensus SNVs, indels and SVs for which allele frequencies were available to the MutationTimer algorithm, described in (Gerstung et al., 2020). MutationTimer assumes each mutation cluster can be modelled by a beta-binomial and calculates probabilities for each mutation belonging to each cluster whilst also taking into account the size of the mutation clusters. This step yielded the final consensus subclonal architecture with the aforementioned key features, while also performing timing of mutations relative to gains to classify mutations as clonal early, clonal not specified, clonal late and subclonal.

#### 2.2.1 Consensus approaches

Because there is no straight answer to which output should be used to construct the consensus subclonal architecture nor how a method can integrate all outputs, we designed three independent consensus approaches, taking different inputs in order to build a consensus:

1) **Sparse clustering for subclonal reconstruction (CSR or "Caesar")**, which takes the co-clustering probability matrix output of each method, i.e. the soft cluster assignment and extract common signals through non-negative matrix factorization (NMF);

2) **Cluster-ID Consensus Clustering (CICC or "Chic")**, which takes the hard cluster assignment of each SNV and performs a hierarchical clustering on the distance of the assignments to identify SNVs that most often cluster together across methods;

3) **Weighted Median (WM or "Weme")**, which takes a weighted median of the locations and the proportion of SNVs assigned to the location to construct a consensus location profile, ignoring the assignment of individual SNVs.

Not only are the consensus methods based on different principles (Fig. 18) but they also vary in their pre- and post-processing steps (see following subsections).

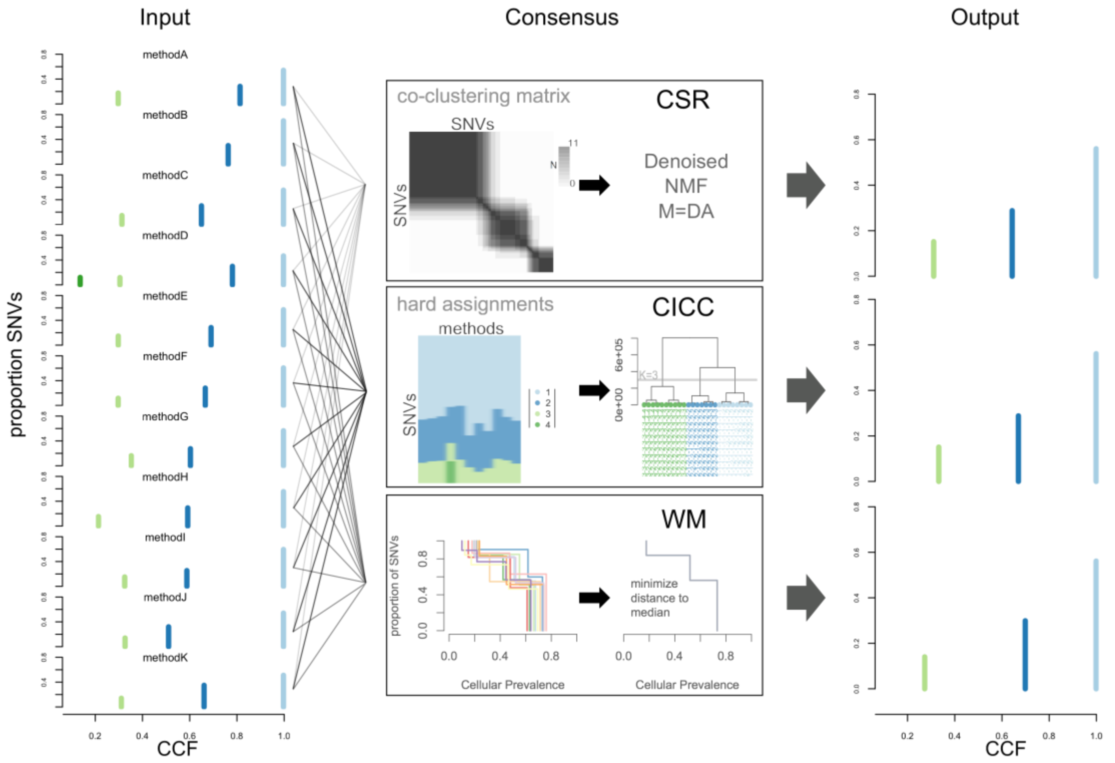


*Fig. 18 Workflow of the consensus approaches.* ***Input.*** *On the left, inputs are represented by their respective solutions in the space defined by CCF and proportion of SNVs of the clusters. Each cluster is represented by a vertical colored bar. Each of the 11 methods also provides mutation assignments corresponding to these solutions.* ***Consensus.*** *In the middle panels,*

*illustration of the three consensus approaches. They take different aspects of the individual methods as input and obtain the consensus from it in different ways (see main text).* ***Output.*** *Right panels, the output solutions of the three consensus methods in the same space as Input. Both CSR and CICC also provide mutation assignments to the clusters, not shown in that space and not used by MutationTimer.*

###### 2.2.1.1 Caesar (CSR – Sparse Clustering for Subclonal Reconstruction)

When there are several subclonal callers that make distinctive calls, we expect that the sum of co-clustering matrices (CCMs) contains a certain amount of incorrectly clustered pairs of SNVs, as well as differences in SNVs that define cluster boundaries. CSR aims to identify the valuable common information from all methods and discard the disagreeing information through decomposition, in order to construct a cluster topology that reflects the strong signals contained in the sum of CCMs across callers. Specifically, CSR decomposes the sum of co-clustering matrices (CCMs) across all individual methods into a sum of a low rank matrix (signal) and a sparse matrix (noise). The low rank matrix captures the clustering information, and the sparse matrix takes care of possible mis-clustering and method-specific errors.

Suppose $M_h$ is a CCM of size $N \times N$ for method $h$, where $N$ is the number mutations. The elements of $M_h$, $m_{ij}^{(h)}$ takes value 1 if the $i$- and $j$-th mutation was clustered together by method $h$, and 0 otherwise. The mean of CCMs from each method is defined as

$$M = \frac{1}{H}\sum_{h=1}^{H} M_h$$

where $H$ is the total number of methods to generate a consensus from.

To account for potential outliers, a thresholding was imposed on the mean CCM:

$$M = (m_{ij}) = \begin{cases} m_{ij}, m_{ij} \geq T \\ 0 \quad, m_{ij} < T \end{cases},$$

In the PCAWG project, there were 11 ($H = 11$) participating methods, and the threshold was set as $T = 0.2$.

Next, the mean CCM, $M$, is decomposed into a sum of two matrices:

$$M = DA + \Sigma$$

where $D \in R^{N\times K}$, $A \in R^{K\times N}$, and $\Sigma \in R^{N\times N}$. The product $DA$ is the low rank matrix (rank up to $K \ll N$), and $\Sigma$ the noise matrix. Such a decomposition is also called dictionary learning, where $D$ is the dictionary, and $A$ the code. Our main interest is matrix $A$, presenting a new set of features for mutations, which are less contaminated than the original features provided by $M$. We use the Python package SPAMS (version 2.1) to estimate $D$ and $A$. Once $A$ is obtained, a k-means algorithm is applied to $A = (a_1, a_2, \ldots a_N), a_i \in R^K$ to identify mutation clusters. CSR requires as input the number of clusters, $K$. In the PCAWG project, $K$ was chosen to be the median of all methods (rounded up if only even number of methods had results on the given sample).

With the clustering results, we further estimate the cellular prevalence (CP) for each cluster. Suppose the CP of SNV $i$ estimated by method $h$ is $\phi_i^{(h)}$, then the consensus

CP of SNV $i$ is defined as $\Phi_i = median\{\ \phi_i^{(h)}\ \}$. Subsequently, the cluster-specific CP is computed as

$$\Phi_C = \frac{1}{\sum_{i=1}^{N} I(i \in C)} \sum_{i=1}^{N} \Phi_i I(i \in C)$$

where $C$ denotes a mutation cluster, and $I(\cdot)$ is an indicator function, taking value of 1 if SNV $i$ belongs to cluster $C$, and 0 otherwise.

To make the obtained cluster architecture more biologically meaningful, several post processing steps are applied: 1) all the clusters with CP $>$ *the consensus purity* (i.e., super-clonal SNVs) are merged with the closest non-super-clonal cluster. 2) Clusters that are too close in CP are merged to form one new cluster. The minimum distance between two nearest clusters is calculated as the median of the shortest distances between two clusters called by individual methods. When most methods call only one cluster, the minimum distance is set as 0.05. 3) Clusters containing too few SNVs will be dropped. The thresholding of tiny clusters is also inferred from results of each method, by taking the median of the smallest cluster sizes. When most methods give only one cluster, the minimum size is set as 5% of total SNV or 50 when the total number of SNVs is fewer than 1,000.

In samples that are hyper-mutated, i.e., their number of SNVs is above 30,000, a subsampling strategy is conducted to address computer memory limitations when constructing the CCMs, to reduce to 25,000 SNVs. The subsampling strategy is based on the distribution of CP for each SNVs and their frequencies of being called by individual methods. Then the CCMs are only computed for the sampled SNVs. This sampling strategy is preferred over a random uniform sampling as we need to ensure the subsampled distribution of CPs to represent the full distribution, so that clusters of smaller sizes are less likely to be missed due to down sampling. Besides, the ordered sampling based on frequencies of being called repeatedly guarantee that the maximum information from each method is used.

##### 2.2.1.2 Chic (CICC – Cluster-ID-based Consensus Clustering)

Let $a_i^{(h)}, h \in \{1, \dots, H\}$, and $i \in \{1, \dots, V\}$ denote the cluster ID/label of the $i$-th SNV assigned by method $h$, and $H$ denote the total number of methods and $V$ the total number of SNVs assigned by at least one method. If the $i$-th SNV was assigned by method $h$, $a_i^{(h)}$ takes value from $\{1, \dots, K_h\}$, where $K_h$ is the number of clusters identified by method $h$; if not assigned it takes value of $-1$.
For the $i$-th SNV, we write the vector of assignment

$$\boldsymbol{A}_i = (a_i^{(1)}, a_i^{(2)}, \dots, a_i^{(H)})$$

If for $i, j \in \{1, \dots, V\}, \boldsymbol{A}_i = \boldsymbol{A}_j$, SNV $i$ and $j$ were assigned to the same cluster by all methods. There will be a limited number of distinct $\boldsymbol{A_i}'$s, let us denote these unique assignment vectors as $\boldsymbol{U}_l, l \in \{1, \dots, L\}$, and denote $N_l, l \in \{1, \dots, L\}$ as the numbers of SNVs having the unique assignment vectors, respectively.

For each pair of $\boldsymbol{U}_i$ and $\boldsymbol{U}_j, i \neq j \in \{1,2, \dots, L\}$, we define a distance between them as:

$$d(\boldsymbol{U}_i, \boldsymbol{U}_j) = \sum_{h=1}^{H} d\left(a_i^{(h)}, a_j^{(h)}\right),$$

Where

$$d\left(a_i^{(h)}, a_j^{(h)}\right) = \begin{cases} 0, & a_i^{(h)} = a_j^{(h)} \; and \; a_i^{(h)} \neq -1, a_j^{(h)} \neq -1 \\ 1, & a_i^{(h)} \neq a_j^{(h)} \; or \; a_i^{(h)} = a_j^{(h)} = -1 \end{cases}$$

An extended distance between $\boldsymbol{U}_i$ and $\boldsymbol{U}_j$ is defined as

$$d_{ij} = 10 \max_{j \in \{1, \dots L\}} N_j d(\boldsymbol{U}_i, \boldsymbol{U}_j) - \max(N_i, N_j)$$

$D = (d_{ij})$ was used to quantify the assignment distance between $\boldsymbol{U}_i$ and $\boldsymbol{U}_j$, and a hierarchical clustering with Ward's criterion was performed on $D$ to obtain the dendrogram.

If $K_{max} \coloneqq median(K_i) < 3$, where $K_i$ is the number of clusters for method $h$, we set $K_{opt} = K_{max}$, where $K_{opt}$ is the optimal number of output clusters. Otherwise, the Proportion of Ambiguous Clusters (PAC) (Şenbabaoğlu et al., 2014) was used to obtain $K_{opt}$. First, $H$ methods were sampled with replacement from the original methods. For each $k \in \{2, \dots, K_{max}\}$ we repeated the procedure 100 times to derive a consensus co-clustering matrix $M_k$ and set $K_{opt} = \text{argmin}_k \, PAC(M_k)$.

We then cut the original dendrogram to obtain $K_{opt}$ clusters of $\boldsymbol{U}_i'$s. SNVs were assigned to the same cluster if their corresponding assignment vector $\boldsymbol{U}'$s were clustered together.

The CCF of each consensus cluster was calculated as the median of the consensus CCFs of the mutations assigned to that cluster. The consensus CCF of a mutation is computed as

$$CCF_i = \frac{p_i}{\rho m_i}\left(\rho C_i^{(T)} + (1-\rho) C_i^{(N)}\right)$$

where $p_i$ is the variant allelic fraction for SNV $i$, $\rho$ is the consensus purity, $C_i^{(T)}$ is the consensus number of clonal copies, $C_i^{(N)}$ is the number of copies in the normal tissues and it is fixed to the diploid state, i.e. $C_i^{(N)} = 2$ in our study, and the multiplicity is the floor of median of the multiplicities given by individual methods: $m_i = \left\lfloor median\left\{m_i^{(1)}, \dots, m_i^{(H)}\right\}\right\rfloor$. The cellular prevalence (CP) of the cluster is $CP_i = \rho CCF_i$.

##### 2.2.1.3 Weme (WM – Weighted Median)

The Weighted Median (WeMe) method takes as input (i) a set of different *clusterings* of 1-d data and (ii) the proportion of outlier clusterings ($r$%) and it outputs a consensus clustering of the 1-d data with $K$ clusters where $K$ is a typical number of clusters in the set of clusterings. This consensus clustering has the property that, among all clusterings with at most $K$ clusters, it minimizes the earth mover distance (EMD) to the *median clustering* of the $(100 - r)\%$ inlier clusterings. The median clustering of a set of clusterings is a clustering that minimizes the sum of the EMDs from it to all members of the set, however, in general, this clustering has many more clusters than $K$. Below we define the term, *clustering* of 1-d data, formally.

A *clustering* of 1-d data is a set of ordered pairs $\{(p_k, f_k\}$, $k = 1, \dots, K_h$, where $p_k$ is the proportion of data objects assigned to cluster $k$, and $f_k$ is the location of cluster $k$ in the 1-d scale. In this study, $f_k$ corresponds to the cellular prevalence of cluster (i.e. lineage) $k$ and $p_k$ the proportion of the SNVs assigned to that cluster. $K_h$ is the number of clusters of method $h$, this number can vary between the different clusterings of the data objects (e.g. SNVs in this study).

Here, each clustering is derived from a subclonal architecture summary generated by an individual reconstruction algorithm. These summaries provide for each subclonal lineage (i.e. cluster): a cellular prevalence of the lineage $k$, $f_k$, and the fraction of mutations assigned to that lineage, $p_k$. Importantly, Weme does not include any permutation information.

To justify and explain the WeMe algorithm, we first note that the EMD between any two clusterings in 1-d is equal to the integral of the absolute difference between their corresponding cumulative distribution functions (CDF). Specifically, the EMD between clusterings $i$ and $j$, $i, j \in \{1,2, \dots, H\}$ whose CDFs are $F_i(x)$ and $F_j(x)$, respectively is

$$EMD(F_i, F_j) = \int_{-\infty}^{\infty} |F_i(x) - F_j(x)| \, dx$$

Where the $F_i(x)$ of a clustering, $\{(p_k, f_k\}$, $k = 1, \dots, K_h$, is:

$$F(x) = \sum_{i \in \{k | f_k \le x\}} p_i$$

Note, as indicated above, that the CDF, $F(x)$, corresponding to a clustering is a piecewise constant function of $x$, with the discontinuities occurring at the values of $f_k$ (Fig. 19 – top left). At these discontinuities, the y-value of the CDF jumps by $p_k$.

Because of this, it is easy to compute the EMD between two methods, namely $i$ and $j$. We just need to compute the area of, at most, $k_i + k_j$ different rectangles, where $k_i$ and $k_j$ are the number of clusters in the $i$- and $j$-th clusterings, respectively. These rectangles are delineated by the union of the cluster locations of the two clusterings - these are the discontinuities in each individual CDF (Fig. 19 – top right).

We defined an unconstrained consensus clustering for a set of clusterings $\{1,2,\dots,H\}$ as a new clustering $H^*$, $\{(p_k^*, f_k^*\}$, that minimizes the sum of the EMDs to all other clusterings, i.e.

$$\text{argmin}_{\{(p_k^*, f_k^*)\}} \sum_{h=1}^{H} EMD(F_h, F_{H^*})$$

It is straightforward to prove that clustering $H^*$ has clusters at each location in the union of the cluster locations from the set of clusterings, and these clusters have their proportions set so that

$$F_{H^*}(x) = median(F_h), \text{for } h = 1,\dots,H$$

Fig. 19 (bottom left) displays a visual proof of this for $H = 3$. Because $H^*$ is defined using the median of the CDFs for the other clusterings, we use the term *median clustering* to refer to the unconstrained consensus clustering $H^*$. However, that the median clustering can have many more cluster locations than a typical clustering in the set. We would prefer a consensus clustering that also has a number of clusters that is typical for the set of clusterings. As such, we define the *$K$-constrained median* clustering as the clustering $H^W$, $\{(p_k^W, f_k^W\}$, with no more than $K$ cluster locations that minimizes $EMD(H^*, H^W)$. Note that, in general, $H^W$ need not have cluster locations corresponding to any of those in the median clustering $H^*$. We will describe how to select $K$ later, and for now, assume $K$ is known.

To find the $K$-constrained median clustering $H^W$, we first compute the median clustering, $H^*$. Then we perform a grid search over possible settings of a vector $\boldsymbol{p}$ of length $K$ whose elements represent the proportions of objects assigned to each of the $K$ clusters. On the CDF, $p_k$ is the jump at the $k$-th discontinuity. Given $\boldsymbol{p}$, we then set the corresponding cluster locations, $\boldsymbol{f}$, to minimize $EMD(H^*, H^W)$. We can minimize this by taking the weighted median of the cluster locations of $F_{H^*}(x)$ within the $y$-range of the step (Fig. 19 – bottom right). We then set the $K$-constrained median clustering to be the one that minimizes this minimal EMD over all settings of $\boldsymbol{p}$.

To define a robust WeMe consensus, we remove outlier clusterings, and compute the $K$-constrained median clustering only on the remaining, *inlier* clusterings. To identify the outliers, we do the following:
We compute the median clustering of all of the methods
We remove the reconstructions with the highest $r\%$ of EMDs to the median clustering (rounded down)

Then WeMe recomputes the median clustering using only the remaining clusterings, and computes the $K$-median clustering based on the new median clustering, setting $K$ to be the ceiling of the median of the remaining clusterings.

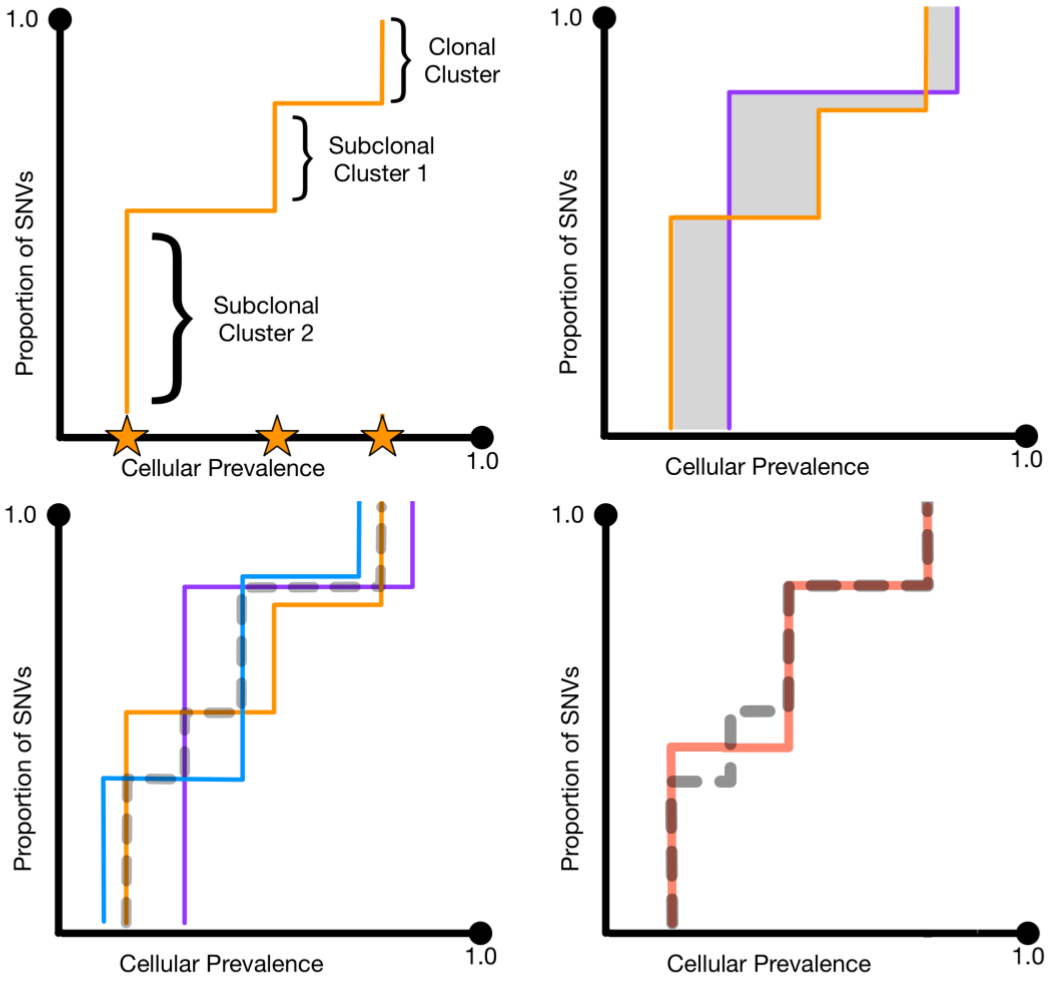


*Fig. 19* ***Illustration of the WeMe algorithm***. *(Top left) CDF representation of a 1-d "clustering". Stars indicate the cellular prevalence of the cluster centers; height of the CDF jumps corresponds to the proportion of mutations assigned to that cluster. (Top right) Two CDFs (orange and purple lines) indicate different clusterings of the same sample. Total area of grey region is equal to the Earth Mover Distance (EMD), also known as Wasserstein distance, between the two clusterings. (Bottom left) Three solid lines (orange, blue, purple) are CDFs for three different clusterings. The median clustering (dashed grey line) corresponds to the CDF made by taking the median in the x-axis direction of the three CDFs for each y-value. Note that the median clustering has four cluster centers. (Bottom right) The weighted median clustering corresponds to the CDF (red line) with the desired number of centers (i.e., jumps) that minimizes the EMD to the CDF for the median clustering (dashed grey line).*

###### 2.2.1.4 Three random methods

The subclonal architecture summary metrics described below can be used to assess the performance of a subclonal architecture caller, where low scores mean a method is performing well. Often methods will not obtain the perfect solution and therefore their scores will deviate from the perfect score. What is not clear however is what performance can be reasonably expected. We reasoned that a caller should be able to outperform a simple random method and that running a method that yields a random subclonal architecture on the same data as the caller would then provide an upper bound of what can be considered reasonable performance. To this end we have developed two simple methods that generate subclonal reconstructions with varying degrees of randomness and one method that returns only clonal tumors.

**1 - Stick breaking**
The stick breaking method starts with drawing a random number between 0 and 6 to determine the number of clusters. It then orders the mutations by their CCF and iteratively breaks a randomly sized chunk of the ordered mutations. Each of these chunks represents a mutation cluster and its location is obtained by taking the mean CCF of the mutations in the chunk. Mutations are automatically assigned as they belong to a particular chunk. The advantage of this approach is that it is more likely to place a cluster where there are large real clusters, but it does also tend to place multiple clusters within large real clusters.

**2 - Informed**
The downside of the stick breaking approach is that it performs a single series of breaks and returns that as a solution. We wondered whether the method could be improved by selecting the best solution from a series of random models. The informed method runs the stick breaking implementation described above 100 times. Contrary to the above approach, the informed method records the size and locations of clusters, but does not record the assignments. It applies the MutationTimer approach (Gerstung et al., 2020), which models each mutation cluster as a beta-binomial and takes into account the size of the cluster. MutationTimer then calculates the proportion of mutations that are poorly explained (i.e. fall in the outermost 5% of the beta-binomial distributions). This proportion is calculated for all 100 runs, after which the run that yields the lowest value is selected as the returned subclonal architecture.

**3 - Single cluster**
This approach places a single cluster to explain the data. It can obtain the single cluster by taking the mean mutation CCF, or it can be forced to place a clonal cluster at a CCF of 1. All mutations are assigned to the single cluster.

#### 2.2.2 Evaluation metrics

For all methods, we evaluated two aspects of the solutions: scoring against the ground truth and the comparison of similarities between pairs of methods. The scoring system is used to assess the performance of all individual and consensus methods in the simulated data, whereas the similarity measures are used to compare the relative performance across methods on the PCAWG data, where there is no ground truth.

**2.2.2.1 Performance on simulated data**

We started using conventional evaluation metrics for clustering that are based on assignments, e.g. rand index and mutual information. However, not only are these metrics not consistent and have their own flaws (Kleinberg, 2003; Vinh et al., 2010; Wagner and Wagner, 2007), but also, in our setting, they turned out to be impractical. Indeed, not all individual methods assigned the same subset of SNVs to construct the subclonal structure. Importantly, the ultimate goal of the consensus methods was to identify the subclonal structure, i.e. number of clusters, number of SNVs within them and cellular prevalence (CP) for each cluster, whereas the assignments of SNVs, Indels, and SVs were performed post hoc by a uniform assignment method.

Therefore, we designed three metrics specifically for this study to measure the performance of the clustering results. These metrics reflected the important biological aspects of the results:

1) **Relative clonal fraction change**, characterizing the proportion of clonal events, is defined as

$$CFC = 1 - \frac{|F_O - F_T|}{F_T}$$

where $F_O$ is the clonal fraction reported by the method, and $F_T$ the true clonal fraction.

2) **Relative Number of clusters difference**, representing the relative subclonal events frequency, has the form of

$$NCD = 1 - \left| e^{\frac{C_O - C_T}{C_T}} - 1 \right|$$

where $C_O$ is the number of clusters from each method, and $C_T$ is the true number of clusters. This metric favored slightly underestimation over overestimation, which was designed to accompany the fact that the PCAWG working group project was reporting a lower bound of number subclones.

3) **Root mean square error,** reflecting overall architecture of subclonal composition, is calculated as

$$RMSE = 1 - \frac{\sqrt{\frac{1}{N}\sum_{n=1}^{N}(\phi_n^O - \phi_n^T)^2}}{\rho}$$

where $\phi_n^O$ is the method reported CP of SNV $n$, $\phi_n^T$ the true CP of SNV $n$, and $\rho$ the purity of the sample.

All metrics are rescaled so that 0 is the worst score and 1 the best. To summarize the metrics into a final scores, a normalized rank approach is used, where the sum of ranks of each metric of method $h$, $r_i^{(h)}$, in sample $i$ is normalized as

$$R_i^{(h)} = \frac{r_i^{(h)} - \min_h\{r_i^{(h)}\}}{\max_h\{r_i^{(h)}\} - \min_h\{r_i^{(h)}\}}$$

so that the best performing method is ranked as 1, and the worst 0.

###### 2.2.2.1 Similarity in real data

We compared pairs of methods to assess the similarities of the solutions within PCAWG and simulated data. We used normalized difference over average to show the relative consistency of these consensus methods on PCAWG data, and investigate the robustness of consensus methods compared to each individual method.

The normalized differences are defined as follows, where $1 \leq i < j \leq H$, and $H$ is the number of all methods including consensus methods:

**1) The fraction of clonal SNVs:**

$$CF_{similarity} = 1 - \frac{2|F_i - F_j|}{F_i + F_j}$$

where $F_i$ denotes the clonal fraction of the $i$-th method.

**2) The number of clusters:**

$$NC_{similarity} = 1 - \frac{2|N_i - N_j|}{N_i + N_j}$$

where $N_i$ is the number of clusters determined by the $i$-th method.

**3) The RMSE:**

$$RMSE_{similarity} = 1 - \frac{\sqrt{\frac{1}{\mathrm{N}} \sum_{n=1}^{N} \left( \phi_n^{(i)} - \phi_n^{(j)} \right)^2}}{\rho}$$

where $\phi_n^{(i)}$ is the $i$-th method reported CP of SNV $n$, and $\rho$ is the purity of the sample.

To account for diversities in dynamic range among samples, a standardization step: M-min(M)) / (max(M) - min(M), where M is a metric, was conducted to ensure each metric is distributed within interval [0,1].

#### 2.2.3 Results

The 11 subclonal structure identification methods relied on different assumptions and clustering mechanisms (Fig. 20).

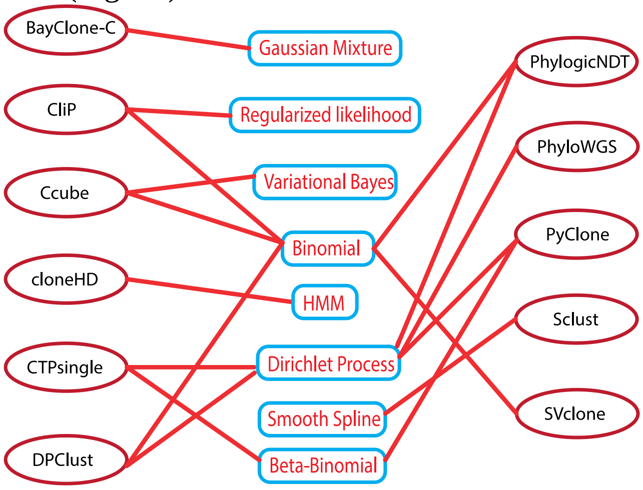


*Fig. 20 Individual methods and their internal assumptions and models. Each method typically does not share assumptions/mechanisms with other methods. Even if they do, like PhylogicNDT and DPClust, they utilize the mechanism differently and there are substantial differences in pre-/post- processing.*

These differences explain why each method leads to different results and tends to perform better on a different subset of samples (Fig. 21).

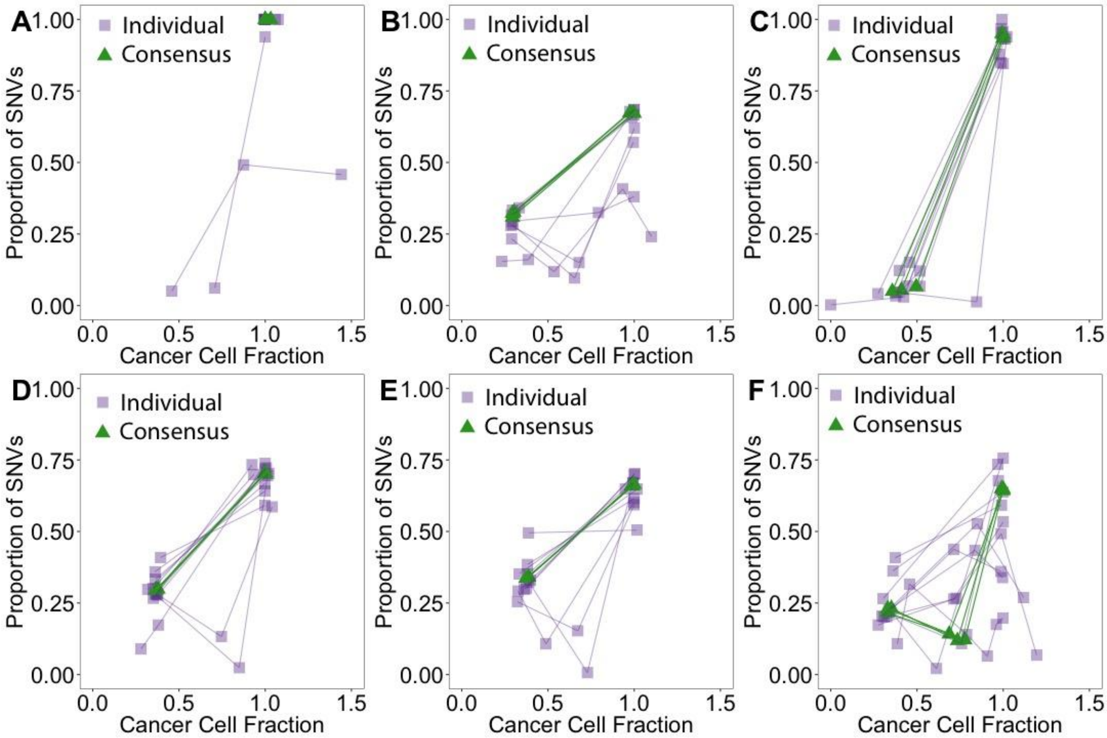


*Fig. 21 Examples of subclonal reconstruction differences across methods. Results across methods are variable, but the consensus methods are able to reconstruct the subclonal structure robustly. Each individual method's subclonal peaks are linked with a line. A) Most solutions agree except two. B) Methods agree on the clonal location and size, but they diverge for the subclonal position and size. C-E) Global agreement on the number of clusters*

*but deviation in position and size. F) High variability and disagreement across methods still leads to stable consensus solutions.*

This motivated the design of consensus methods that might combine the strengths of individual methods and be more robust across various tumor subclonal topologies (e.g. Fig. 21F).

To assess the performance of the methods, two independent simulation sets were constructed. The first set, PhylogicSim500, contained 500 samples, where the copy number profiles were sampled from the PCAWG samples and the other parameters were independently sampled from fixed distributions (**2.4.1 Simulation of subclonally heterogeneous samples – PhylogicSim500**). To allow individual methods to debug and tune their parameters, the true subclonal architectures were released along with the PhylogicSim data. The second set, SimClone1000, contained 965 samples and used a grid design to cover the collection of scenarios encountered in the PCAWG samples. The ground truth of SimClone1000 data was not released before the final evaluation of all methods; the design was also kept hidden and made the inference of the grid difficult.

As described in **2.2.2 Evaluation metrics**, to assess the performance of all the methods, we established an evaluation system using high-level summary metrics, corresponding to the biological variable of interest:

1) clonal fractions, which describes how well a method is able to estimate the fraction of (sub)clonal SNVs;
2) number of clusters, which is essential in clustering analysis and represented how well each method identifies the number of distinct subclones; and
3) root mean squared error (RMSE) of mutation assignments, which characterizes how well the overall subclonal architecture matches another architecture (see supplementary methods).

To gain an overall performance assessment, the total score for a given method in a given sample was obtained by averaging the ranks of the three metrics in the sample for this method.

**Individual methods lead to variable results while consensus methods lead to more robust and accurate results in simulated data**

The total scores (Fig. 22) show that the consensus methods were ranked at the top, together with the best individual methods. There are large variations in performance across individual methods within and across two simulation settings. However, we observed the two simulation sets yielded similar results with the consensus methods, which confirmed that the consensus methods achieved robust and consistent results as compared to individual methods.

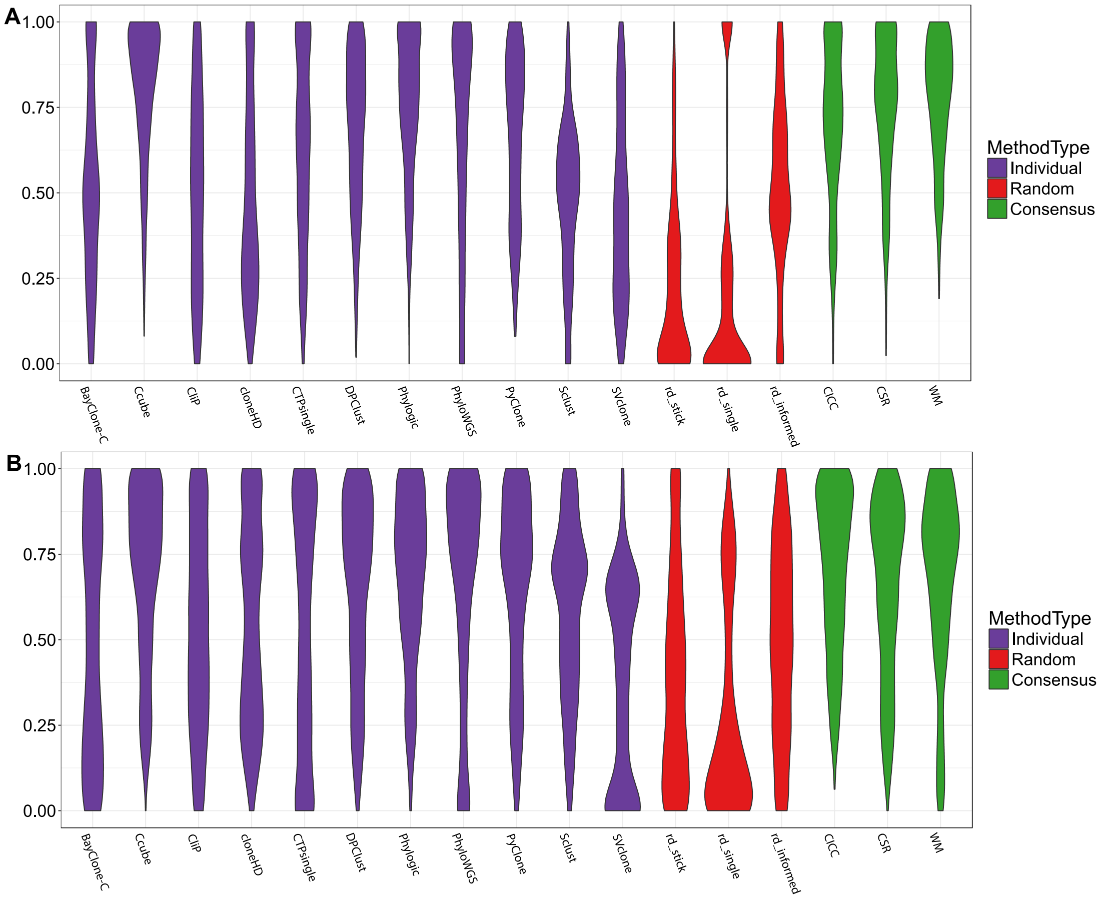


*Fig. 22 The overall scores of all individual (purple), random (red), consensus (green) subclonal architecture reconstruction methods on A) PhylogicSim500 and B) SimClone1000.*

In the detailed comparison of the three metrics themselves, each individual method topped a metric in a fraction of the samples but not all. Whereas some individual methods were globally significantly better than others on the simulated data, the scores of the consensus methods were not significantly lower than those of individual methods on both datasets - between the best consensus and the best individual method the Mann Whitney U tests fdr=0.029 for PhylogicSim500, and fdr=0.336 for SimClone1000.

Given the design of the SimClone1000 simulations, we could ask how parameters such as true number of subclones and the number of reads per chromosome copy (nrpcc) influenced the absolute and relative performances of the methods (Fig. 23). As expected we observed: an improvement of the performance with increasing power (nrpcc); that CliP systematically overestimated the number of subclones when there were none; and that most methods performed well at integrating copy number changes, except from CTPsingle, whose relative performance significantly improved in fully diploid tumors (not shown). Most importantly, the consensus methods performed competitively across the grid of nrpcc and number of subclones (Fig. 23), further suggesting that they combine the strengths of individual methods. This was also true when we looked at performance only in samples for which the scores for number of subclones were equal, demonstrating that the high performance of the consensus did not stem only from taking the median number of subclones across individual methods (not shown).

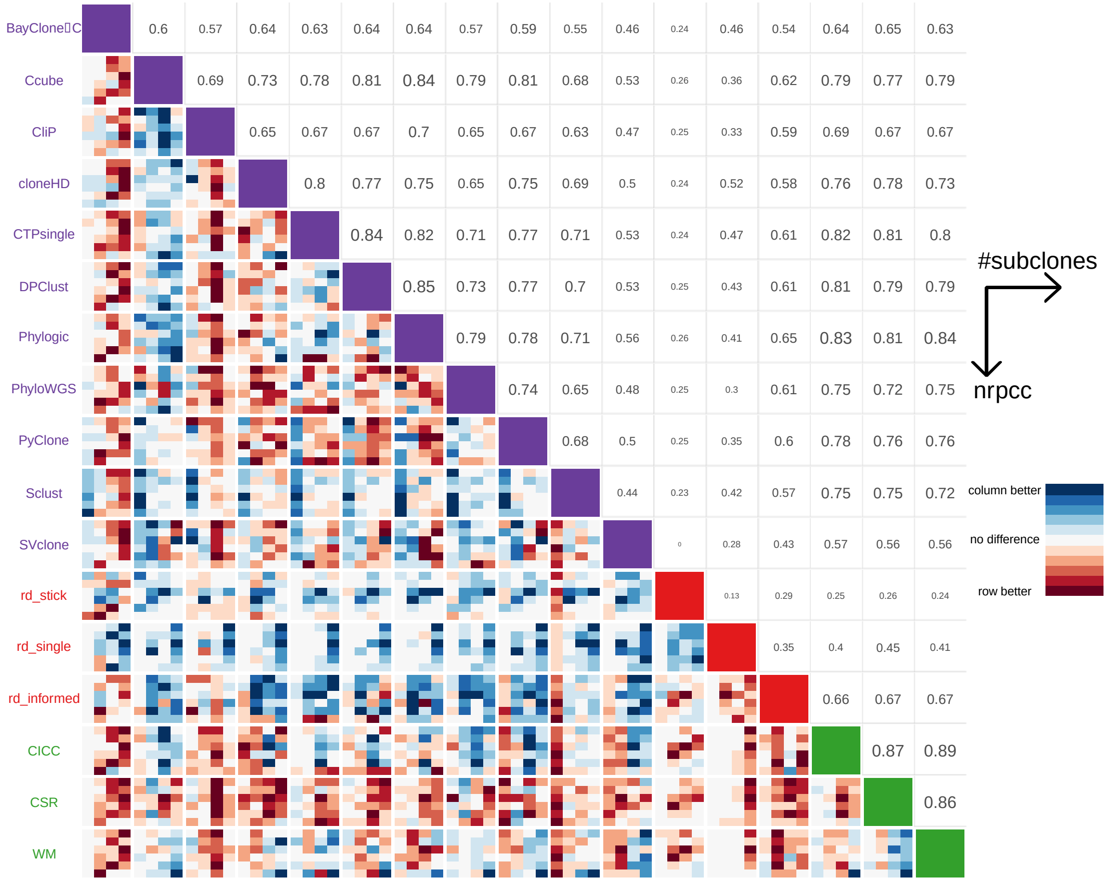


*Fig. 23 Upper triangle: relative similarities between methods (individual methods are in purple, random methods are in red and consensus methods are in green). The size and transparency scale linearly with the similarity score. Lower triangle: heatmaps of relative ranks across simulations according to two grid parameters: number of subclones on the x-axis of each heatmap, nrpcc on the y-axis of each heatmap (the legend for these axes is on the top right). The scores are average across all simulations falling into the grid cell and a minmax normalization is then applied across the grid values, cells are then colored from red (method on the row is better) to blue (method on the column is better) with the intensity of the color scaling with the normalized score.*

#### Top performing methods lead to more similar results in both simulations sets and real data, suggesting that their performance remains the same in real data

Top performers displayed higher similarity (Fig. 24). Whereas individual method's performance varied across independent simulation sets, the consensus methods consistently showed higher similarities than individual methods. Interestingly, individual methods were more similar to the consensus methods than they were to each other.

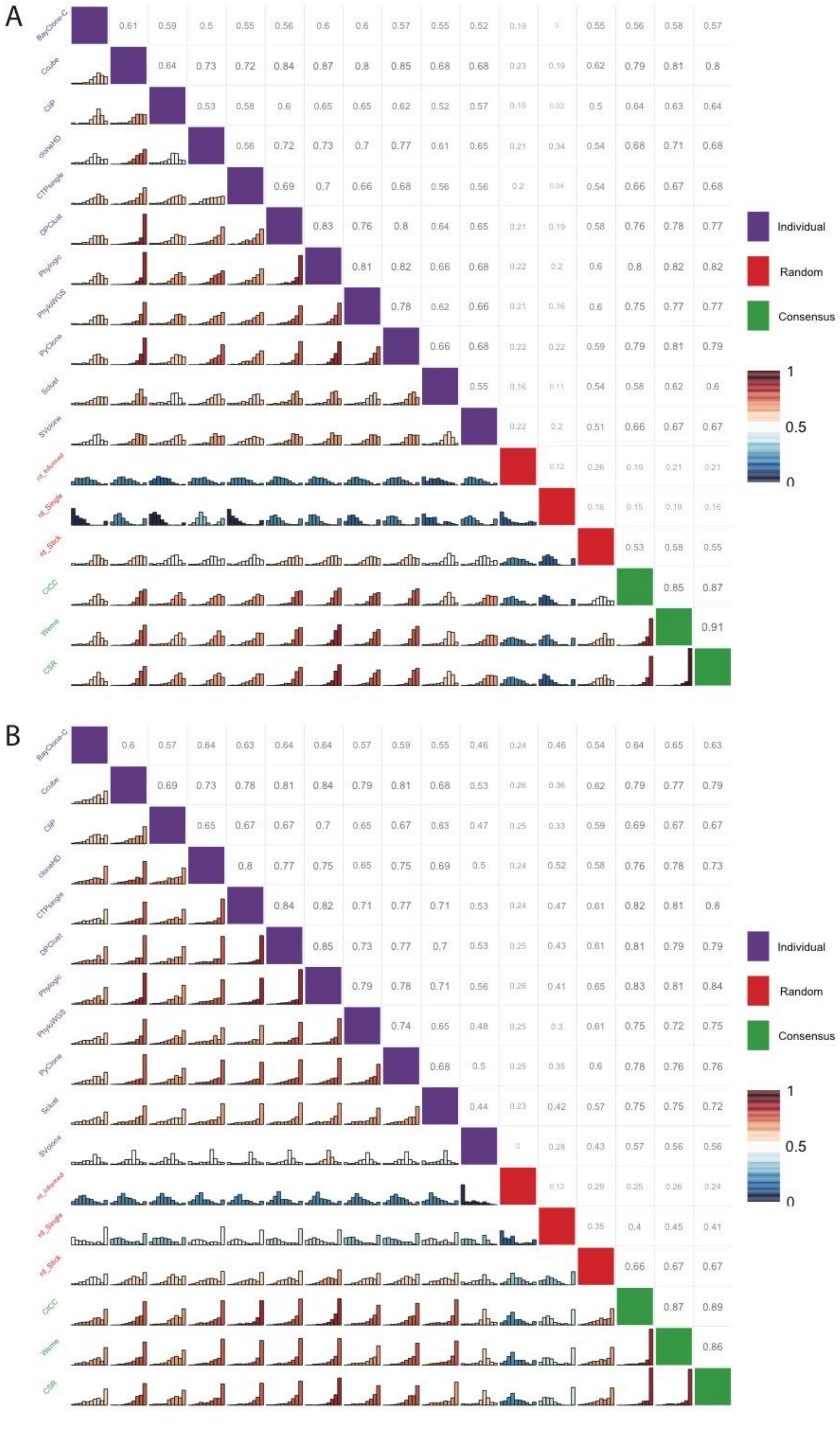
A
B
BayClone-C
Ccube
CliP
cloneHD
CTPsingle
DPClust
Phylogic
PhyloWGS
PyClone
Sclust
SVclone
rd_informed
rd_Single
rd_Slick
CICC
Weme
CSR
Individual
Random
Consensus
1
0.5
0

*Fig. 24 Pairwise distributions of the similarities across samples (lower triangle; bar width was set to 10%) and median similarities across samples (upper triangle) for A) PhylogicSim500 and B) SimClone1000.*

As expected, the consensus methods were the most similar to each other. Few pairs of methods showed high similarity, although not as consistent across the two datasets as the three consensus methods. As desired, the random methods had a quite uniform distribution of similarities against any method.

**Running on more methods leads to better results**

We assessed the performance of the consensus methods running on 2 to 11 input individual methods. We ran CSR, WM, and CICC on all 965 simulated tumors from SimClone1000 for all of the 2,035 combinations of input methods, i.e. 1,963,775 times. This showed that performance increased with number of methods taken as input, suggesting a consensus of all methods is guaranteed to provide results at a median-ranked performance, and safeguarded from low performance. (Fig. 25).

Integrating more methods into the consensus improved the lower bound and the median of the median performances. Few combinations of a subset of the methods significantly outperformed the individual methods on this dataset. These particular combinations, however, would be dataset-specific and not generalizable to real PCAWG data.

**Conclusion**

In the absence of ground truth the high similarities between consensus methods in real PCAWG and in simulated samples, together with their consistently high performance on simulated data, suggested robust accurate reconstructions through consensus clustering on PCAWG data. Each individual method showed high performance across a range of scenarios and added to stabilize the consensus results.

Therefore, we applied the consensus approaches to the outputs of the 11 subclonal structure reconstruction methods on 2,778 PCAWG tumor samples and settled on using the output of one of the consensus methods, WeMe – this choice of consensus approach was arbitrary, as all three led to similar results.

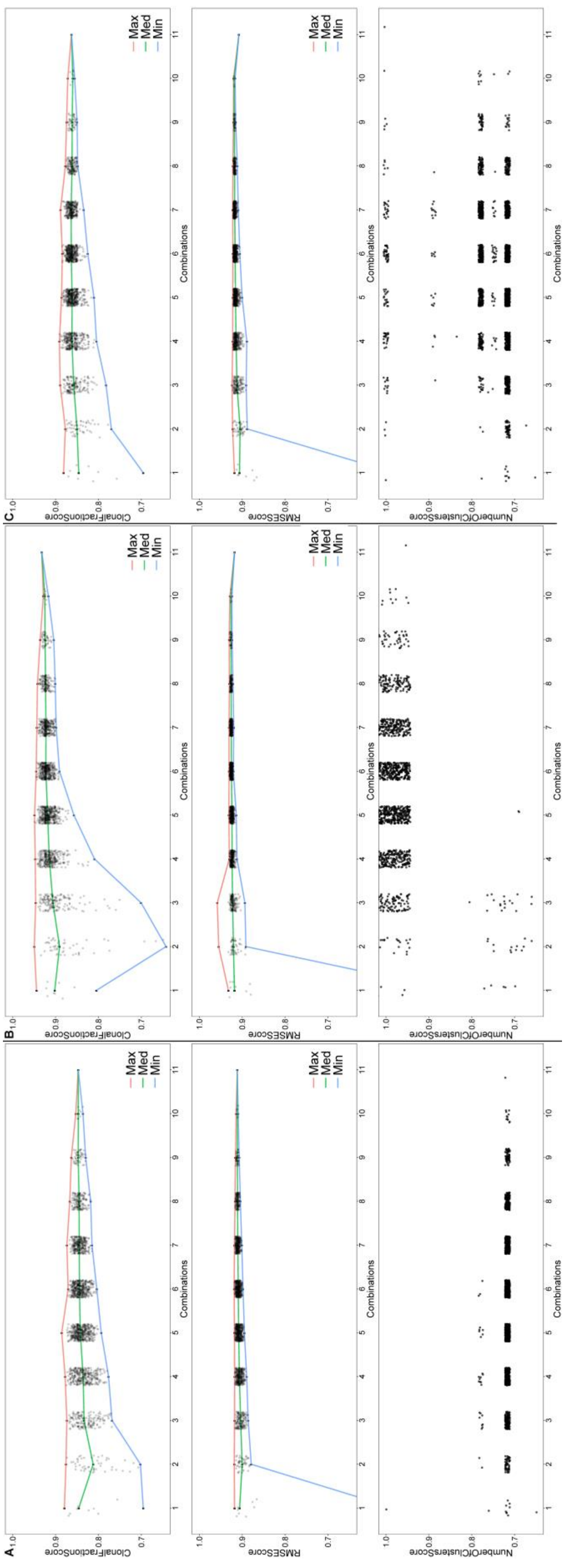
A
B
C
ClonalFractionScore
RMSEScore
NumberOfClustersScore
Combinations
Max
Med
Min

*Fig. 25 Median performances of consensus methods on SimClone1000 with different numbers of methods included, from 1 (individual methods) to 11 (the reported consensus results), for clonal fraction, number of subclones and RMSE, from top to bottom. The green line connects the medians of the medians; the red and blue connect the extreme top and bottom values, respectively. Consensus was obtained using CSR (A), CICC on a subset of 680 out of 965 samples (B) or WM (C).*

### 2.3 “Winner’s curse” correction

Somatic mutation callers require several high-quality supporting reads to reliably call a mutation. In samples with low purity and/or low allele fraction subclones, the observed allele fraction and cancer cell fraction (CCF) of the clones will be biased upwards, since the signal from allele count fluctuations that fall below the caller’s detection limit will be unaccounted for (variants are not detected). If subclonal reconstruction is done without correcting for of such biases, the lower CCF cluster positions would be inaccurate and subclonal cluster sizes (number of mutations) would be significantly underestimated. This effect has been dubbed the “Winner's curse” (WC).

We wish to correct for the effects of WC on both the inferred CCF of identified subclonal lineages and the number of SNVs belonging to that lineage. For this purpose, we have developed two orthogonal methods as described below.

#### 2.3.1 PhylogicCorrectBias

Ignaty Leshchiner, Dimitri Livitz, Gad Getz

To measure and correct the bias caused by the WC effect, we employed a simulation-based search method, PhylogicCorrectBias (a component of the PhylogicNDT package), that empirically estimates the correct CCF cluster position and its mutational composition. In case there is no bias, the CCF of the simulated clusters will match that of the detected sample.

To perform the simulation, a potentially truncated CCF cluster of mutations is investigated as follows. Simulated mutations are randomly placed on an allele along the genome according to the copy number profile and bases at risk at a chromosome/allele, while their expected allele fraction is selected according to the current cluster CCF and local coverage. Subsequently, the algorithm defines the measured number of alternative and reference reads given the expected allele fraction according to a binomial distribution and decides if the mutation is “detected” based on the filtering and detection criteria of the callers used. The simulated cluster CCF is then calculated from the “detected” mutations. In the event the values of the simulated and experimentally measured clusters do not match, the algorithm iteratively lowers the real cluster CCF downwards in 0.01 increments (maximum CCF resolution) until the “detected” cluster CCF and mutation number match the experimentally measured cluster position, within 0.01 CCF delta. On convergence of the algorithm the corrected cluster position and expected mutation count is reported.

#### 2.3.2 SpoilSport

Amit Deshwar, Quaid D. Morris

To correct for the WC, we make some simplifying assumptions about how the cellular prevalence (CP) estimates for each lineage were derived. Specifically, we assume that the CPs were maximum likelihood estimates under a binomial sampling model where all variants are at average read depth and exist as a single copy in a region of copy number equal to the average ploidy of the tumor. To estimate the magnitude of the WC effect we first convert from CP to a probability of observing a variant read using the following equation:

$$p_v = \frac{\theta}{\rho\,\psi + (1 - \rho)2}$$

Where $\theta$ is the CP of the lineage, $\rho$ is the purity of the tumor and $\psi$ is the tumor ploidy. The denominator of this equation is the average copy number in the sample.
Given $p_v$, we use moment-matching to find the truncated binomial distribution with mean observed success probability equal to $p_v$. The PDF of the truncated binomial distribution truncated at T successes is:

$$\text{TruncBinom}(x,p,n,T) = \begin{cases} 0, & x \le T \\ \dfrac{\text{Binomial}(x,p,n)}{\sum_{j=0}^{T} \text{Binomial}(j,p,n)}, & x > T \end{cases}$$

There is no closed form equation for the mean but it can be calculated conditioned on $T$, $p$ and $n$ by summing over values of $x$ from 0 to $n$:

$$E[\text{TruncBinom}(p,n,T)] = \sum_{x=0}^{n} x\ \text{TruncBinom}(x,p,n,T)$$

We set $n$ to equal the rounded mean read depth of variants in the lineage we are correcting and find $p$ by solving the following optimization problem:

$$p^* = \underset{p}{\text{argmin}} |E[\text{TruncBinom}(p,n,T)] - p_v|$$

We solve this problem using the bounded *localmin* procedure as implemented in the R function *optimize*.
After finding $p*$ we convert this back to CP by multiplying by the average copy number. This is the WC corrected CP used for downstream analysis.
We additionally infer the ratio of true variants to observed variants (sf) by inverting the untruncated proportion of the probability mass of the binomial distribution:

$$sf = \frac{1}{1 - \sum_{j=0}^{T} \text{Binomial}(j,p^*,n)}$$

##### 2.3.3 “Winner’s curse” correction consensus

Ignaty Leshchiner, Amit Deshwar, Dimitri Livitz, Quaid D. Morris, Gad Getz

Consensus correction for the "winner's curse"-like effect was produced by integrating the values reported by the two methods above. Consensus CCF shift correction was obtained as the mean between the reported values, while for the corrected number of mutations in the subclone a geometric average was used. In a small set of samples a most conservative solution was used when available.

#### 2.4 Validation

##### 2.4.1 Simulation of subclonally heterogeneous samples – PhylogicSim500

Ignaty Leshchiner, Daniel Rosebrock, Dimitri Livitz, Gad Getz

To generate simulated data, we followed specific guidelines to ensure biologically realistic and heterogeneous data. The software simulator, PhylogicSim, (another component of the PhylogicNDT package) was used to generate 500 samples with diverse heterogeneity profiles. First, we randomly generated cluster CCFs, as well as the number of mutations in each cluster. A clonal cluster was included in each simulated sample. The number of subclones (N) in each sample was chosen randomly from the set (0, 1, 2, 3, 4) for samples with linear evolution, and (2, 3, 4) for samples with branched evolution, where samples with branched evolution always contained two dominant sibling clones whose CCFs summed to 1. Cluster CCFs were furthermore chosen to have a distance of 0.1 from the nearest cluster CCF, and to be at least 0.1 in magnitude. The number of mutations in each sample was chosen from a uniform distribution with domain 2,000 - 7,000, U[2000,7000]. The proportion of mutations assigned to each cluster was then estimated by taking N random numbers from U[0,1], dividing by the sum, and assigning each of these proportions randomly to a cluster.

Samples were chosen by inspecting the PCAWG consensus copy number profiles across all cancer types, selecting tumors where at least 70% of the genome was covered by clonal 3-star segments (N=853 samples). We then used the ABSOLUTE output as the copy number profile for the simulated tumor. Each segment with a subclonal allelic copy number state was rounded to the nearest clonal copy number state. Segments were then assigned randomly to subclones in the same CCF space as the mutation clusters. For each sample, purity was chosen from the distribution of consensus purities across all called PCAWG samples.

We used a beta binomial model to simulate coverage profiles (number of reads at each locus in the genome) for each simulated sample. To do this, all normal (non-tumor) coverage profiles from all PCAWG samples were fit to a beta-binomial distribution, optimizing parameters alpha and beta in the beta-binomial pdf, leaving n=500 fixed. For each simulated sample, we chose a mean coverage by sampling from the distribution of mean coverages of all tumor samples in PCAWG (total mean coverage = 53.6x). We then chose alpha, beta, and n from the beta-binomial fits of all normal coverage profiles that had the closest mean to the sampled tumor coverage. Coverage at each site in the simulation was then chosen at random from a beta-binomial distribution using these alpha, beta, and *n* parameters, scaling accordingly to account for local copy number changes.

Using the actual (true) CCF of the mutation, multiplicity, and allelic copy number information at the site of the mutation, we calculate expected allelic fraction for each SNV site. Finally, the measured variant allele count for each SNV site is chosen from a binomial distribution ~ B(n,p), where n is the coverage at the site, and p is the expected allelic fraction of the mutation.

For each simulated SNV the following information is used to calculate the expected (true) allelic fraction:

- $mult$ = multiplicity of mutation
- $CCF_{mut}$ = ccf of mutation
- $\rho$ = sample purity
- $CN_{cl}$ = local absolute copy number (alleles A+B) in clonal fraction (i.e. $1 - CCF_{scna}$)
- $CN_{sub}$= local absolute copy number (alleles A+B) in subclonal fraction
- $CCF_{scna}$ = ccf for subclonal copy number event

To calculate the expected (true) allelic fraction we used the following measurements.

a) in case of no sub-clonal CN alterations:

$$af = CCF_{mut}\, mult\, \rho \;/\; 2(1.0 - \rho) + \rho CN_{cl}$$

b) in case of sub-clonal CN alterations:

$$mult = mult_{sub}\, CCF_{scna} + mult_{cl}(1 - CCF_{scna})$$

$$af = \frac{CCF_{mult}\;\; mult\;\; \rho}{2(1.0 - \rho)} + CN_{sub}\, CCF_{scna}\rho + CN + CN_{cl}\, \rho(1 - CCF_{scna})$$

we allow a) $CCF_{mut} = 1$ or b) $CCF_{mut} = CCF_{scna}$ and $mult_{cl} = 0$ or c) $CCF_{mut} = 1 - CCF_{scna}$ and $mult_{sub} = 0$

We have used the above-simulated datasets for extensive quality control and validation of the individual reconstruction methods as well as the performance of the consensus subclonal reconstruction. In general, 11 subclonal reconstruction methods as well as the consensus approaches used in this manuscript produced consistent results (Fig. 26) on the simulated datasets. In some samples, specific methods deviated from the majority, justifying the consensus approaches.
As an additional metric “obtainable” truth was calculated from the true cluster positions, based only on mutations that mutation callers would discover (which results in a CCF shift of cluster positions and sizes) and by performing additional pair-wise cluster permutation significance test to identify pairs of clusters which would be statically undistinguishable from a single cluster (p-value>0.05).

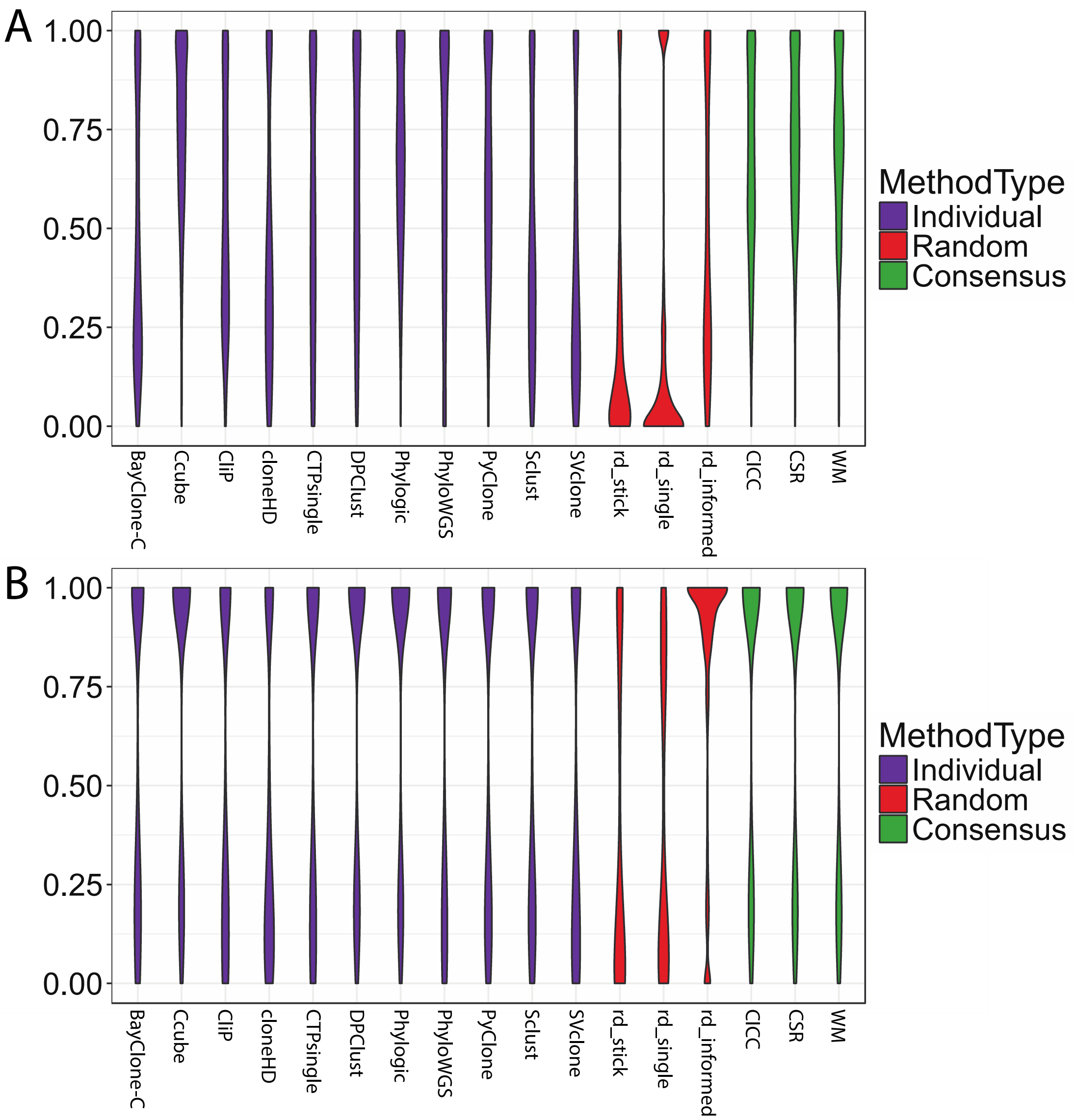


*Fig. 26 Summary of subclonal reconstruction on simulation data. General consistency between individual methods and consensus method results:* ***A)*** *fraction of clonal mutations and* ***B)*** *number of subclonal clusters.*

We have generated a set of simulated samples to specifically explore the resolution of single sample subclonal reconstruction. We have simulated sample sets with varying purities and cluster-to-cluster distance series ranging from 0.2 to 0.03 CCF inter-cluster distance. In summary, even in the purest samples, none of the methods were able to consistently resolve clusters that were less than 0.07 CCF apart. These results provide an extra argument for potential additional intra-tumor heterogeneity within subclonal reconstructions in sequenced samples, especially when subclonal CCFs overlap in single sample cases.

The simulated datasets were used to extensively benchmark and validate the “winner’s curse” correction methods. When run on the “detected” portion of simulated variants they accurately recapitulated the true CCF of the subclonal clusters and properly corrected the associated number of mutations.

#### 2.4.2 A grid approach to simulations of tumors – SimClone1000

Maxime Tarabichi, Stefan Dentro, Quaid Morris, Wenyi Wang, Moritz Gerstung, David Wedge, Peter Van Loo

In addition to the PhylogicSim500 simulations, we simulated an independent set of 700 tumors using the SimClone simulator, for which the truth was hidden. We opted for a grid approach to explore the performance of the methods across a range of values for four parameters: number of reads per chromosome copy (nrpcc), number of subclones, fraction of clonal SNVs, and number of clonal SNVs.
Additionally, we resimulated 300 of the 700 samples in a fully diploid setting with the exact simulation parameters, including nrpcc, allowing us to assess the impact of copy number aberrations on performance.

**Simulating subclonal architectures with SimClone**
We simulated subclonal architectures by first establishing a series of required parameters (see the next section) and provide these to the SimClone simulator. In this instance, SimClone takes a subclonal architecture design (number of mutation clusters, the proportions of tumor cells that each cluster takes up and the number of mutations per cluster) and a copy number profile (genome wide copy number and a purity value).

SimClone makes a number of assumptions during its simulation runs: Both the mutation and wild type alleles are supported by a number of reads. We assume that the distribution of the number of mutation-supporting-reads takes on the shape of a binomial distribution. To model variation on the total number of reads covering the locus where the mutation has occurred, we assume that the depth can be modelled through a Poisson distribution. The mutation is carried by a number of chromosome copies (multiplicity). The shape of this distribution is partly determined by the copy number profile, that bounds the possible multiplicity states, and by cancer type specific development, i.e. if gains occur late there will be many SNVs on multiple copies, while if gains occur early there will be few. Multiplicity is modelled through a Poisson, and for the SimClone1000 data set its parameter was set to 1. Finally, as a simplification, subclonal mutations cannot be carried by more or less than 1 chromosome copy.

With cluster locations and sizes provided as input, SimClone now simulates the mutations per cluster. Further input is required in a copy number profile, a tumor purity value, coverage and a multiplicity λ parameter (which is set to 1 by default). Mutations are generated by calculating the expected number of reads reporting the mutation and wild-type alleles. But the multiplicity must first be determined before those can be calculated.

The multiplicity ($m_m$) is drawn from a Poisson distribution with the provided λ parameter as input.

$$m_m \sim \text{Poisson}(\lambda)$$

The mutations are randomly assigned to a copy number segment in the provided profile. If the multiplicity is not possible given the major and minor allele of the selected segment we adjust it to the copy number of the major allele.

Then the number of reads per chromosome copy for the tumor ($c_t$) and normal ($c_n$) cells are calculated from the total coverage ($C$), tumor purity (ρ) and tumor ploidy ($\psi_t$). The total copy number of the normal cells is assumed to be 2:

$$c_t = C \frac{\rho}{\rho\psi_t + 2(1-\rho)}$$

$$c_n = C \frac{1-\rho}{\rho\psi_t + 2(1-\rho)}$$

The expected number of mutant alleles $r_m$ is determined by the multiplicity of the mutation, the mutation's fraction of tumor cells (*f*) and the number of reads per tumor chromosome copy ($c_t$):

$$\mathbb{E}(r_m) = m_m f c_t$$

The expected number of wild type alleles $r_w$ consists of three components: (1) Reads from normal cells (can be zero when the sample is pure and does not contain normal cells), (2) reads from whole chromosome copies from tumor cells that are not carrying the mutation (can also be zero when the copy number is 1+0) and (3) if the mutation is subclonal, an additional number of reads from cells that are not part of the subclone that carries the mutation:

$$\mathbb{E}(r_w) = 2c_n + m_w c_t + m_m(1-f)c_t$$

The total number of observed reads is then drawn from a Poisson distribution:

$$r_d \sim \text{Poisson}(\mathbb{E}(r_m) + \mathbb{E}(r_w))$$

And the final mutant and wild type alleles are determined by a draw from a binomial distribution:

$$r_m \sim \text{Binomial}(r_d, \frac{\mathbb{E}(r_m)}{\mathbb{E}(r_m) + \mathbb{E}(r_w)}$$

**Parameter values**

The values for the grid parameters were taken to cover the range of values seen in the 2,778 samples from PCAWG and beyond.

The coverage of the simulations was fixed to the average coverage in PCAWG, $coverage = 48.46621$.

Six purity clusters were selected using *kmeans* on the PCAWG purity values (Fig. 27). Five clusters for the number of clonal mutations were defined by *kmeans*, one of which was fixed at $N_{SNV} = 10^5$, to better cover hypermutators, i.e. an important corner case in real data. Similarly, five clusters for the fraction of clonal mutations were defined using *kmeans*, one of which was fixed at $f_{clonal} = 0.995$, to cover the typical corner case of melanomas, i.e. high $N_{SNV}$ and high $f_{clonal}$. Finally, four clusters of number of subclones were selected, corresponding to $N_{subclones} = \{0,1,2,3^+\}$, where the number of subclones in the $3^+$ categories was drawn uniformly from $\{3,4,5,6,7\}$.

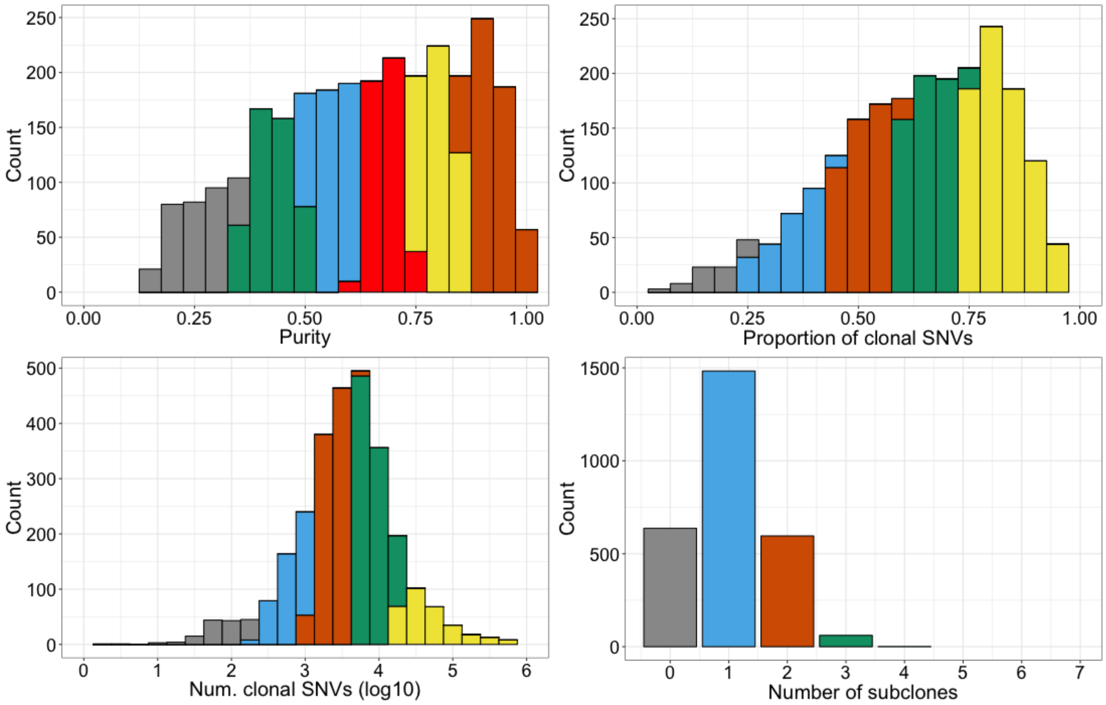


*Fig. 27 The simulated data set was created as a grid with four axis. Each axis represents a type of measurement that can be obtained from real data. This figure shows the histogram of these four measurements from the PCAWG data and the colors represent bins along each grid axis. A simulated tumor falls somewhere on the grid, which amounts to a combination of 4 bins (one on each axis). The parameters for this sample are then generated by sampling a single value from each of the 4 bins.*

This led to $N = 6 \times 5 \times 5 \times 4 = 600$ combinations of grid parameter values.

#### Limit overfitting of the grid - adding noise

To limit the ability for individual methods to infer the grid design and thus tune their output to better (over-)fit the grid, we added noise in the input simulations at multiple levels:

##### i. grid coverage

From 600 combinations of parameters for the grid, i.e. the *wholebatch*, we randomly selected 700 combinations with replacement in two passes: a first selection of 600 combinations with replacement from *wholebatch* yielded *batch1*; a second selection of 100 combinations with replacement from $\{subbatch \mid subbatch \in wholebatch, subbatch \notin batch1\}$ yielded *batch2*. The final batch was the union of *batch1* and *batch2*. This two-pass strategy was selected as to maximize the coverage of the grid values (Fig. 28).

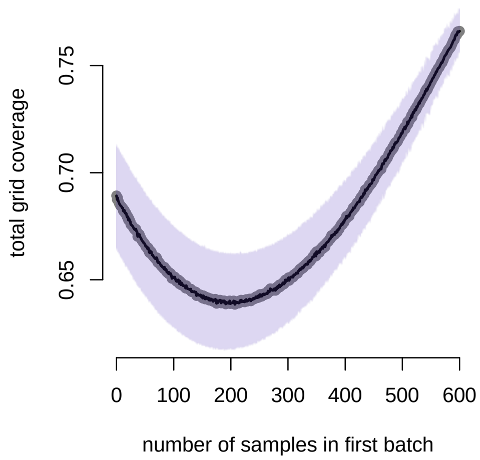


*Fig. 28 Number of samples N1 out of the 600 samples to select from with replacement in the first batch batch1. The second batch batch2 of 700-N1 samples is selected with replacement from the {wholebatch-batch1} to get to a total of 700 simulations. Y- axis: proportion of grid coverage.*

Using this two-pass strategy, we randomly selected 700 combinations, covering 462 out of the 600 combinations, i.e. 77% of the grid.

##### ii. input values

Input values were randomly selected from the clusters of PCAWG values defined by the *kmeans* clustering for purity, fraction clonal and number of clonal mutations.

##### iii. purity input values

The selected purity values *purity*$_{in}$ were used to generate the simulations. However, the purity values given to the individual methods as input were defined as

$$purity_{out} = purity_{in} + purity_{error}$$

where $purity_{error} \sim U(-mad_{purity}, mad_{purity})$. The median absolute deviations (mad) on the purity values were taken from the consensus purity values in PCAWG. This explains why some of the purity values were slightly greater than 1. However, we verified that no purity values were negative.

**Copy number profiles**

The copy number profiles were the consensus copy number profiles taken from the PCAWG sample from which the purity value was used. We only selected samples for which the fraction of genome aberrated $f_{aberrated} > 0.1$, as measured from the consensus copy number profiles.

To assess the impact of copy number on the reconstructions, a random set of 300 samples from the 700 were simulated with fully diploid genomes, leading to a total of 1,000 simulations. Their purity was adapted to maintain the same average nrpcc, i.e.

$$purity_{diploid} = \frac{2}{ploidy_{non-diploid}} purity_{non-diploid}$$

**Defining the cluster sizes and locations**

The CCFs of the subclones were drawn from a uniform distribution $CCF_{subclone} \sim U(0.1, 0.9)$. The sizes of the clusters were defined using a stick-breaking procedure. Starting with a number of subclonal mutations $N_{subclonal} = \frac{N_{clonal} - f_{clonal} N_{clonal}}{f_{clonal}}$, we iteratively selected a proportion $p_{N_{clonal}} \sim U(0.1, 0.9)$ of the remaining SNVs and randomly assigned the obtained numbers of SNVs to the subclones.

##### Winner's curse correction

The simulator removes mutations for which $alt_{count} < min_{count}$, with $min_{count}$=3, which is at the origin of a *Winner's curse* effect.
To approach the target $N_{SNV}$ output by the simulator after this filtering, we compute $N_{input}$, such that over 10 simulations the average $N_{output} \in \left[N_{SNV} - \frac{N_{SNV}}{20}, N_{SNV} + \frac{N_{SNV}}{20}\right]$. We find $N_{input}$ using a dichotomic search.

##### Filtering step

We further removed simulations for which the total number of SNVs $N_{tot} < \min_{i \in PCAWG} N_i$ and for which the generation lasted longer than two days on our cluster nodes - due to the high number of SNVs. This led to a final set of 965 simulations.

### 3. Results of reconstructing the subclonal architecture of tumors

#### 3.1 Power analysis

Kerstin Haase, Peter Van Loo

When determining the number of subclones in each sample, we are limited by multiple factors influencing the sensitivity of our subclonal reconstruction methods. Given that the PCAWG mutation-calling pipeline requires at least three variant reads to be able to detect a mutation, ploidy and purity of a sample as well as sequencing coverage impact the probability of a mutation being identified.
To account for varying purity, ploidy and sequencing depth in the analyzed samples, we calculated the number of reads per clonal copy (nrpcc) to uniformly quantify the power to detect subclonal mutation clusters:

$$\text{nrpcc} = \frac{\rho \cdot \text{cov}}{\rho\psi_t + (1 - \rho)\psi_n}$$

where $\rho$ is the determined purity of the sample and $\psi_t$ and $\psi_n$ denote the ploidy of the matching tumor and normal sample, respectively. As we assume all germline samples to be diploid, $\psi_n$ is set to two by default. We verified that the nrpcc is a strong factor influencing the number of identified subclones in a sample, whereas the total number of mutations identified does not impact the reconstruction (**Supp. Fig. 2a**).

Given that a mutation can be detected when it is supported by at least three variant reads, we are theoretically able, in a sample with nrpcc of at least 10, to detect mutations in 30% of tumor cells or more. This consideration is supported by the reconstructions (**Supp. Fig. 2b**). As a result, we decided to limit the downstream analyzes to samples with a minimum nrpcc of 10 to exclude tumors that would appear completely clonal simply because of a lack of power to detect subclonal mutations.

#### 3.2 Results by individual methods are consistent with consensus results

Kerstin Haase, David Wedge, Peter Van Loo

After the results of the 11 subclonal reconstruction methods were combined by the consensus approach (section 2.2), we wanted to confirm that the final subclonal architecture does indeed reflect the individual solutions. For this purpose, we compared the fraction of subclonal mutations per cancer type between the 11 input methods and the consensus reconstruction. Overall, the distributions were very similar, and the ranking of cancer types by median fraction of clonal mutations was mostly conserved (Fig. 29).

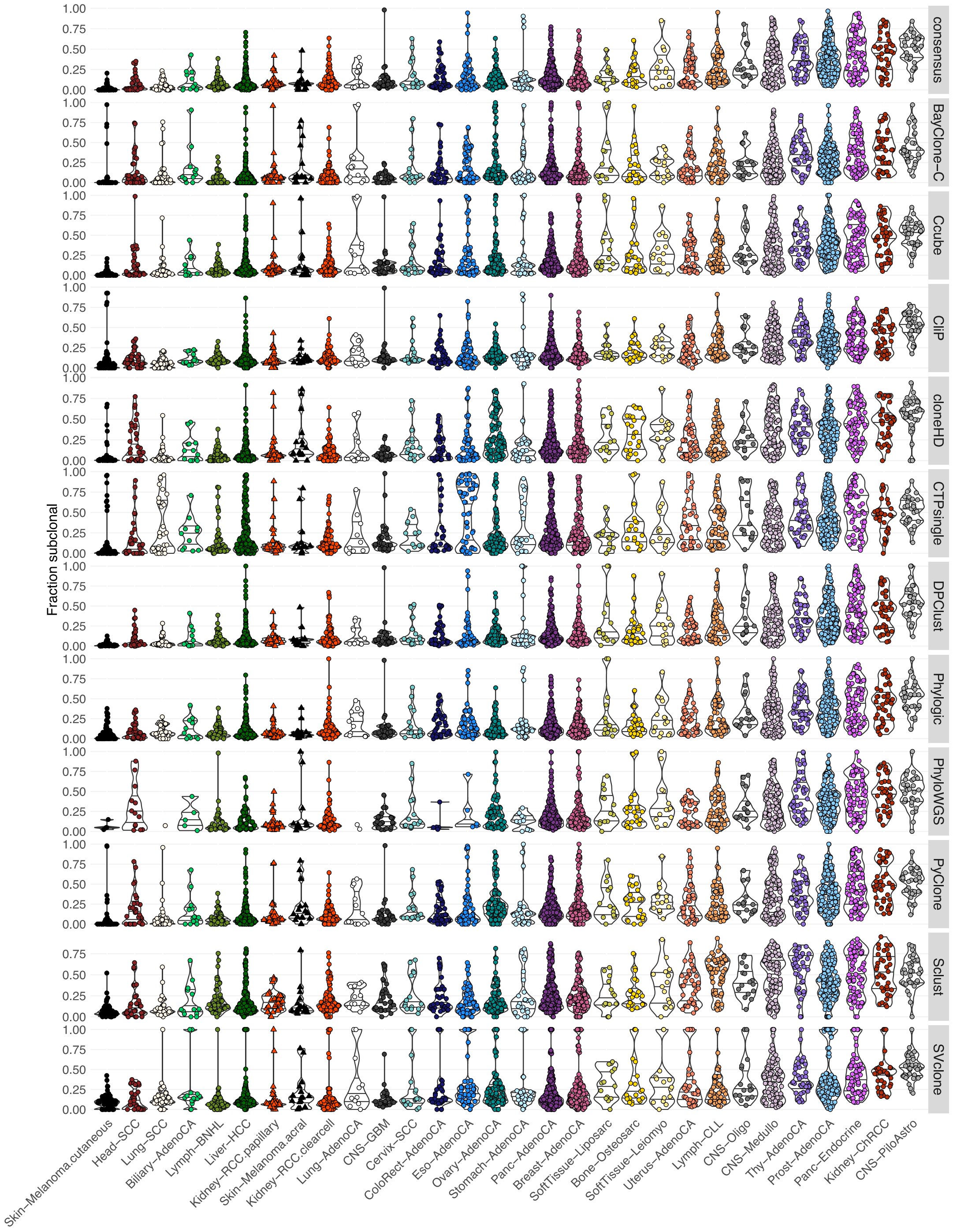


*Fig. 29. Fraction of subclonal mutations for all eleven subclonal reconstruction methods and the consensus is shown for 1,705 samples with sufficient power to detect subclones at CCF > 30%. Samples have been limited to those with less than 2% tumor contamination in the matched normal sample and no activity of any of the identified artefact signatures (Alexandrov et al., 2020). Only representative samples (The ICGC/TCGA Pan-Cancer Analysis of Whole Genomes Consortium, 2020) from multi-sample cases are shown. Cancer types are ordered by median fraction of subclonal mutations in the consensus reconstruction. The distributions of the fraction of subclonal mutations per cancer type, as determined by the individual methods, are very similar to the reported consensus architecture.*

#### 3.3 Correlation between fraction of subclonal mutations and mutation burden

Kerstin Haase, Maxime Tarabichi, Peter Van Loo

As the main **Fig. 3** seemed to show a negative correlation between overall mutation burden and the fraction of subclonal mutations over the whole cohort, we investigated the Pearson's correlation within each cancer type. After correction for multiple testing using Benjamini-Hochberg (Benjamini and Hochberg, 1995) only glioblastoma, medulloblastoma, and colorectal adenocarcinoma show a significant correlation (FDR<0.05; Table 1), demonstrating that there is no systematic technical bias causing the amount of subclonal mutations to be dependent on mutation burden.

*Table 1 Table showing the Pearson's correlation p-values of fraction of subclonal mutations and mutation burden per cancer type. Adjustment was performed according to Benjamini and Hochberg (Benjamini and Hochberg, 1995).*

| Cancer type | p-value | Adj. p-value |
|---|---|---|
| Biliary-AdenoCA | 0.446 | 0.668 |
| Bone-Osteosarc | 0.171 | 0.376 |
| Breast-AdenoCA | 0.126 | 0.342 |
| Cervix-SCC | 0.214 | 0.427 |
| CNS-GBM | 0.000 | 0.000 |
| CNS-Medullo | 0.002 | 0.018 |
| CNS-Oligo | 0.012 | 0.089 |
| CNS-PiloAstro | 0.229 | 0.429 |
| ColoRect-AdenoCA | 0.002 | 0.018 |
| Eso-AdenoCA | 0.173 | 0.376 |
| Head-SCC | 0.115 | 0.342 |
| Kidney-ChRCC | 0.175 | 0.376 |
| Kidney-RCC.clearcell | 0.015 | 0.089 |
| Kidney-RCC.papillary | 0.828 | 0.928 |
| Liver-HCC | 0.080 | 0.266 |
| Lung-AdenoCA | 0.646 | 0.896 |
| Lung-SCC | 0.342 | 0.570 |
| Lymph-BNHL | 0.686 | 0.896 |
| Lymph-CLL | 0.069 | 0.260 |
| Ovary-AdenoCA | 0.784 | 0.928 |
| Panc-AdenoCA | 0.029 | 0.146 |
| Panc-Endocrine | 0.871 | 0.933 |
| Prost-AdenoCA | 0.716 | 0.896 |
| Skin-Melanoma.acral | 0.970 | 0.970 |
| Skin-Melanoma.cutaneous | 0.969 | 0.970 |
| SoftTissue-Leiomyo | 0.042 | 0.180 |
| SoftTissue-Liposarc | 0.419 | 0.661 |
| Stomach-AdenoCA | 0.261 | 0.461 |
| Thy-AdenoCA | 0.708 | 0.896 |
| Uterus-AdenoCA | 0.835 | 0.928 |

### 4. Selection and driver genes

#### 4.1 Selection and dN/dS

Maxime Tarabichi, Iñigo Martincorena, Peter Van Loo

dN/dS is the ratio between the rates of nonsynonymous and synonymous substitutions. This metric has been extensively used for the detection of selection acting on protein-coding genes since its conception in the 1980s and has a long history in the field of molecular evolution. More recently, dN/dS has also been used to study selection in cancer genomes. In this study, we use a Poisson implementation of dN/dS, initially developed in (Greenman et al., 2006)), and further developed in (Martincorena et al., 2015)).

This implementation is feasible in the context of cancer genomics owing to the very low density of mutations per genome (typically < $1x10^{-5}$ mutations/bp), and it allows the use of complex trinucleotide substitution models that avoid biases due to context-dependent mutational processes (Martincorena et al., 2017). Point substitutions are classified according to two criteria: (1) the 192 possible strand-specific trinucleotide changes $i$ considering one base up- and downstream of the mutant base, and (2) their functional impact (as synonymous $s$, missense $m$, nonsense $n$ and essential splice site $e$ substitutions). For example, the number of C>T missense mutations in an A[C]G context observed in a cohort of samples can be modelled as:

$$n_{ACG>ATG,m} \sim Poisson(\lambda = r_{ACG>ATG} L_{ACG>ATG,m} \omega_m)$$

Where, $r_{ACG>ATG}$ is the rate of ACG>ATG mutations per ACG site, $L_{ACG>ATG,m}$ is the number of ACG trinucleotides in the sequences analyzed whose change for an ATG would lead to a missense amino acid change, and $\omega_m$ is the dN/dS ratio for missense mutations. Given that there are 192 by 4 possible nucleotide changes based on this classification, there are 768 equations like the one above. Maximum-likelihood estimates for all parameters as well as confidence intervals, including for all 192 parameters and the 3 possible ω (*i.e.* dN/dS) parameters are estimated using Poisson regression.

Using this framework, dN/dS ratios can be calculated for different groups of mutations, such as clonal and subclonal mutations in known cancer genes, which yields insights about the density of driver mutations in each group of mutations. With the dndscv (Martincorena et al., 2015) R package version 0.0.0.9 dN/dS analysis was run on the clonal and subclonal mutations as described above using the 566 cancer gene census from COSMIC v84 in *Tier1* (i.e. high confidence cancer genes) that were included in the hg19 *refCDS* gene list from dndscv. The maximum number of coding mutation per sample was set to 500. This revealed widespread positive selection in both clonal and subclonal mutations of both neutral and non-neutral tumor groups (main **Fig. 3d**).

#### 4.2 Gene set analysis of subclonally mutated genes

Ignaty Leshchiner, Dimitri Livitz, Gad Getz

The top 30 subclonal driver genes were subjected to gene set analysis using *GO_molecular_function* gene sets from MSigDB (Subramanian et al., 2005). We derived p-values from the hypergeometric distribution for (k-1, K, N - K, n), where k is the number of genes in the intersection of the query set with a set from MSigDB; K is the number of genes in the set from MSigDB; N is the total number of genes; n is the number of genes in the query set. We obtained the false discovery rates by adjusting the hypergeometric p-values for multiple-hypothesis testing according to Benjamini-Hochberg. We found the following gene sets to be significantly enriched (Table 2).

*Table 2 Table of the 10 top significant gene sets (q<0.1) in the subclonally mutated genes. The columns in order give: the gene set name, the number of genes in the gene set, the p-value of the hypergeometric test, the FDR q-value.*

| Gene Set Name | # Genes | p-value | FDR q-value |
|---|---|---|---|
| GO_MACROMOLECULAR_COMPLEX_BINDING | 17 | 7.49E-17 | 3.89E-14 |
| GO_TRANSCRIPTION_FACTOR_BINDING | 13 | 8.64E-17 | 3.89E-14 |
| GO_CHROMATIN_BINDING | 11 | 2.34E-14 | 7.03E-12 |
| GO_ENZYME_BINDING | 16 | 5.79E-14 | 1.30E-11 |
| GO_REGULATORY_REGION_NUCLEIC_ACID_BINDING | 12 | 7.90E-13 | 1.42E-10 |
| GO_RNA_POLYMERASE_II_TRANSCRIPTION_FACTOR_BINDING | 7 | 1.96E-12 | 2.94E-10 |
| GO_PROTEIN_COMPLEX_BINDING | 11 | 8.81E-11 | 1.13E-08 |
| GO_DOUBLE_STRANDED_DNA_BINDING | 10 | 2.62E-10 | 2.95E-08 |
| GO_TRANSCRIPTION_FACTOR_ACTIVITY_PROTEIN_BINDING | 9 | 5.99E-10 | 5.99E-08 |
| GO_RECEPTOR_BINDING | 12 | 7.06E-10 | 6.36E-08 |

### 5. Evidence of additional ITH

#### 5.1 Analysis of phaseable mutation pairs

Jonas Demeulemeester, Amit Deshwar, Stefan Dentro, Ignaty Leschchiner, Maxime Tarabichi, Peter Van Loo, Quaid D. Morris

Phased SNV pairs can provide direct evidence of whether two SNVs co-occur in the same cell, and thus are in collinear subclonal lineages, or whether they are in separate cells, i.e., are mutually exclusive. Specifically, if read pairs cover the loci of two nearby SNVs, each such read pair is a pairwise haplotype and the pattern of these haplotypes can be examined to establish the ancestral relationships between the subclonal lineages containing the SNVs. Assuming that the haplotypes are accurate and controlling for violations of the infinite sites assumption (see 5.1.1), if two SNVs are in-*cis* (they appear on the same haplotype), they must be collinear. Furthermore, if two SNVs are in-*cis,* and one of the haplotypes contains only one of the SNVs, then that SNV occurred prior to the other along the lineage. In contrast, if two SNVs are in-*trans*, and they appear in a haploid region (e.g. on chromosome X in a male patient, or in a clonal copy number 1+0 region), then the two SNVs are mutually exclusive and must have occurred on distinct branches of the phylogenetic tree.

We obtained phasing information for all SNV pairs that are within 700bp. Upon first inspection, we noted that a lot of putative pairs harbouring both somatic SNVs were located in low-complexity regions of the genome or contained multiple mismatches in the read (a sign of alignment artefact). We therefore applied the following stringent filtering, retaining only pairs with mapping quality $\geq 20$, mismatch bases quality $\geq 25$, no hard or soft-clipped bases, that were properly paired, were not flagged as duplicates and did not have a failed vendor quality control flag. Furthermore, we removed read pairs with indels and those that had $\geq 2$ mismatches in a single read or $\geq 3$ in the whole pair (if both phased SNVs were spanned by different reads in the pair). To further reduce biases we removed all pairs in regions known to undergo somatic hypermutation (T-cell receptor $\alpha, \beta, \gamma, \delta$ and the immunoglobulin heavy and light chain loci), and those flagged up as being part of kataegis events (The ICGC/TCGA Pan-Cancer Analysis of Whole Genomes Consortium, 2020). In addition, pairs in the pseudo-autosomal regions of X and Y in males are excluded, as well as all pairs on consensus copy number segments with confidence level below d (see 1.3 Consensus copy number approach). Lastly, pairs on copy number segments which show evidence for a shifted CCF distribution of the SNVs on that segment, compared to the SNVs on all other confidence level a–d segments are also excluded (one-sided Mann–Whitney *U* test, Benjamini-Hochberg corrected $q \leq 0.01$).

Having obtained high quality phasing information, we identify collinear phylogenetic relationships by looking for SNV pairs that meet the following criteria:

- $\geq 2$ read pairs supporting both mutant alleles (Mut-Mut)
- $\geq 2$ read pairs supporting only one of the two mutant alleles (either Mut-WT or WT-Mut, but not both)
- the SNVs fall within a single segment with major allele copy number equal to 1

- the lower bound estimate on the number of infinite sites violations < 1 or the copy number of the minor allele is equal to 0 and Battenberg reports no evidence for subclonality (see 1.2.3 Battenberg).

Branching phylogenetic relationships are revealed by looking for SNV pairs that meet the following criteria:

- no read pairs supporting both mutant alleles (Mut-Mut)
- $\geq 2$ read pairs each in support of the two tumor haplotypes (Mut-WT and WT-Mut)
- the SNVs fall within a single haploid segment (copy number 1+0) and Battenberg reports no evidence for subclonality (see 1.2.3 Battenberg).

A total of 149,452 SNV pairs samples pass the quality filters above, are in the appropriate copy number context and have at least two reads supporting each haplotype. Of these, 140,259 are fully phased (93.8%), 8,140 are in-*trans* in a diploid region (5.5%), 774 and 277 support collinear and branching evolution, respectively (0.52% and 0.19%), and only 2 pairs (0.0013%) have inconsistent phasing.

The identification of collinear or branching evolution from phased SNV pairs in powered samples may be considered largely independent of the consensus subclonal reconstruction. While they both require copy number calls, this is only to identify informative regions for phasing. One can therefore use phasing results to assess the performance of the consensus subclonal reconstruction. Indeed, tumors identified to contain higher numbers of consensus subclones are enriched for collinear and branching pairs ($\chi^2$-test *p*-value = $3.9\text{x}10^{-15}$)

The frequency of branching versus linear evolution can be compared directly by subsetting linear pairs to only those that fall in regions where branching calls can also be made (i.e clonal haploid regions, see above). Unlike branching pairs, linear pairs can additionally encode clone-subclone relations. We therefore subset all linear and branching pairs to those where both SNVs have been assigned to subclones in the consensus. Considering the initially greater number of linear pairs and the probabilistic cluster assignments, we still expect the number of subclone-subclone reporting linear pairs to be inflated. As a result, the odds of branching vs linear subclonal evolution estimated here reflect a lower bound on the underlying distribution of subclonal evolutionary pathways.

#### 5.1.1 Infinite Sites Assumption

Querying phylogenies through phasing analysis of proximal SNVs requires the infinite sites assumption to hold. Specifically, phasing to mutations that have occurred in parallel on both copies in a diploid region may be misinterpreted as evidence for a linear phylogeny. We opted for a simulation approach to establish a lower bound estimate of the expected number of such violations in any given sample.

In a single simulation, the observed SNVs in a sample are permuted across the callable regions of a diploid genome, retaining their trinucleotide context. Overlapping trinucleotides are considered to be independent from one another and any position on an allele can be hit only once (i.e. no back or forward mutation). For each sample, 1000 simulations are run, and the average number of these parallel mutations (double hits) counted. While retaining the trinucleotide context of mutations, these simulations assume a uniform distribution of mutations across the callable genome. Results from this model are hence regarded as a lower bound on the number of expected violations, since any deviation from uniform will only increase the rate of infinite sites violations.

##### 5.1.2 Analysis of TRACERx 100 phylogenetic trees

To compare the odds of branching versus linear evolution obtained from phased subclonal SNV pairs in PCAWG to what is expected, we analyzed the detailed phylogenetic trees obtained through multiregion sequencing on the TRACERx 100 non-small-cell lung cancer cohort (Jamal-Hanjani et al., 2017). From each tree in the cohort, two subclones were randomly sampled and their relation on the phylogenetic tree (sibling or parent-child) was assessed. This analysis was performed 1,000 times. Every iteration, the odds were computed as the number of times sibling subclones were sampled divided by the number of times parent-child subclones were sampled. The median and 95% confidence interval are reported.

#### 5.2 Tracking signature activities across cancer timelines

Yulia Rubanova, Quaid D. Morris

Mutations accumulate in cancer cells due to external (e.g., smoking, exposure to UV light) or internal (e.g. copy errors, failure of DNA damage repair) causes. Each source can be characterized by a probability distribution over 96 mutation types, known as a mutational *signature* (Alexandrov et al., 2013). The rate at which new mutations are generated by a certain process can vary over time and is known as the *signature activity*. Note that multiple mutational sources can be active in the same tumor.

We used a probabilistic method called TrackSig to fit the evolutionary trajectories of signature activities in individual tumors. This method is described in detail in a separate manuscript (Rubanova et al., 2020). Here we briefly describe (i) the source of the mutation signatures, (ii) the major steps in trajectory reconstruction by Trackature, (iii) how significance of signature activity change was computed, (iv) how the estimates of coincidence between signature activity change and clonal / subclonal boundaries (and subclonal / subclonal boundaries) were computed.

**(i) Source of mutation signatures**

We used the Beta2 release of the mutation signatures by PCAWG working group 7 (Alexandrov et al., 2020). For each sample, we only considered mutation signatures marked as active.

**(ii) Trackature fitting of evolutionary trajectories of signature activity**

*Assessing pseudo-time ordering of mutations*

First, we order the mutations based on their approximate relative order of occurrence in the tumor. Generally, if the mutation occurred early in the tumor development, it will be present at high cellular prevalence (CP), whereas late mutations have lower CP. We exploit this observation to infer the pseudo-time of mutation occurrence. We take the variant allele frequency (VAF) of the mutations as computed from the read counts. To eliminate the effects of quantization noise we then re-sample the VAF value from the posterior distribution given the variant and reference read counts. VAF estimates are converted into "average mutation copy number/cell"-space by multiplying the CP by the copy number. This space yields a “pseudo-time” ordering of mutations, but how this relates to real time is unknown. For convenience, we divide the time-ordered mutations into equal bins of 100 mutations. The resulting bins, each corresponding to one time point, constitute a pseudo-timeline of tumor development, which we leverage to map the signature trajectories. Timelines are constructed separately for each sample.

This method does not consider branching tumor evolution. If two subclones have a similar cellular prevalence, mutations from those subclones are likely to fall into the same time points. Note that this issue may be tackled by estimating the trajectories separately for each subclone.

*Activity estimation*

Next, we estimate the signature proportions. At each time point, the SNVs are classified into 96 types based on their trinucleotide context. Then we fit a mixture of multinomials, where each component multinomial describes one of the known active signature distributions over the 96 types. Derived mixture component coefficients correspond to the signature activities and sum to 1 for each time point. The activity value reflects the fraction of the mutations generated by the associated mutational process. For ease of interpretation, activities are given as percentage. By estimating the activities at each time point, a set of trajectories is obtained. Each trajectory represents the activity of the corresponding mutational process during the evolutionary history of the tumor sample.

*Detecting change points*

Finally, we seek to find the time points at which signature activities change substantially, as these represent changes in the activity of the different mutational processes. If certain mutational processes have become more or less active in a subclone when compared to the clone (e.g. inactivation of double strand break repair in the subclone), its emergence may be accompanied by a drastic shift in the trajectories. Otherwise, we expect the trajectories to remain relatively constant.

To detect the signature change points, we use Pruned Exact Linear Time (PELT) algorithm (Killick et al., 2012)**.** PELT is based on dynamic programming. The algorithm iterates over the time points and re-computes the multinomial mixtures for all the mutations at both sides from the potential change point, treating each side as a single bin. We estimate the likelihood of the data split according to the potential change point. A point that maximizes the likelihood is considered a new change point. The Bayesian Information Criterion (BIC) is used to determine the optimal number of change points: introducing any new change point has to be accompanied by a sufficient increase in likelihood.

Note that change points are inferred without using any information from the subclonal reconstruction, allowing the method to make independent conclusions about signature behavior and its relation to subclones.

**(iii) Evaluating uncertainty**

To evaluate the uncertainty in activities, we resample the set of mutations 30 times and re-compute activity estimates. From our observations, the standard error across these bootstrapped activities does not exceed 5% absolute activity (with mean standard error 2%). Therefore, we omit change points with changes less than 5%. To evaluate overall signature changes in the tumors, we compute the maximal change of each signature as the difference between the maximum and minimum activity across pseudo-time. We consider that a tumor has a significant overall signature change if this difference exceeds 10%.

**(iv) Comparing change points to CCF clustering**

We investigate the relation between the signatures changes and CCF clustering. First, we determine where the CCF clustering boundaries map on our pseudo-timeline by dividing it according to the proportion of mutations per time point that belong to each CCF cluster. Distances in time are computed between those boundaries and our change points. To determine whether a subclonal boundary is supported by an activity change-point, we plotted the relative enrichment of change-points at a given offset from subclonal boundaries (Fig. 30). Using this figure, we deem a boundary to be supported if it has a change-point within 3 time points (tp).

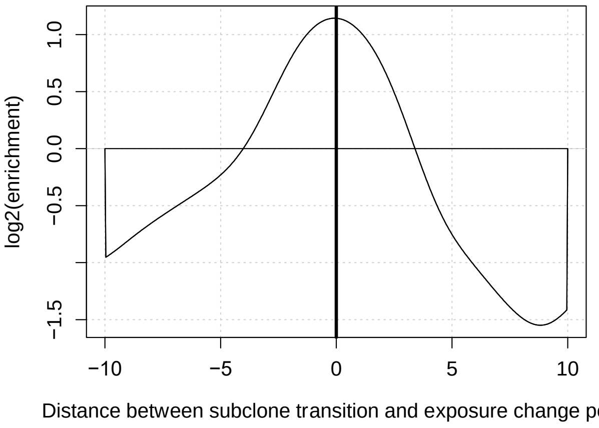


*Fig. 30 Changes in signature activities occur near subclone boundaries. Plot displays relative enrichment (y-axis) versus random control of activity change points at given time point offset (x-axis) from subclone boundaries. The smoothing window used when computing activity trajectories spans three time points, so sub clone boundaries between offsets -3 to 3 are deemed coincident with the activity change points.*

We compute the percentage of clonal/subclonal and subclonal/subclonal boundaries that are supported by signature change points. In total, 54.7% of clonal-subclonal and 57.3% of subclonal-subclonal boundaries in all tumors are supported by our change-points (red bars on **Fig. 6d** in the main text). These estimates are upper bounds on the number of supported boundaries, because some of the boundaries would be supported even if the change-points were randomly assigned to a time point.

To determine a lower bound on these percentages, we sample random points from the timeline and re-compute the distances between the boundaries and these random points. On average, 34.52% of clonal-subclonal and 34.49% of subclonal-subclonal boundaries are supported across 1000 random samples (green bars on **Fig. 6d** in the main text).

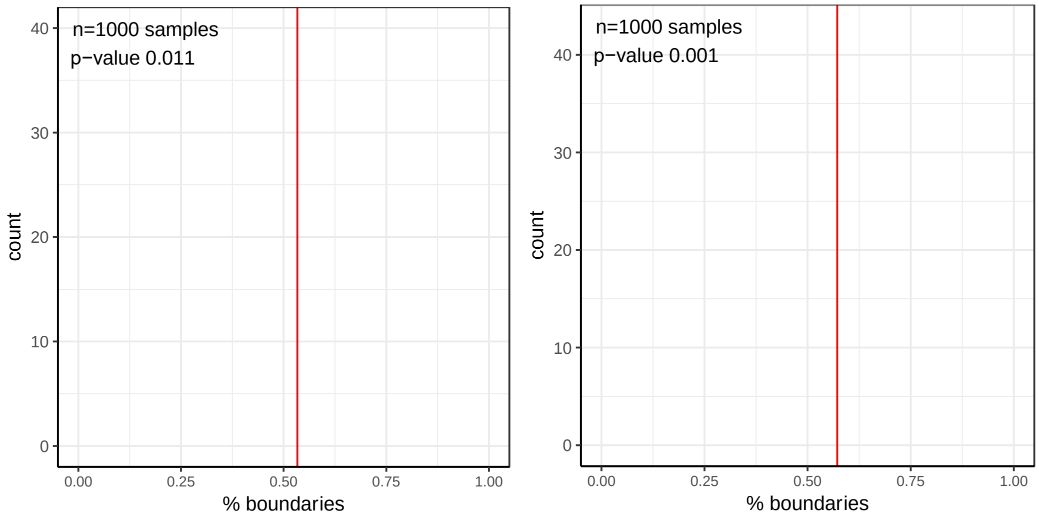


*Fig. 31 The distribution of the proportion of boundaries supported by randomly sampled points. The red line shows the proportion of boundaries supported by* ***real*** *change-points.*

To estimate the significance of boundary support, we sample random points 1000 times for each tumor and show the distribution of proportions of supported boundaries on Fig. 31. The empirical p-value is computed as the number of occurrences when then proportion of boundaries supported by random exceeds the then proportion of boundaries supported by real change points of signature activities.

### 6. Analysis of drivers in clinically targetable genes

Stefan Dentro, David Wedge

To assess the potential clinical relevance of subclonal architecture inference, we considered driver mutations (SNVs and indels) in genes and pathways for which drugs have been developed or are currently in development. This allows us to estimate frequency of cases in which the only currently actionable mutation is subclonal. A list of targetable drivers was provided by the PCAWG driver group (Rheinbay et al., 2020). In this analysis we excluded the relapse tumors and all metastases except for the melanomas. For multi-sample cases, we only considered the PCAWG provided preferred sample for each donor.

Our consensus subclonal architecture approach produces probabilistic cluster assignments for each mutation. The procedure also identifies a mutation cluster as clonal (CCF of 1), allowing us to establish the probability that a mutation is clonal or subclonal. The procedure for driver mutations or protein-altering mutations is as follows: for each sample, establish the probability that all such mutations are clonal, that they are subclonal, and that there is at least one pair where one is clonal and one is subclonal.

The probability $(p)$ of observing all $n$ such mutations as clonal is:

$$\prod_{i=1}^{n} p_{i,clonal}$$

The probability of $(p)$ of observing all $n$ mutations as subclonal is:

$$\prod_{i=1}^{n} p_{i,subclonal}$$

Then the probability of observing at least one pair of mutations where one is clonal and one is subclonal is:

$$1 - \left(\prod_{i=1}^{n} p_{i,clonal} + \prod_{i=1}^{n} p_{i,subclonal}\right)$$

The three probabilities were summed to create the three classes shown in the figures: *Clonal*, *Subclonal* and *Both*.

### 7. SV analysis and fusion clonality detection

#### 7.1 Clonality analysis of recurrent structural variants

Geoff Macintyre, Ruben Drews, Florian Markowetz

The PCAWG Structural Variation Working Group identified 52 genomic regions containing significantly recurrent breakpoints (SRBs) and significantly recurrently juxtaposed regions (SRJs) (Rheinbay et al., 2020). As SRJs were mostly observed in small numbers of patients across individual tumor types, we focused our analysis here on SRBs, which were observed in larger numbers of patients across multiple tumor types.

Regions of SRBs ranged in size from 451 kb up to 5,651 kb (mean: 739 kb; median: 501 kb). 20 of the regions were enriched for amplifications, 9 for deletions, 9 for fusion events, and 5 for neutral rearrangements. Another 9 regions were characterized by general genomic instability and were therefore labelled “fragile” (Rheinbay et al., 2020).
SVs in fragile sites were considered candidate passengers rather than candidate driver events as they most likely arose due to genomic instability. To test this, we compared the difference in means of the number of subclonal SVs observed in fragile and the non-fragile loci with a Wilcoxon rank sum test. With a p-value of 0.024, we confirmed an enrichment of subclonal SRBs in fragile loci, which supports the model that SRBs in fragile sites are indeed likely passenger events (Fig. 32). Hence, fragile SRB loci were excluded from our analysis.

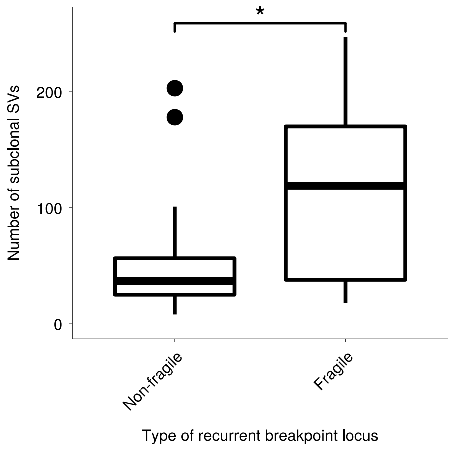


*Fig. 32 Comparison of the number of subclonal SVs observed in SRB loci annotated as fragile or non-fragile. * represents a significant difference ($p<0.05$) using the Wilcoxon rank sum test.*

The remaining 43 SRBs, covering 31,843 kb of the human genome, were used to determine which SVs were recurrent, and thus considered candidate driver SVs, and which were not recurrent, therefore considered candidate passenger SVs. For simplicity, SRBs were labelled by the name of the likely target gene lying within the

SRB. Note: this did not imply that this particular gene was mutated or altered in any way. A minimum overlap of at least 1bp with an SV was required. A SV having both breakpoints inside a SRB was counted as one hit. An SV with one breakpoint outside and one inside was also considered as one hit for this particular SRB and the breakpoint outside was not used.

From 2,658 samples originating from 37 cancer histotypes, we filtered samples using the following criteria: samples classified as greylisted or excluded (75 samples) were excluded from analysis. For patients with multiple specimens, the preferred sample was chosen as determined by the latest PCAWG metadata file (36 samples excluded). Samples with nrpcc (number of reads per chromosome copy) values of less than 10 were excluded as they showed a strong bias towards clonal SVs. Within each of the 36 tumor histotypes, samples having total SV counts more than two standard deviations from the mean were considered "mutator phenotypes" and excluded from further analysis (77 samples). Cancer types with fewer than 10 patients were also excluded (8). The final filtered set of SVs contained 129,219 SVs from 28 tumor histotypes originating from 1,517 patients.

**Testing for differences in SRB and passenger SV clonality**

We hypothesized that candidate driver SVs differed in their subclonal probability compared to passenger SVs. We used 1 - the probability of an SV being clonal as output by *mutationTimer* (Gerstung et al., 2020) for SVs preprocessed by *SVclone* (Cmero et al., 2020). As the distribution of clonal probabilities was assumed to be of non-parametric character and non-identical, we decided to infer the significance of the difference in subclonal probability by permutation testing. Since we discarded overly mutated samples, we assumed that the distribution of subclonal probabilities of SVs were independent of the samples.

When calculating medians, we weighted them linearly by a sample's individual nrpcc value by using the *weightedMedian* function of the R package *matrixStats.* This function allowed for the interpolation of results when ties arose.

The alternative hypothesis $\boldsymbol{H_a}$ was formed by taking the difference in weighted medians of the subclonal probabilities of candidate drivers and passengers. The permutation testing established a null distribution $\boldsymbol{H_0}$ of weighted differences between candidate driver and passenger probabilities by randomly assigning classification labels 10,000 times and taking the difference of the resulting medians. A p-value was generated by taking the proportion of the alternative hypothesis lying outside the generated null distribution. The resulting p-values were corrected for multiple hypothesis testing by multiplicity correction as proposed by Benjamini and Hochberg (Benjamini and Hochberg, 1995).

When plotting **Figure 6A,** the weighted interquartile ranges were used with the same linear weighing method applied as explained above. The IQRs were used as to represent the probabilities more closely to their density functions than the 95% confidence intervals as they tend to be very large due to the inherent noise of SV VAFs determined using short-read sequencing data (Cmero et al., 2020).

**Testing for clonal and subclonal SRB enrichment**

We used a GSEA-like test to see if any of the 43 SRB loci were enriched for clonal or subclonal SVs. For each SV, we transformed the subclonal probability, scaling by a weight vector representing the “detection power” of the sample in which each SV was observed. The weight vector was defined as the number of reads per chromosome copy for the sample, divided by the total across the dataset. All SVs were then ranked by their scaled subclonal probability. Using the fgsea package in R (Sergushichev, 2016), we computed an enrichment score for hits across the ranking for each SRB locus. The significance of the observed enrichment score was computed using a GSEA-like permutation test. p-values were corrected for multiple testing correction using the Benjamini and Hochberg method (Benjamini and Hochberg, 1995). Significantly enriched loci and the proportion of tumor types contributing to the enrichment appear in **Supplementary Figure 5**.

### 7.2 Fusion clonality analysis

Ignaty Leshchiner, Daniel Rosebrock, Gad Getz

A list of known driver fusions was curated by subsetting to fusions which appeared in at least 3 cases across the Jackson Lab fusion data set (http://www.tumorfusions.org/), or which were recorded as drivers in the COSMIC Fusion data set (http://cancer.sanger.ac.uk/cosmic/fusion) with at least 5 previously identified patient samples. Non-driver fusions were chosen by subsetting to fusions not classified as in-frame with known fusion partners from above. SV breakpoints were genotyped with BreakPointer (Drier et al., 2013) and SVClone, where mutant allele count was estimated by aggregating both chimeric read and read pair support of the fused allele, while reference allele count was estimated by summing both breakpoint spanning reads and read pairs supporting the reference allele. The likelihood of a fusion originating from a clonal or a subclonal cell population was then estimated with correction for local copy number and purity, according to PhylogicNDT model described above.
